# PTP-MEG2 regulates quantal size and fusion pore opening through two distinct structural bases and substrates

**DOI:** 10.1101/822031

**Authors:** Yun-Fei Xu, Xu Chen, Zhao Yang, Peng Xiao, Chun-Hua Liu, Kang-Shuai Li, Xiao-Zhen Yang, Yi-Jing Wang, Zhong-Liang Zhu, Zhi-Gang Xu, Sheng Zhang, Chuan Wang, You-Chen Song, Wei-Dong Zhao, Chang-He Wang, Zhi-Liang Ji, Zhong-Yin Zhang, Min Cui, Jin-Peng Sun, Xiao Yu

## Abstract

Tyrosine phosphorylation of secretion machinery proteins is a crucial regulatory mechanism for exocytosis. However, the participation of protein tyrosine phosphatases (PTPs) in different exocytosis stages has not been defined. Here we demonstrated that PTP-MEG2 controls multiple steps of catecholamine secretion. Biochemical and crystallographic analyses revealed key residues that the interactions between govern the PTP-MEG2 and NSF-pY^83^ site, specify PTP-MEG2 substrate selectivity and modulate the fusion of catecholamine-containing vesicles. Unexpectedly, delineation of PTP-MEG2 mutants along with the NSF binding interface revealed that PTP-MEG2 controls the fusion pore opening through non-NSF dependent mechanisms. Utilizing bioinformatics search and biochemical and electrochemical screening approaches, we discovered that PTP-MEG2 regulates the opening and extension of the fusion pore by dephosphorylating the DYNAMIN2-pY^125^ and MUNC18-1-pY^145^ site. Further structural and biochemical analysis confirmed the interaction of PTP-MEG2 with MUNC18-1-pY^145^ or DYNAMIN2-pY^125^ through a distinct structural basis compared with that of the NSF-pY^83^ site. Our studies extended mechanistic insights in complex exocytosis processes.

**HIGHLIGHTS:** PTP-MEG2 regulates multiple steps of exocytosis.

A crystal structure of the PTP-MEG2/phosphor-NSF-pY^83^ segment was obtained.

Functional delineation of the PTP-MEG2/NSF interface led to the discovery of new PTP-MEG2 substrates.

PTP-MEG2 regulates fusion pore opening and extension through the DYNAMIN2-pY^125^ site and MUNC18-1 pY^145^ site.

The distinct structural basis of the recognition of substrates by PTP-MEG2 allows selective inhibitor design.

## INTRODUCTION

Secretion via vesicle exocytosis is a fundamental biological event involved in almost all physiological processes (Dittman and Ryan, 2019; Neher and Brose, 2018; Wu et al., 2014). The contents of secreted vesicles include neuronal transmitters, immune factors, and other hormones (Alvarez de Toledo et al., 1993; Magadmi et al., 2019; Sudhof, 2013). There are three major exocytosis pathways in secretory cells, namely, full-collapse fusion, kiss-and-run, and compound exocytosis, which possess different secretion rates and release amount (Sudhof, 2004). It has previously been reported that the phosphorylation of critical proteins at serine/threonine or tyrosine residues participates in stimulus–secretion coupling in certain important exocytosis processes, for example, the secretion of insulin from pancreatic β cells and the secretion of catecholamine from the adrenal medulla (Laidlaw et al., 2017; Ortsater et al., 2014; Seino et al., 2009). However, the exact roles of protein tyrosine phosphatases (PTPs) in the regulation of key hormone secretion procedures are not fully understood.

The 68-kDa PTP-MEG2, encoded by *ptpn9*, is a non-receptor classical PTP encompassing a unique N-terminal domain with homology to the human CRAL/TRIO domain and yeast Sec14p (Alonso et al., 2004; Cho et al., 2006; Gu et al., 1992; Huynh et al., 2004; Zhang et al., 2016; Zhang et al., 2012). The N-terminal Sec14p homology domain of PTP-MEG2 recognizes specific phospholipids in the membrane structure and is responsible for its specific subcellular location. In secretory immune cells, PTP-MEG2 has been suggested to regulate vesicle fusion via direct dephosphorylation of the pY^83^ site of N-ethylmaleimide-sensitive fusion protein (NSF) (Huynh et al., 2004). However, many key issues regarding PTP-MEG2-regulated cell secretion remain controversial or even unexplored. For example, it is uncertain whether PTP-MEG2 regulates vesicle exocytosis only within immune cells (Zhang et al., 2016) and only play insignificant roles in other hormone secretion processes. It remains elusive whether PTP-MEG2 regulates vesicle trafficking pathways other than the NSF-mediated vesicle fusion.

Functional characterization of PTP-MEG2 in vivo normally requires a knockout model; however, PTP-MEG2-deficient mice show neural tube and vascular defects, and the lack of PTP-MEG2 is embryonic lethal (Wang et al., 2005). Alternatively, a specific small-molecule inhibitor of PTP-MEG2 has fast response withno compensatory effects, enabling it to serve as a powerful tool to investigate PTP-MEG2 functions (Yu and Zhang, 2018; Zhang et al., 2012). Recently, we have developed a potent and selective PTP-MEG2 inhibitor, Compound 7, that has a Ki of 34 nM and shows at least 10-fold selectivity for PTP-MEG2 over more than 20 other PTPs (Zhang et al., 2012). The application of this selective PTP-MEG2 inhibitor in combination with electrochemical approaches enabled us to reveal that PTP-MEG2 regulates multiple steps of catecholamine secretion from the adrenal medulla by controlling the vesicle size, the release probabilities of individual vesicles and the initial opening of the fusion pore during exocytosis. Further crystallographic studies of the PTP-MEG2 protein in complex with the pY^83^-NSF fragment and enzymological kinetic studies captured the transient interaction between PTP-MEG2 and NSF and provided the structural basis for PTP-MEG2 substrate specificity. Interestingly, by delineating the substrate specificity of deficient PTP-MEG2 mutants in the study of catecholamine secretion from primary chromaffin cells, our results suggested that PTP-MEG2 regulates the initial opening of the fusion pore during exocytosis by regulating substrates other than the known NSF through a distinct structural basis. We therefore took advantage of this key knowledge and utilized bioinformatics analysis, GST pull-down screening, enzymological and electrochemical techniques to identify the potential key PTP-MEG2 substrates involved in fusion pore opening. These experiments led to the identification of several new PTP-MEG2 substrates in the adrenal medulla, of which DYNAMIN2-pY^125^ and MUNC18-1-pY^145^ are the crucial dephosphorylation sites of PTP-MEG2 in the regulation of initial pore opening and expansion. Further crystallographic analyses and functional assays with MUNC18-1 Y^145^ and DYNAMIN2-pY^125^ revealed the mechanism underlying the recognition of MUNC18-1 and DYNAMIN2 by PTP-MEG2 and how these PTP-MEG2-mediated dephosphorylation events regulate fusion pore dynamics.

## RESULTS

### Phosphatase activity of PTP-MEG2 is required for catecholamine secretion from adrenal glands

Endogenous PTP-MEG2 expression was readily detected in mouse adrenal gland and chromaffin cell line PC12 cells (Supplemental Fig. 1A-B). Because PTP-MEG2 knockout is embryonic lethal (Wang et al., 2005), we therefore applied our newly developed active-site directed and specific PTP-MEG2 inhibitor (Compound 7) to investigate the functional roles of PTP-MEG2 in catecholamine secretion from adrenal glands (Supplemental Fig. 1C). Compound 7 is a potent and cell-permeable PTP-MEG2 inhibitor, with a Ki value of 34 nM and extraordinary selectivity against other phosphatases (Zhang et al., 2012). The administration of either high concentrations of potassium chloride (70 mM) or 100 nM Angiotensin II (AngII) significantly increased the secretion of both epinephrine (EPI) and norepinephrine (NE) from the adrenal medulla as previously reported (Liu et al., 2017; Teschemacher and Seward, 2000), and this effect was specifically blocked by pre-incubation with Compound 7 (400 nM) for 1 hour (Fig. 1A-D). Notably, basal catecholamine secretion also decreased after pre-incubation with Compound 7 (Fig. 1A-D). However, the intracellular catecholamine contents did not change in response to Compound 7 incubation (Supplemental Fig. 1D-E). These results indicate that PTP-MEG2 plays an essential role in catecholamine secretion from the adrenal medulla.

**Figure 1.**
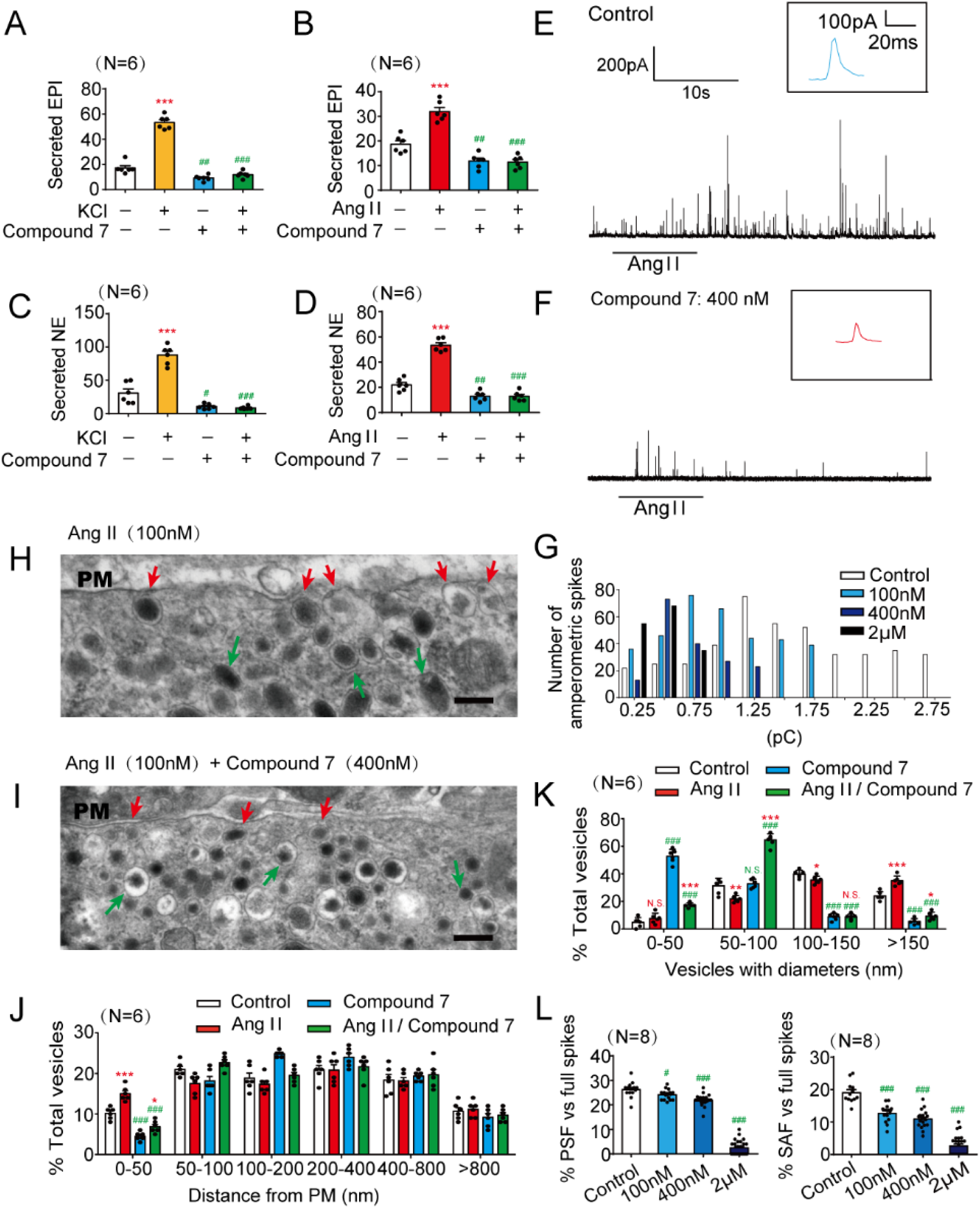
Inhibition of PTP-MEG2 reduced the spike amplitude and ‘foot’ probability of catecholamine secretion. (A-D). The epinephrine (A-B) or norepinephrine (C-D) secreted from the adrenal medulla was measured after stimulation with high KCl (70 mM) (A,C) or Angiotensin II (AngII, 100 nM) (B,D) for 1 min, with or without pre-incubation with a specific PTP-MEG2 inhibitor (400 nM) for 1 hours. Data were obtained from 6 independent experiments and displayed as the mean ± SEM. ***, P<0.001: cells stimulated with KCl or AngII compared with un-stimulated cells. #, P<0.05; ##, P<0.01; ###, P<0.001: cells pre-incubated with PTP-MEG2 inhibitors compared with control vehicles. All the data were analyzed using Student’s t-test. (E-F). Ang II (100 nM) induced amperometric spikes by primary mouse chromaffin cells after incubation with PTP-MEG2 inhibitor at different concentrations. (G). The distribution of the quantal size of AngII (100 nM)-induced amperometric spikes by primary chromaffin cells after incubation with different concentrations of PTP-MEG2 inhibitor. Histograms show the number of amperometric spikes of different quantal sizes. (H and I). Secretory vesicles of primary chromaffin cells were examined by transmission electron microscopy before or after 100 nM AngII stimulation, with or without pre-incubation with 400 nM Compound 7. Scale bars: H, I, 150 nm. Red arrows indicate morphologically docked LDCVs and green arrows stands for undocked LDCVs. PM represents plasma membrane. (J). Vesicle numbers according to different distances from the plasma membrane were calculated in the presence or absence of 100 nM AngII or 400 nM Compound 7. PM stands for plasma membrane. (K). The percentage of vesicles with different diameters was measured after 100 nM AngII stimulation or incubation with 400 nM Compound 7. Compound 7 significantly decreased vesicle size under AngII stimulation. (J)-(K) *, ** and *** indicate P<0.05, P<0.01 and P<0.001, respectively, between cells stimulated with AngII compared with un-stimulated cells. ### indicate P<0.001 between incubated with the control vehicles and cells incubated with Compound 7. N.S. means no significant difference. All the data were analyzed using Student’s t-test. (L). The percentage of pre-spike foot (PSF) (left panel) and stand-alone foot (SAF) (right panel) were calculated after incubation with the indicated concentrations of Compound 7. # indicates incubation with the indicated concentrations of Compound 7 compared with the subgroup with AngII and without Compound 7. # indicates P<0.05 and ### indicates P<0.001. All the data were analyzed using one-way ANOVA.

### PTP-MEG2 inhibition reduces the quantal size and the release probabilities of catecholamine secretion from individual vesicles

We used the carbon fiber electrode (CFE) to characterize the effects of PTP-MEG2 inhibition on the kinetics of catecholamine secretion from primary cultured chromaffin cells (Chen et al., 2005; Harada et al., 2015) (Fig. 1E-F and Supplemental Fig. 1F-G). The Ang II-induced catecholamine secretion was gradually attenuated by increasing the concentration of Compound 7 after pre-incubation with the primary chromaffin cells, from 20% at 100 nM Compound 7 to 80% at 2 μM Compound 7 (Fig. 1E-F and Supplemental Fig. 1 F-H). We then compared individual amperometric spikes of chromaffin cells pre-incubated with different concentrations of Compound 7 to determine the effect of PTP-MEG2 inhibition on quantal size (total amperometric spike charge) and vesicle release probabilities. The application of PTP-MEG2 inhibitor reduced quantal size, as indicated by statistical analysis of the quantal size distribution and averaged amperometric spike amplitude (Fig. 1G and Supplemental Fig. 1I). Specifically, the peak of the amperometric spikes decreased from 1.1 pC to 0.5 pC after incubation with 400 nM Compound 7 (Fig. 1G). The number of AngII-induced amperometric spikes also significantly decreased in the presence of Compound 7 (Supplemental Fig. 1J).

We then used transmission electron microscopy to examine the location of the large-dense-core vesicles (LDCVs) in the adrenal medulla after incubation with AngII and Compound 7 (Fig. 1H-I and Supplemental Fig. 2A-B). The intracellular distribution of the LDCVs was summarized by 50 nm bins according to their distance from the chromaffin plasma membrane (Fig. 1J). Vesicles with distance less than 50nm were generally considered to be in the docking stage. Here we found that the application of 400 nM Compound 7 significantly decreased the number of docking vesicles in contact with the plasma membrane (Fig. 1J). Moreover, Compound 7 significantly increased the number of LDCVs with a diameter less than 50 nm and decreased the number of LDCVs with a diameter greater than 150 nm (Fig. 1K), consistent with the statistical analysis of the total amperometric spike charge obtained by electrochemical measurements (Supplemental Fig. 1H-M). This change of vesicle size in the adrenal medulla agreed with the functional studies of PTP-MEG2 in immunocytes, which identified that vesicles were excessively larger if PTP-MEG2 was overexpressed (Huynh et al., 2004).

### PTP-MEG2 regulates the initial opening of the fusion pore

Generally, the presence of pre-spike foot (PSF) is a common phenomenon preceding large amperometric spikes which indicate catecholamine secretion in chromaffin cells, while stand-alone foot (SAF) is considered to represent “kiss-and-run” exocytosis (Chen et al., 2005). Both PSF and SAF are usually considered as indications of the initial opening of the fusion pore (Alvarez de Toledo et al., 1993; Zhou et al., 1996). In the present study, Compound 7 substantially decreased the PSF frequency from 25% to 3% and attenuated the SAF frequency from 18% to 4% in response to AngII stimulation (Fig. 1L). Consistent with PSF/SAF frequence, the inhibitor Compound 7 also decreased the average duration, amplitude and charge (Supplemental Fig.2C-J)

### The crystal structure of the PTP-MEG2/phospho-NSF complex reveals significant structural rearrangements in the WPD loop and β3-loop-β4

PTP-MEG2 is known to modulate interleukin-2 secretion in macrophages via dephosphorylation of NSF, a key regulator in vesicle fusion (Huynh et al., 2004). In response to stimulation with either high potassium chloride or AngII, the two stimulators for catecholamine secretion from the adrenal medulla, the tyrosine phosphorylation of NSF increased, indicating that NSF phosphorylation actively participates in catecholamine secretion (Fig. 2A and Supplemental Fig. 3A-B). Moreover, a significant portion of NSF co-localized with PTP-MEG2 in the adrenal medulla upon AngII stimulation, and the substrate-trapping mutant PTP-MEG2-D^470^A interacted with the tyrosine-phosphorylated NSF from the adrenal medulla treated with peroxide (Fig. 2B-D and Supplemental Fig. 3C-3E), suggesting that PTP-MEG2 regulates catecholamine secretion in chromaffin cells through direct dephosphorylation of NSF.

**Figure 2.**
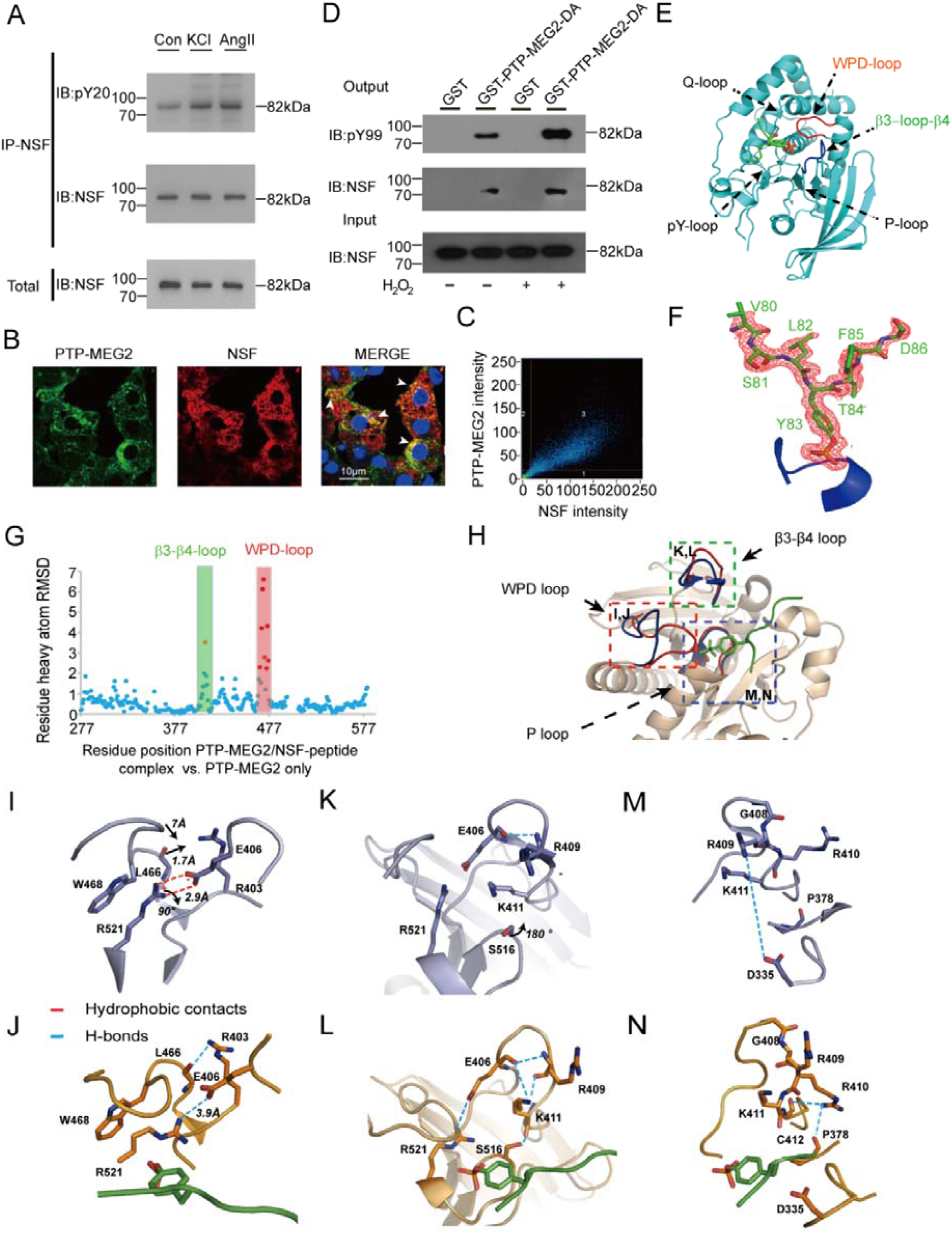
Interaction of PTP-MEG2 and tyrosine-phosphorylated NSF in the medulla and the crystal structure of the PTP-MEG2/phospho-NSF-segment complex. (A). NSF was phosphorylated after stimulation with AngII or KCl. Adrenal medulla cells were stimulated with 100 nM AngII or 70 mM KCl for 5 min and lysed. NSF was immunoprecipitated with a specific NSF antibody coated with Protein A/G beads. A pan-phospho-tyrosine antibody pY^20^ was used in Western blotting to detect the tyrosine-phosphorylated NSF in the adrenal medulla under different conditions. (B). NSF was co-localized with PTP-MEG2 in the adrenal medulla. After stimulation with 100 nM AngII, NSF and PTP-MEG2 in the adrenal medulla were visualized with immunofluorescence. White arrow stands for co-localization of the PTP-MEG2 and NSF. (C). Analysis of PTP-MEG2 and NSF fluorescence intensities by Pearson’s correlation analysis. The Pearson’s correlation coefficient was 0.7. (D). Phosphorylated NSF interacted with PTP-MEG2. PC12 cells were transfected with FLAG-NSF. After stimulation of the cells with 100nM AngII, the potential PTP-MEG2 substrate in cell lysates was pulled down with GST-PTP-MEG2-D^470^A or GST control. (E). The overall structure of PTP-MEG2-C^515^A/D^470^A in complex with the NSF-pY^83^ phospho-segment. (F). The 2Fo-Fc annealing omit map (contoured at 1.0σ) around the NSF-pY^83^ phospho-segment. (G). Plot of distance RMSDs of individual residues between the crystal structures of the PTP-MEG2/NSF-pY^83^ phospho-peptide complex (PDB:6KZQ) and the PTP-MEG2 native protein (PDB:2PA5). (H). Superposition of the PTP-MEG2/NSF-pY^83^ phospho-peptide complex structure (red) on the PTP-MEG2 native protein structure (PDB:2PA5, blue). The structural rearrangement of the WPD loop and β3-loop-β4 is highlighted. (I and J). The closure of the WPD loop and corresponding conformational changes in the inactive state (I) and the active state (J) of PTP-MEG2. The rotation of R^521^ leads to the movement of W^468^ and a corresponding 7 Å movement of the WPD loop. (K and L). The structural rearrangement of β3-loop-β4 of PTP-MEG2 in the active state (L) compared with the inactive state (K). The disruption of the salt bridge between E^406^ and R^521^ contributed to the new conformational state of the N-terminal of β3-loop-β4. (M and N). The conformational change of the P-loop of the inactive state (M) and the active state of PTP-MEG2 in response to NSF-pY^83^ segment binding (N). The disruption of the charge interaction between R^409^ and D^335^ resulted in the movement of the main chain from G^408^ to K^411^.

We therefore co-crystallized PTP-MEG2/phospho-NSF-E^79^-pY^83^-K^87^, and the structure was solved at 1.7 Å resolution (Table 1). The 2Fo-Fc electron density map allowed for the unambiguous assignment of the phospho-NSF-E^79^-pY^83^-K^87^ in the crystal structure (Fig. 2E-F). Importantly, the binding of phospho-NSF-E^79^-pY^83^-K^87^ induced substantial conformational changes in both the WPD loop and β3-loop-β4 compared to the crystal structure of PTP-MEG2 alone (Barr et al., 2009) (Fig. 2G-H). Specifically, the interaction of the phosphate group of pY^83^ of NSF with the guanine group of R^521^ of PTP-MEG2 induced rotation of approximately 90 degrees, which resulted in the movement of W^468^ and a traverse of 7 Å of the WPD loop for a closed state (Fig. 2I-J). Unique to the PTP-MEG2/substrate complex, the movement of L^466^ in the WPD loop by 1.7 Å enabled the formation of a new hydrogen bond between its main chain carbonyl group and the side chain of R^403^ (Fig. 2I-J). The disruption of the salt bridge between E^406^ and R^521^ also contributed to the new conformational state of the N-terminal of the β3-loop-β4 (Fig. 2K-L). The presence of the phosphate in the PTP-MEG2 active site C-terminal to β3-loop-β4 caused a 180-degree rotation of the side chain of S^516^, allowing its side chain oxygen to form a new hydrogen bond with the main chain carbonyl of group K^411^ (Fig. 2K-L). This structural rearrangement altered the side chain conformation of K^411^, which pointed to the solvent region and formed new polar interactions with E^406^ and the carbonyl group of R^409^ (Fig. 2K-L). Moreover, the presence of the phospho-NSF-E^79^-pY^83^-K^87^ peptide between R^409^ and D^335^ disrupted their charge interactions, enabling a movement of the main chain from G^408^ to K^411^, accompanied by a side chain movement of R^410^ and the formation of new polar interactions with the main chain of P^378^ and C^412^ (Fig. 2M-N). These structural rearrangements that occurred in the WPD loop and β3-loop-β4 enabled the accommodation of the phospho-substrate of PTP-MEG2 and may be important for its appropriate interactions with its physiological substrates/partners and subsequent activation.

**Table 1.** Crystallographic Data and Refinement Statistics.

| Data Collection | PTP-MEG2 · NSF<br>pY<sup>83</sup> Peptide | PTP-MEG2 · MUNC18-1<br>pY<sup>145</sup> Peptide |
| --- | --- | --- |
| Space group | p2<sub>1</sub>2<sub>1</sub>2 | p2<sub>1</sub>2<sub>1</sub>2 |
| Cell Dimensions |  |  |
| a (Å) | 71.447 | 71.048 |
| b (Å) | 83.708 | 84.074 |
| c (Å) | 48.615 | 48.753 |
| α (deg) | 90 | 90 |
| β (deg) | 90 | 90 |
| γ (deg) | 90 | 90 |
| Resolution (Å) | 80-1.70(1.76-1.70)<sup>a</sup> | 54.3-2.08(2.08-2.20)<sup>a</sup> |
| Unique observations | 32748(3167)<sup>a</sup> | 18019(1773)<sup>a</sup> |
| Completeness (%) | 99.6(98.3)<sup>a</sup> | 99.8(99.8)<sup>a</sup> |
| Redundancy | 3.8(3.8)<sup>a</sup> | 6.1(6.3)<sup>a</sup> |
| <I>/<σ> | 22.22(2.34)<sup>a</sup> | 6.5(2.3)<sup>a</sup> |
| R<sub>merge</sub> | 0.064(0.477)<sup>a</sup> | 0.191(0.927)<sup>a</sup> |
| Structure Refinement | 80-1.70 |  |
| Resolution (Å) | 1.70 | 2.08 |
| Reflections used for R<sub>work</sub>/R<sub>free</sub> | 31212/1536 | 17995/896 |
| R<sub>work</sub> /R<sub>free</sub><sup>b</sup> (%) | 0.1900/0.2173 | 0.2102/0.2304 |
| Average B-factor |  |  |
| Protein | 20.98 | 35.36 |
| Peptide | 31.525 | 53.25 |
| RMSD ideal bonds (Å) | 0.008 | 0.008 |
| RMSD ideal angles (deg) | 1.212 | 0.968 |
| RamachandranPlot (%) |  |  |
| Most favored | 93.73 | 93.56 |
| Allowed | 4.88 | 6.10 |
| Generously allowed | 1.39 | 0.34 |
| Disallowed | 0 | 0 |

<sup>a</sup> Each dataset was collected from a single crystal.
The values in parentheses correspond to the highest resolution shell.

### Structural basis of the PTP-MEG2-NSF interaction

The structural analysis identified critical residues for the phospho-substrate recognition by PTP-MEG2 (Fig. 3A). PTP-MEG2 Y^333^ forms extensive hydrophobic interactions with the phenyl ring of pY^83^ in NSF. Mutation of this residue caused a significant decrease of more than 8-fold in activity for both p-nitrophenyl phosphate (pNPP) and the phospho-NSF-peptide (Fig. 3B-C), suggesting that this residue is important for PTP-MEG2 recognition of all substrates with phenyl rings. N-terminal to pY^83^, the side chain oxygen of S^81^ formed a hydrogen bond with the carbonyl oxygen of the main chain of R^332^ of PTP-MEG2. The carbonyl oxygen of the main chain of S^81^ formed a hydrogen bond with the amide group of G^334^, and L^82^ formed hydrophobic interactions with the side chain of D^335^ (Fig. 3A). Specifically, mutation of G^334^ to R impaired the activity of PTP-MEG2 towards the NSF-pY^83^ phospho-peptide but did not affect on its intrinsic phosphatase activity as measured by using pNPP as a substrate, suggesting that G^334^ plays an important role in the recognition of the N-terminal conformation of the peptide substrate (Fig. 3B-C and Supplemental Fig. 4A-B).

**Figure 3.**
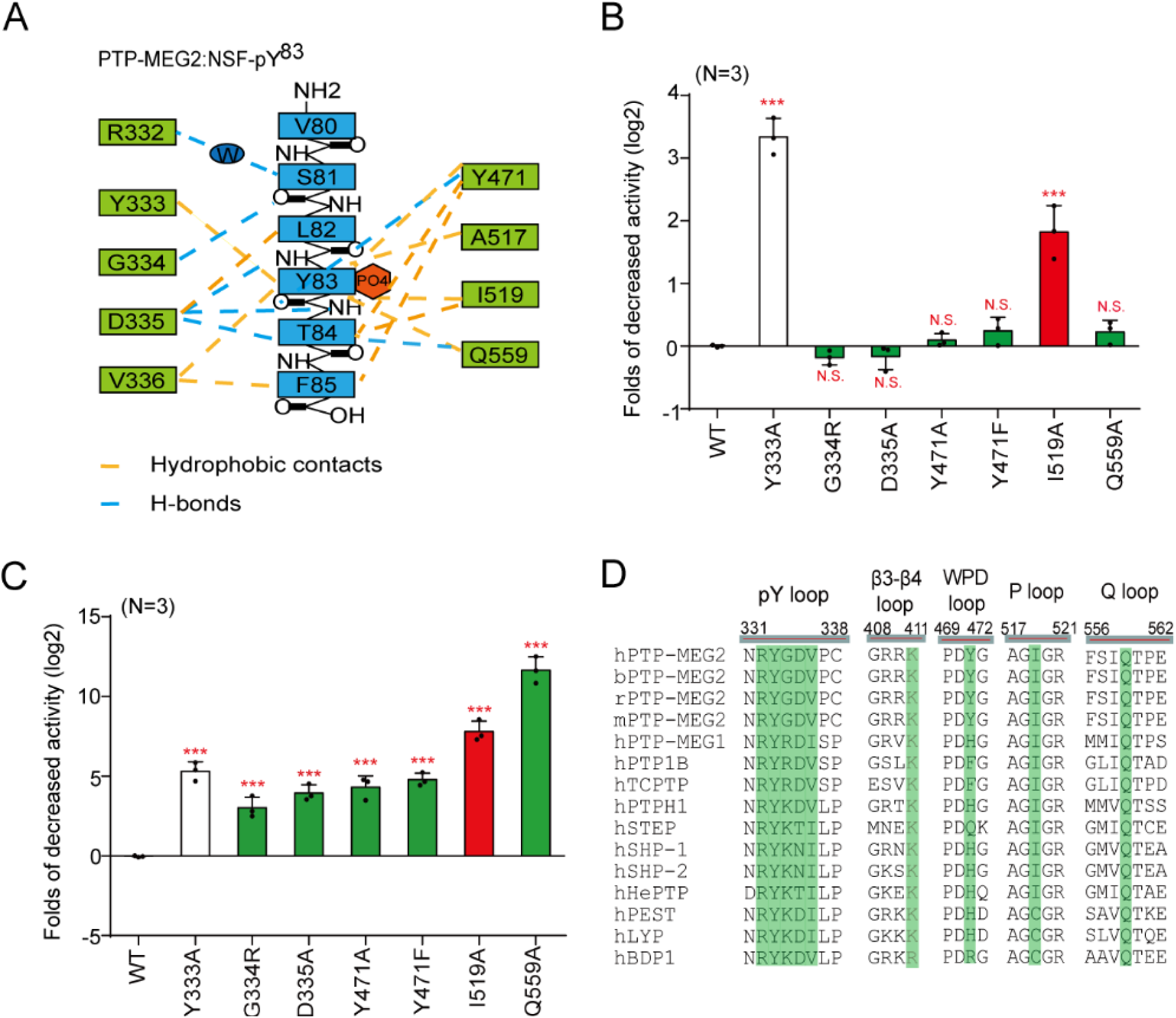
Molecular determinants of PTP-MEG2 interaction with the NSF-pY^83^ site. (A). Schematic representation of interactions between PTP-MEG2 and the NSF-pY^83^ site. (B). Relative values of the decreased phosphatase activities of different PTP-MEG2 mutants towards pNPP (p-Nitrophenyl phosphate) compared with wild-type PTP-MEG2. (C). Fold-change decreases in the phosphatase activities of different PTP-MEG2 mutants towards the NSF-pY^83^ phospho-segment compared with wild-type PTP-MEG2. The red column indicates that the mutation sites have greater effects on the enzyme activities toward phospho-NSF segement than the PNPP. The green column indicates that the mutation sites only have significant effects on the enzyme activity toward NSF phospho-segement but not pNPP. The white column stands for the mutants that have similar effects on pNPP and NSF phospho-segement. (D). Sequence alignment of PTP-MEG2 from different species and with other PTP members. Important structural motifs participating in phosphatase catalysis or the recognition of the NSF-pY^83^ site are shown, and key residues contributing to the substrate specificity are highlighted. *** P<0.001; PTP-MEG2 mutants compared with the control. Data were obtained from 3 independent experiments. N.S. means no significant difference. All the data were analyzed using one-way ANOVA.

The D^335^ in the pY binding loop of PTP-MEG2 is also critical for determining the peptide orientation of the substrate by forming important hydrogen bonds with the main chain amide and carbonyl groups of pY^83^ and T^84^ in NSF. Residues C-terminal to NSF-pY^83^, T^84^ and F^85^ formed substantial hydrophobic interactions with V^336^, F^556^, Q^559^, and Y^471^ (Fig. 3A). Accordingly, mutations D^335^A or Q^559^A showed no significant effects on pNPP activity but substantially decreased their activities towards the phospho-NSF peptide (Fig. 3B-C and Supplemental Fig. 4A-B). Mutation I^519^A caused a decrease in the intrinsic activity of PTP-MEG2 and a further decrease of approximately 256-fold in its ability to dephosphorylate phospho-NSF-peptide. Moreover, Y^471^ formed extensive hydrophobic interactions with T^84^ and F^85^ and also a hydrogen bond with the carboxyl group of pY^83^. Mutation of Y^471^ to either A or F greatly reduced its activity towards the phospho-NSF-E^79^-pY^83^-K^87^ peptide but had few effects on pNPP dephosphorylation. Taken together, the structural analyses and enzymological studies identified G^334^, D^335^, Y^471^, I^519^, and Q^559^ as critical residues for the substrate recognition of NSF by PTP-MEG2. Importantly, although none of these residues are unique to PTP-MEG2, the combination of these residues is not identical across the PTP superfamily but is conserved in PTP-MEG2 across different species, highlighting the important roles of these residues in mediating specific PTP-MEG2 functions (Fig. 3D and Supplemental Fig. 5).

### Molecular determinants of Q^559^:D^335^ for the substrate specificity of PTP-MEG2

The pY+1 pocket is an important determinant of substrate specificity in different PTP superfamily members (Barr et al., 2009; Li et al., 2016; Wang et al., 2014; Yu et al., 2011). The pY+1 pocket of PTP-MEG2 was found to consist of D^335^, V^336^, F^556^, and Q^559^ (Fig. 4A). Unique to the PTP-MEG2/NSF-E^79^-pY^83^-K^87^ complex structure, a relatively small T^84^ residue occurred at the pY+1 position, in contrast to L^993^ in the PTP1B/EGFR-pY^992^ complex structure, D^76^ in the LYP/SKAP-HOM-pY^75^ complex structure, and L^1249^ in the PTPN18/phospho-HER2-pY^1248^ complex structure (Fig. 4A). Although several PTPs have an equivalent D: Q pair similar to D^335^:Q^559^ of PTP-MEG2 determining the entrance of the pY+1 residue into the pY+1 pocket, such as PTP1B, LYP, PTPN18, STEP, PTP-MEG1, SHP1, PTPH1, and SHP2, structural analysis indicated that the Cβ between Q^559^ and D^335^ is the smallest in PTP-MEG2, at least 1 Å narrower than the other PTP structures examined (Fig. 4B). The narrower pY+1 pocket entrance could be a unique feature for substrate recognition by PTP-MEG2.

**Figure 4.**
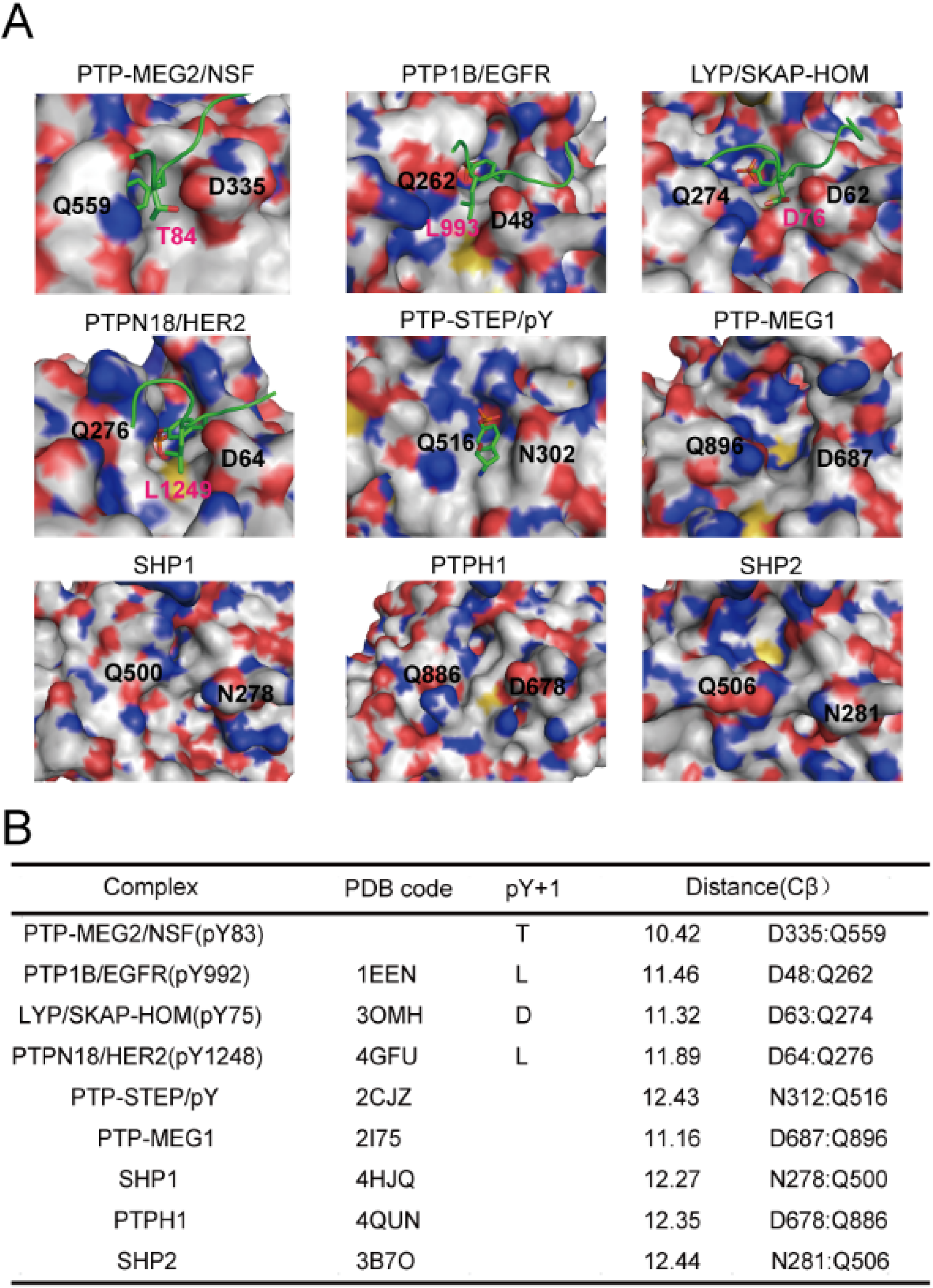
The tip opening of the pY+1 pocket is critical for PTP-MEG2 substrate specificity. (A). Surface representations of the complex structures of PTP-MEG2-C^515^A/D^470^A/NSF-pY^83^, PTP1B-C^215^A/EGFR-pY^992^ (PDB:1EEN), LYP-C^227^S/SKAP-HOM-pY^75^ (PDB:3OMH), PTPN18-C^229^S/HER2-pY^1248^ (PDB:4GFU), and PTP-STEP/pY (PDB:2CJZ). Crystal structures of PTP-MEG1 (PDB:2I75), SHP1 (PDB:4HJQ), PTPH1 (PDB:4QUN) and SHP2 (PDB: 3B7O). The pY+1 sites are highlighted. (B). Summary of the distance between the Cβ atoms of D^335^ and Q^559^ (corresponding to PTP-MEG2 number) of the pY+1 pocket in PTP-MEG2 and other classical non-receptor PTPs bearing the same residues at similar positions.

### PTP-MEG2 regulates two different steps of catecholamine secretion through dephosphorylation of different substrates

We then infected primary chromaffin cells with lentivirus encoding wild-type PTP-MEG2 or different mutants for electrochemical investigation of the structure-function relationship of PTP-MEG2 in the regulation of catecholamine secretion (Fig. 5A, Supplemental Fig. 6A-H and Supplemental Fig. 7A-D). In addition to the wild-type PTP-MEG2, we chose 6 PTP-MEG2 mutants, including G^334^R, D^335^A, Y^471^A, Y^471^F, I^519^A, and Q^559^A, whose positions are determinants of the interactions between PTP-MEG2 and NSF from the pY^83^-1 position to the pY^83^+2 position (Fig. 5B). Cells with approximately 15-fold of overexpressed exogenous PTP-MEG2 wild type or mutants than the endogenous PTP-MEG2 were selected for electrochemical studies (Supplemental Fig. 6E-F). The PTP-MEG2 mutations did not significantly affect its interaction with NSF, suggesting that residues not located within the active site of PTP-MEG2 may also participate in NSF associations (Supplemental Fig. 6G-6H). The overexpression of wild-type PTP-MEG2 significantly increased both the number and amplitude of the amperometric spikes, which are indicators of the release probabilities of individual vesicles and their quantal sizes, respectively (Fig. 5C-E and Supplemental Fig. 7E). In contrast, the expression of G^334^R, D^335^A, Y^471^A, Y^471^F, I^519^A, and Q^559^A all significantly decreased the quantal size, the release probabilities of individual vesicles, the half-width and the rise rate of each spike (Fig. 5C-E, Supplemental Fig. 7E-F and Supplemental Fig. 7H). However, there was no difference in the rise time between these mutations and the wild type (Supplemental Fig. 7G). These results suggest that the interaction of PTP-MEG2 with NSF is important for controlling vesicle size and the release probability of catecholamine secretion.

**Figure 5.**
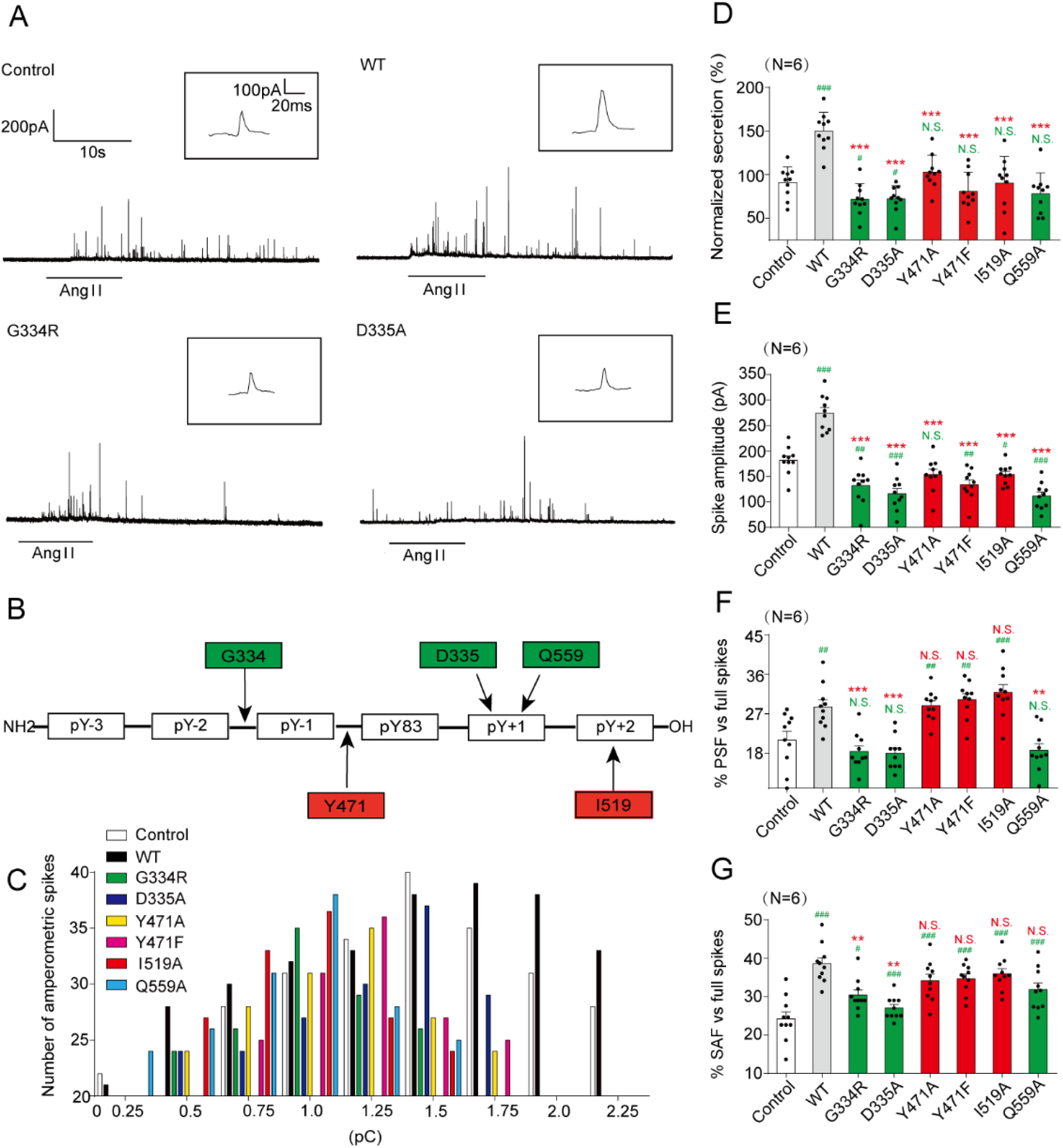
Effects of different PTP-MEG2 mutants on properties of catecholamine secretion from primary chromaffin cells. (A). Primary mouse chromaffin cells were transduced with a lentivirus containing the gene for wild-type PTP-MEG2 or different mutants with a GFP tag at the C-terminus. Positive transfected cells were confirmed with green fluorescence and selected for electrochemical analysis. Typical amperometric current traces evoked by AngII (100 nM for 10 seconds) in the control (transduced with control vector) (top left panel), WT (top right panel), G^334^R (bottom left panel), and D^335^A (bottom right panel) are shown. (B). The schematic diagram shows key residues of PTP-MEG2 defining the substrate specificity adjacent to the pY^83^ site of NSF. (C). The distribution of the quantal size of chromaffin cells transduced with lentivirus containing the genes encoding different PTP-MEG2 mutants. (D). Statistical diagram of the quantal size in Fig. 5A. The secretion amount of each group was standardized with respect to the control group. (E-G). Calculated parameters of secretory dynamics, including the spike amplitude (E), PSF frequency (F), and SAF frequency (G). D-G, * indicates PTP-MEG2 mutants overexpression group compared with the WT overexpression group. # indicates the overexpression group compared with the control group. **, P<0.01; *** P<0.001 and #, P<0.05; ##, P<0.01; ### P<0.001. N.S. means no significant difference. All the data were analyzed using one-way ANOVA.

Unexpectedly, the PTP-MEG2 mutants showed different effects on the probabilities of both PSF and SAF (Fig. 5F-G). G^334^R mutation which disrupted the recognition of the N-terminal conformation of the peptide substrate of PTP-MEG2, and D^335^A and Q^559^A, which are the determinants of pY+1 substrate specificity, significantly reduced the AngII-induced foot probabilities. The mutations of I^519^ and Y^471^, which formed specific interactions with T^84^ and F^85^ of NSF and are determinants of the C-terminal region of the central phospho-tyrosine involved in the substrate specificity of PTP-MEG2, showed no significant effects (Fig. 5B, Fig. 5F-G and Supplemental Fig. 7I-N). These results indicate that PTP-MEG2 regulates the initial opening of the fusion pore via a distinct structural basis from that of vesicle fusion, probably through dephosphorylating other unknown substrates. As D^335^A and Q^559^A of PTP-MEG2 maintained the occurrence of the foot probability, the unknown PTP-MEG2 substrate that regulates the fusion pore opening should have a small residue, such as G, A, S, or T at the pY+1 position. Conversely, the unknown PTP-MEG2 substrate should have a less hydrophobic residue at the pY+2 position because Y^471^F, Y^471^A, and I^519^A of PTP-MEG2 had no significant effects on the foot probability.

### Identification of new PTP-MEG2 substrates that contributed to the initial opening of the fusion pore

The effects of PTP-MEG2 mutations along the PTP-MEG2/NSF-phospho-segment interface on catecholamine secretion indicated that a PTP-MEG2 substrate other than NSF with distinct sequence characteristics contributes to the regulation of “foot probability” (Fig. 5). We therefore utilized this key information to search for new potential PTP-MEG2 substrates by bioinformatics methods (Fig. 6A). First, we searched for the keywords “fusion pore”, “secretory vesicle” and “tyrosine phosphorylation” using the functional protein association network STRING and the text mining tool PubTator, which resulted in a candidate list of 54 proteins. Second, we applied UniProt by selecting proteins located only in the membrane or vesicle, which limited the candidates to 28 members. Third, as our experiments were carried out in the adrenal gland, we used the Human Protein Atlas database for filtering to exclude the proteins which is not detectable in adrenal gland, which narrowed the candidate list to 23 proteins. Finally, we exploited the post-translational-motif database PhosphoSitePlus to screen candidate proteins with potential phospho-sites that matched the sequence requirements at both the pY+1 position and the pY+2 position, which are “G, S, A, T, V, P” and “G, A, S, T, C, V, L, P, D, H”, respectively. These positions were further evaluated by surface exposure if a structure was available. The combination of these searches produced 12 candidate lists with predicted pY positions (Fig. 6B) (Supplemental Table 1-3).

**Figure 6.**
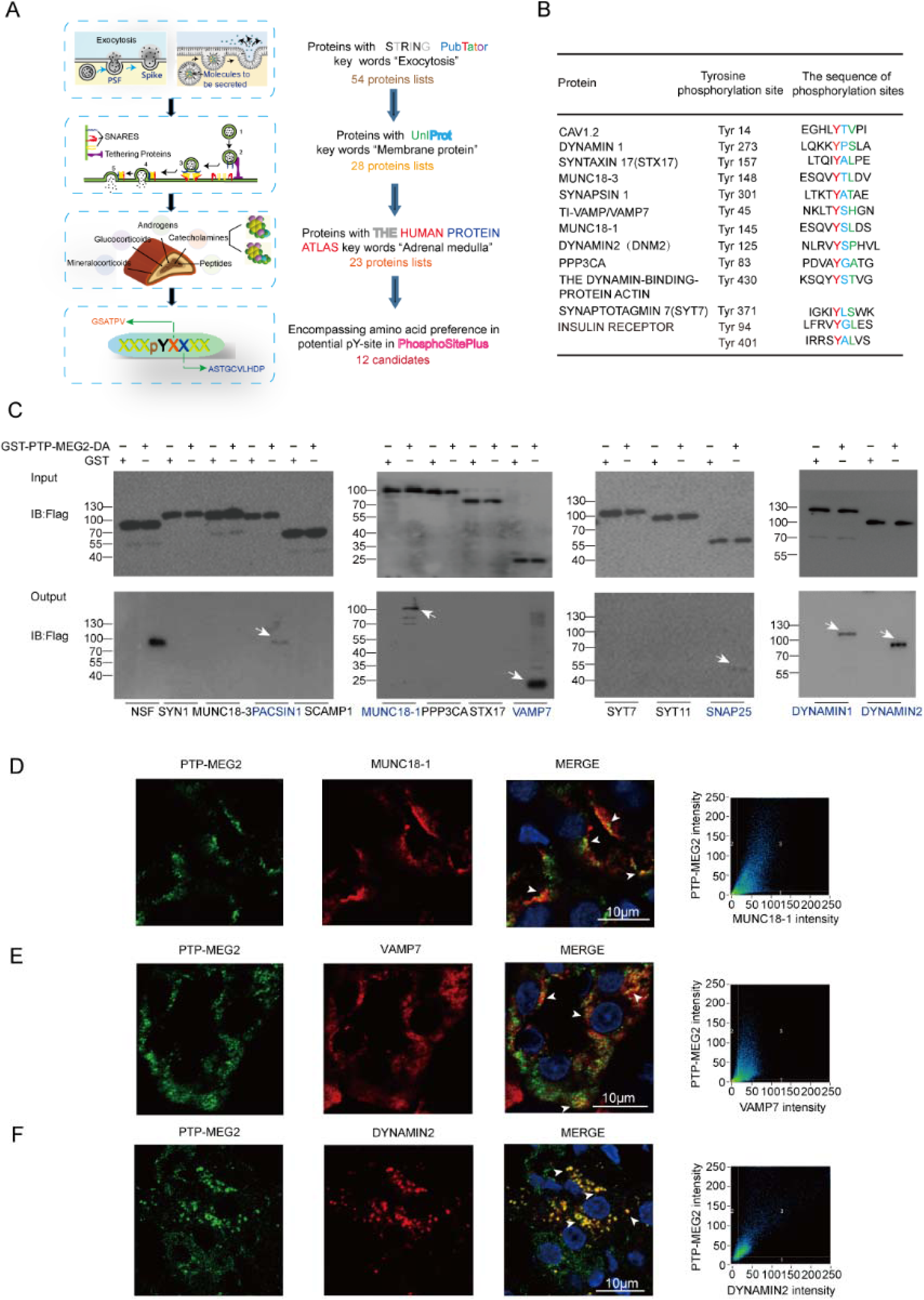
Identification of potential candidate substrates of PTP-MEG2 that participate in fusion pore initiation and expansion. (A). Flowchart for the workflow to predict the candidate substrates of PTP-MEG2 during fusion pore initiation and expansion. A total of 54 proteins were enriched with the functional protein association network STRING and the text mining tool PubTator by searching the keywords “fusion pore”, “secretory vesicle” and “tyrosine phosphorylation”. These proteins were filtered with UniProt by selecting proteins located only in the membrane or vesicle, which resulted in 28 candidates. The Human Protein Atlas database was then applied to exclude proteins with no expression in the adrenal gland. Finally, we used the post-translational-motif database PhosphoSitePlus to screen candidate proteins with potential phospho-sites that matched our sequence motif prediction at the pY+1 or pY+2 positions. (B). After the bioinformatics analysis, a total of 12 candidate PTP-MEG2 substrates that may participate in fusion pore initiation and expansion and their potential phospho-sites were displayed. (C). The GST pull-down assay suggested that PACSIN1, MUNC18-1, VAMP7, SNAP25, DYNAMIN1 and DYNAMIN2 directly interact with PTP-MEG2. PC12 cells were transfected with plasmids of candidate substrates, including SYN1, MUNC18-3, PACSIN1, SCAMP1, MUNC18-1, PPP3CA, STX17, VAMP7, SYT7, SYT11, SNAP25, DYNAMIN 1, and DYNAMIN2 stimulated with 100 nM AngII. The tyrosine phosphorylation of these proteins was verified by specific anti-pY antibodies (Supplemental Fig. 9A-E). The potential substrates of PTP-MEG2 in cell lysates were pulled down with a GST-PTP-MEG2-D^470^A trapping mutant and then detected by Western blotting. White arrow stands for Western blot band consistent with the predicted molecular weight of the potential PTP-MEG2 substrate. (D-F). Co-immunostaining assays of PTP-MEG2 with potential substrates in the adrenal medulla. MUNC18-1, VAMP7 and DYNAMIN2 all showed strong co-localization with PTP-MEG2 after 100 nM AngII stimulation in the adrenal medulla. White arrow stands for co-localization of PTP-MEG2 with MUNC18-1 or VAMP7. The Pearson’s correlation coefficients for D, E and F were 0.61, 0.65 and 0.79 respectively. The co-immunostaining results of PTP-MEG2 with other potential substrates are shown in Supplemental Fig. 8.

To biochemically characterize whether these proteins are substrates of PTP-MEG2, we transfected the plasmids encoding the cDNAs of these candidate proteins into PC12 cells, stimulated the cells with AngII, and performed a pull-down assay with the GST-PTP-MEG2-D^470^A trapping mutant or GST controls. The known PTP-MEG2 substrate NSF was used as a positive control. Notably, six candidates, including PACSIN1, MUNC18-1, VAMP7, SNAP25, DYNAMIN1 and DYNAMIN2 showed specific interactions with PTP-MEG2 after AngII stimulation in PC12 cells (Fig. 6C and Supplemental Fig. 8A-B). Whereas the protein of MUNC18-1, VAMP7, DYNAMIN2 and SNAP25 were readily detected in the adrenal medulla, DYNAMIN1 and PACSIN1 showed substantially lower expression than that in the liver and brain (Supplemental Fig. 8C). Moreover, whereas MUNC18-1, VAMP7 and DYNAMIN2 strongly co-localized with PTP-MEG2 in the adrenal medulla (Fig. 6D-F), the co-localization of SNAP25 and PACSIN1 with PTP-MEG2 was relatively weak (Supplemental Fig. 8D-E). Therefore, MUNC18-1, VAMP7 and DYNAMIN2 are more likely candidate PTP-MEG2 substrates which were further strengthened by the fact that the high potassium chloride- or AngII-stimulated tyrosine phosphorylation of these proteins in the adrenal medulla was significantly dephosphorylated by PTP-MEG2 in vitro (Supplemental Fig. 9A-C, E), whereas the tyrosine phosphorylation of SNAP25 had no obvious change (Supplemental Fig. 9D-E).

### PTP-MEG2 regulated the initial opening of the fusion pore through dephosphorylating MUNC18-1-pY^145^

The predicted PTP-MEG2 dephosphorylation site on MUNC18-1 (also called STXBP1) is Y^145^, which is localized on the β sheet linker and forms extensive hydrophobic interactions with surrounding residues (Hu et al., 2011; Yang et al., 2015) (Fig. 7A and Supplemental Fig. 10A). Moreover, the phenolic-oxygen of Y^145^ forms specific hydrogen bonds with the main chain amide of F^540^ and the main chain carbonyl oxygens of I^539^ and G^568^ (Fig. 7B). These key interactions might be involved in regulating the arrangement of the arc shape of the three domains of MUNC18-1, by tethering the interface between domain 1 and domain 2. The phosphorylation of Y^145^ likely abolishes this H-bond network and changes its ability to associate with different snare complexes participating in vesicle fusion procedures (Fig. 7B). Interestingly, a missense mutation of MUNC18-1 Y^145^H was found to be associated with early infantile epileptic encephalopathy (Stamberger et al., 2017) (Fig. 7C).

Notably, the Y^145^H missense mutation may not be phosphorylated properly. We therefore overexpressed wild-type MUNC18-1, Y^145^A, a non-phosphorylable mutant, Y^145^E, a phosphomimetic mutant, Y^145^F, a non-phosphomimetic mutant, and the disease-related Y^145^H mutant in PC12 cells stimulated with AngII and then examined their abilities to interact with PTP-MEG2 trapping mutant. The GST pull-down results indicated that all MUNC18-1 Y^145^ mutations significantly decreased their ability to associate with PTP-MEG2 (Fig. 7D, supplemental Fig. 9G-9H). In contrast, mutation of the predicted phosphorylation sites of SNAP25, the SNAP25-Y^101^A, had no significant effect on their interactions with PTP-MEG2 (Supplemental Fig. 9F-H). These results suggested that pY^145^ is one of the major sites of MUNC18-1 regulated by PTP-MEG2 (Fig. 7D). Moreover, the effects of the muations of MUNC18-1 and DYNAMIN2 on protein stability were detected in response to CHX treatments. Consequently, the mutations of MUNC18-1 and DYNAMIN2 had no significant influence on the stability (Supplemental Fig. 9I).

**Figure 7.**
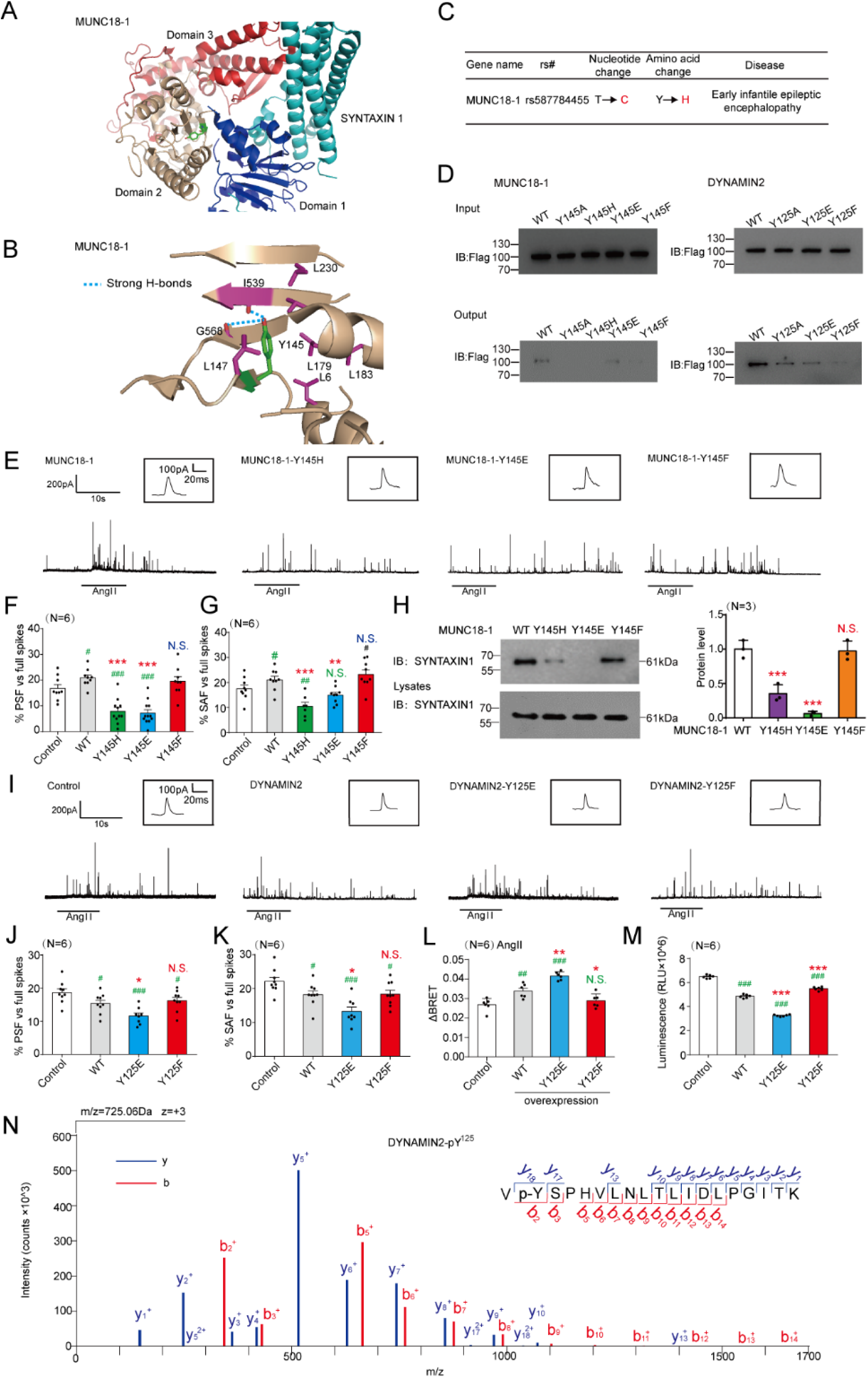
Dephosphorylation of MUNC18-1 at the pY^145^ site by PTP-MEG2 determines the “foot” probability of catecholamine secretion from chromaffin cells. (A). Structural representation and location of MUNC18-1-Y^145^ in the complex structure of MUNC18-1-SYNTAXIN1 (PDB: 3PUJ). (B). Detailed structural representation of Y^145^ and its interacting residues. Y^145^ of MUNC18-1 interacts with the main chain carboxylic oxygen of residues I^539^ and G^568^ and forms hydrophobic interactions with L^6^, I^539^, L^147^, L^179^, L^183^, and L^230^ to tether the arc shape of the three domains of MUNC18-1 (PDB: 3PUJ). (C). Association analysis of SNPs of MUNC18-1 with human disease. (D). Interactions of the PTP-MEG2-trapping mutants with the MUNC18-1-Y^145^ mutants and DYNAMIN2-Y^125^ mutants. PC12 cells were transfected with FLAG-MUNC18-1-Y^145^, FLAG-DYNAMIN2-Y^125^ and different mutations of the FLAG-MUNC18-1-Y^145^A, Y^145^H, Y^145^E or Y^145^F mutants, FLAG-DYNAMIN2-Y^125^A, Y^125^E, Y^125^F mutants, 24 hours before stimulation with 100 nM AngII, respectively. The cell lysates were then incubated with GST-PTP-MEG2-D^470^A for 4 hours with constant rotation. The potential PTP-MEG2 substrates were pulled down by GST-beads, and their levels were examined by the FLAG antibody with Western blotting. (E). Primary chromaffin cells were transduced with lentivirus containing the gene encoding wild-type MUNC18-1 or different mutants. These cells were stimulated with 100 nM AngII. The amperometric spikes were detected with CFE. Typical amperometric traces are shown. (F). The percentages of pre-spike foot for wild-type MUNC18-1 or different mutants were calculated. (G). The percentages of stand-alone foot for wild-type MUNC18-1 or different mutants were calculated. (H). The MUNC18-1-Y^145^ mutation decreased the interaction between MUNC18-1 and SYNTAXIN1. PC12 cells were transfected with plasmid encoding SYNTAXIN-1. The proteins in cell lysates were pulled down with purified GST-MUNC18-1-Y^145^ and the GST-MUNC18-1-Y^145^H/E/F, and detected with SYNTAXIN1 antibody. The right histogram shows the quantified protein levels. (I). Primary chromaffin cells were transduced with lentivirus containing the gene encoding wild-type DYNAMIN2 or different mutants. These cells were stimulated with 100 nM AngII. The amperometric spikes were detected with CFE. Typical amperometric traces are shown. (J). The percentages of pre-spike foot for wild-type DYNAMIN2 or different mutants were calculated. (K). The percentages of stand-alone foot for wild-type DYNAMIN2 or different mutants were calculated. (L). Plasmids encoding DYNAMIN2 (DYNAMIN2-WT, DYNAMIN2-Y^125^E or DYNAMIN2-Y^125^F) and LYN-YFP, AT1aR-C-RLUC were co-transferred into HEK293 cells at 1:1:1 ratio. The BRET experiment was performed to monitor the mutation effects of DYNAMIN2 on AT1aR endocytosis in response to AngII (1μM) stimulation. (M). A GTPase assay was performed on His-DYNAMIN2-WT (500ng), His-DYNAMIN2-Y^125^E (500ng), His-DYNAMIN2-Y^125^F (500ng). Purified DYNAMIN2 proteins were incubated with 5μM GTP in the presence of 500μM DTT for 90 minutes, followed by addition of GTPase-Glo™ Reagent and detection reagent. The luminescence were measured after 37-mins incubation. Bar representation of the results from six independent experiments. (N). Phosphorylated Y^125^ of DYNAMIN2 was identified by LC-MS/MS. The PTP-MEG2-DA trapping mutant was used to pull down potential PTP-MEG2 substrates from the adrenal lysates after AngII stimulation. Trypsin-digested potential PTP-MEG2 substrates were subjected to LC-MS/MS analysis. The doubly charged peptide with m/z 725.06 matches VYSPHVLNLTLIDLPGITK of DYNAMIN2 with Y^125^ phosphorylation. F-H, J-M * indicates MUNC18-1, DYNAMIN2 mutants overexpression group compared with the WT overexpression group. # indicates MUNC18-1, DYNAMIN2 overexpression group compared with the control group. *, P<0.05; **, P<0.01; *** P<0.001 and #, P<0.05; ##, P<0.01; ### P<0.001. N.S. means no significant difference. All the data were analyzed using one-way ANOVA.

We next infected primary chromaffin cells with a lentivirus encoding wild-type MUNC18-1, the MUNC18-1 Y^145^-tyrosine phosphorylation-deficient mutant Y^145^F, the MUNC18-1 Y^145^-tyrosine phosphorylation mimic mutant Y^145^E and the disease-related mutant Y^145^H. The Y^145^E is a phosphorylation mimic mutant of pY^145^, which generate a negative charge at this specific site. In contrast, the Y^145^F could not be phosphorylated because of lack of the phenolic oxygen. We thus examined the effects of these mutants using primary chromaffin cells harbouring approximately 20∼30 folds MUNC18-1 overexpression related to that of endogenous expression levels (Supplemental Fig. 10B-D). Interestingly, either Y^145^E or Y^145^H significantly reduced the PSF and SAF percentage of catecholamine secretion in response to AngII stimulation, whereas the phosphorylation-deficient mutant Y^145^F showed no significant effects on the PSF or SAF percentage compared to that of the wild type (Fig. 7E-G, and Supplemental Fig. 10 and 11).

To further dissect the mechanism underlying the phosphorylation of MUNC18-1 Y^145^ as well as the disease-related mutant Y^145^H in the regulation of hormone secretion, we compared the interactions of wild-type and mutant MUNC18-1 with the binding partner SYNTAXIN1 (Lim et al., 2013). Importantly, both the phosphorylation mimic mutant MUNC18-1-Y^145^E and the disease-related mutant Y^145^H significantly impaired the interaction between MUNC18-1 and SYNTAXIN1 (Fig. 7H, Supplemental Figure 11C). The effects of MUNC18-1 phospho-mimic Y^145^E or phospho-defective Y^145^F mutants on fusion pore dynamics are correlated with the data of their interaction with the SYNTAXIN1, thus indicating that the the binding between the MUNC18-1 and SYNTAXIN1, which was controlled by the phosphorylation status of Y^145^ of MUNC18-1, may play an important role in the formation of the foot during catecholamine secretion. In contrast, the dephosphorylation of pY^145^ of MUNC18-1 by PTP-MEG2 promoted initial pore opening and fusion. Collectively, these results suggested that the tyrosine phosphorylation of Y^145^ impaired the initial opening of the fusion pore in agonist-induced catecholamine secretion in primary chromaffin cells, probably via either disruption of the arc shape of MUNC18-1 or impairing the interaction between MUNC18-1 and Syntaxin1.

### PTP-MEG2 regulated the initial opening of the fusion pore through dephosphorylating DYNAMIN2-pY^125^

Except for MUNC18-1, two additional substrates of PTP-MEG2, which are the Y^45^ site of VAMP7 and the Y^125^ site of DYNAMIN2, were confirmed by GST-pull down assay together with specific mutating effects on the corresponding phosphorylation sites (Fig. 7D, supplemental Fig. 9F-H). It was well known that both DYNAMIN1 and DYNAMIN2 are active regulators of pore fusion dynamics, involved in the pore establishment, stabilization, constriction and fission processes (Jones et al., 2017; Shin et al., 2018; Zhao et al., 2016). The dephosphorylation site of DYNAMIN2 by PTP-MEG2, the Y^125^, is located in a β-sheet of DYNAMIN GTPase domain according to the solved DYNAMIN1 structure (PDB ID: 5D3Q). Previous studies have identified that phosphorylation of DYNAMIN close to the G domain significantly increased its GTPase activity, thus promoting endocytosis (Kar et al., 2017). We therefore hypothesized that the phosphorylation of the DYNAMIN2 pY^125^ site increased pore fission by up-regulating its GTPase activity, whereas dephosphorylation of DYNAMIN2 pY^125^ site by PTP-MEG2 reversed it. Consistent with this hypothesis, the primary chromaffin cells transduced by lentivirus encoding DYNAMIN2 wild type and the Y^125^F mutant significantly decreased percentage of the PSF and SAF of catecholamine secretion elicited by AngII, whereas the phosphorylation mimic mutant DYNAMIN2 Y^125^E mutant exhibited significantly more reduction for the PSF and SAF percentage (Fig. 7I-K and Supplemental Fig. 12). It has been established that the DYNAMIN mediated pore fission process is closely correlated with the receptor endocytosis. Here, whereas overexpression of the DYNAMIN2 wild type or Y^125^E phospho-mimic mutant markedly increased the AngII-induced AT1aR endocytosis, the Y^125^F mutant has no significant effect, suggesting DYNAMIN2 Y^125^E is a gain-of-function mutant to promote the membrane fission (Fig. 7L) (Liu et al., 2017). Consistent with these cellular results, the in vitro GTPase activity further suggested that the phosphorylation of DYNAMIN2 at pY^125^ site up-regulated its activity (Fig. 7M). Moreover, the LS-MS/MS verified the phosphorylation of DYNAMIN2 at pY^125^ site in primary chromaffin cells under AngII stimulation (Fig. 7N, Supplemental Table 4), indicating that DYNAMIN2 pY^125^ phosphorylation occurred under AngII stimulation and participated in the AngII-induced catecholamine secretion.

### Molecular mechanisms of the PTP-MEG2/MUNC18-1-pY^145^ and PTP-MEG2-DYNAMIN2-pY^125^ interaction

The *k*_cat_/*K*_m_ of PTP-MEG2 towards a phospho-segment derived from MUNC18-1-pY^145^ is very similar to that obtained with a phospho-segment derived from the known substrate pY^83^ site of NSF (Supplemental Fig. 4A-B and Supplemental Fig. 14). We therefore crystallized the PTP-MEG2 trapping mutant with the MUNC18-1-E^141^-pY^145^-S^149^ phospho-segment and determined the complex structure at 2.2 Å resolution (Table 1). The 2Fo-Fc electro-density map allowed the unambiguous assignment of seven residues of the phospho-MUNC18-1-E^141^-pY^145^-S^149^ segment in the crystal structure (Fig. 8A). Importantly, comparing with the phospho-NSF-E^79^-pY^83^-K^87^ segment, the phospho-MUNC18-1-E^141^-pY^145^-S^149^ displayed different interaction patterns with the residues in the PTP-MEG2 active site, forming new interactions with R^409^ and R^410^ but lost interactions with Y^471^ and I^519^ (Fig. 8B). Whereas PTP-MEG2 Y^471^ formed extensive hydrophobic interactions with NSF-T^84^ and NSF-F^85^, as well as a well-defined hydrogen bond (2.4 Å) with the carbonyl oxygen of NSF-pY^83^, PTP-MEG2 Y^471^ formed only a weaker H-bond with the carbonyl oxygen of MUNC18-1-pY^145^ due to a 0.62 Å shift of Y^471^ away from the central pY substrate (Fig. 8C and Supplemental Fig. 13A). Similarly, PTP-MEG2 I^519^ did not form specific interactions with the MUNC18-1-E^141^-pY^145^-S^149^ segment except for the central pY. In contrast, PTP-MEG2 I^519^ formed specific hydrophobic interactions with NSF-T^84^ (Fig. 8D and Supplemental Fig. 13B). Consistently, mutations of PTP-MEG2 Y^471^A, Y^471^F or I^519^A significantly decreased the phosphatase activity towards the phospho-NSF-E^79^-pY^83^-K^87^ segment (Fig. 3C) but had no significant effect on the phospho-MUNC18-1-E^141^-pY^145^ -S^149^ segment (Fig. 8E and Supplemental Fig. 14). Meanwhile, R^409^A and R^410^A mutations substantially decreased the catalytic activity toward phospho-segements derived from both MUNC18-1-pY^145^ and DYNAMIN2 pY^125^ sites, indicating that recognition of these substrates was mediated by common residues (Fig. 8F and Supplemental Fig. 14). Notably, the PTP-MEG2 catalytic domain mutations shows no significant effects on the interactions between PTP-MEG2 with the NSF, MUNC18-1 and DYNAMIN2, indicating other domain of PTP-MEG2, such as its FYVE domain, participated into the association of PTP-MEG2 with these proteins (Supplemental Fig. 6G and Supplemenatl Fig. 13C-F). With CFE, we further demonstrated that R^409^A and R^410^A mutations decreased the probability of both PSF and SAF (Fig. 8G, Supplemental Fig. 15). Combined with the results that PTP-MEG2 Y^471^A, Y^471^F and I^519^A affect only the spike number and amount but show not the foot probability (Fig. 5B-G), these data suggested that PTP-MEG2 regulated intracellular vesicle fusion by modulating the NSF-pY^83^ phospho-state but regulated the process of vesicle fusion pore initiation by dephosphorylating MUNC18-1 at pY^145^ site and DYNAMIN2 at pY^125^ site (Fig. 8H).

**Figure 8.**
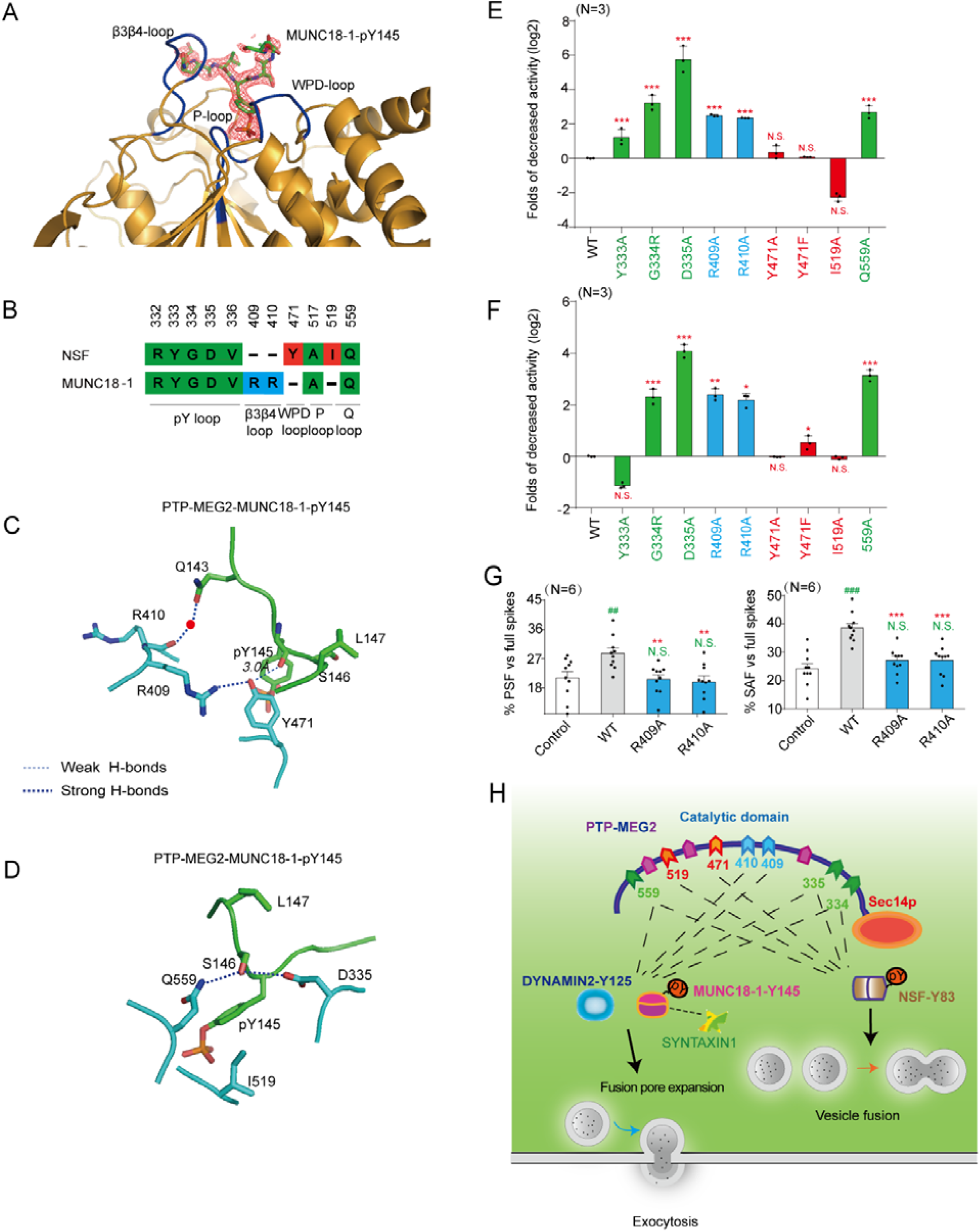
The structural details of the PTP-MEG2-MUNC18-1-Y^145^ interaction and catalytic bias of PTP-MEG2 towards NSF-pY^83^ and MUNC18-1-pY^145^. (A). The 2Fo-Fc annealing omit map (contoured at 1.0σ) around MUNC18-1-pY^145^ phospho-segment. (B). Comparison of residues of PTP-MEG2 interacting with NSF and MUNC18-1. Amino acid residues of NSF and MUNC18-1 are coloured as follows: green, residues interacting with both NSF and MUNC18-1; red, residues specifically contributing to NSF recognition; blue, residues selectively contributing to MUNC18-1 interaction. (C). The structural alteration of the interactions surrounding Y^471^ of PTP-MEG2 with the MUNC18-1-pY^145^ site. (D). The structural alteration of the interactions surrounding I^519^ of PTP-MEG2 with the MUNC18-1-pY^145^ site. (E). Relative phosphatase activities of different PTP-MEG2 mutants towards the MUNC18-1-pY^145^ phospho-segment compared with wild-type PTP-MEG2. (F). Relative phosphatase activities of different PTP-MEG2 mutants towards the DYNAMIN 2-pY^125^ phospho-segment compared with wild-type PTP-MEG2. E-F Residues are coloured according to Figure 8B. (G).The percentages of PSF and SAF for PTP-MEG2-R^409^A or R^410^A were calculated. (H). Schematic illustration of the PTP-MEG2-regulated processes of vesicle fusion and secretion in chromaffin cells via the dephosphorylation of different substrates with distinct structural basis. PTP-MEG2 regulates vesicle fusion and vesicle size during the catecholamine secretion of adrenal gland by modulating the phosphorylation state of the pY^83^ site of NSF, which relies on the key residues G^334^, D^335^ (pY loop), Y^471^ (WPD loop), I^519^ (P-loop) and Q^559^ (Q loop). PTP-MEG2 regulates the fusion pore initiation and expansion procedures of catecholamine secretion by the adrenal gland (also designated as foot probability) by modulating the newly identified substrate MUNC18-1 at its pY^145^ site and DYNAMIN2 at its pY^125^, through distinct structural basis from that of its regulation of NSF phosphorylation. E-F, * P<0.05, * P<0.01, *** P<0.001; PTP-MEG2 mutants compared with the control. N.S. means no significant difference. Data were obtained from 3 independent experiments. All the data were analyzed using one-way ANOVA.

## DISCUSSION

Posttranslational modifications of secretion machinery proteins are known as powerful ways to regulate exocytosis. In contrast to the well-characterized serine/threonine phosphorylation, the importance of tyrosine phosphorylation in exocytosis has only recently begun to be appreciated (Cijsouw et al., 2014; Gabel et al., 2019; Jewell et al., 2011; Laidlaw et al., 2017; Meijer et al., 2018; Seino et al., 2009). In addition to the phosphorylation of NSF at its pY^83^ site, recent studies have shown that tyrosine phosphorylation of MUNC18-3 at the pY^219^ and pY^527^ sites, Annexin-A2 at pY^23^, and MUNC18-1 at pY^473^ actively participates in the vesicle release machinery to explicitly regulate exocytosis processes (Gabel et al., 2019; Jewell et al., 2011; Meijer et al., 2018). The tyrosine phosphorylation at specific sites of the signalling molecule is precisely regulated by protein tyrosine kinases and protein tyrosine phosphatases (Tonks, 2006; Yu and Zhang, 2018). Although tyrosine kinases such as the insulin receptor, Src and Fyn are acknowledged to play critical roles in hormone secretion (Jewell et al., 2011; Meijer et al., 2018; Oakie and Wang, 2018; Soares et al., 2013), only a very few PTPs that regulate the vesicle release machinery have been identified, and the structural basis of how these PTPs selectively dephosphorylate the key tyrosine phosphorylation sites governing exocytosis was unknown. In the current study, we demonstrated that PTP-MEG2 is an important regulator of hormone secretion from the chromaffin cell, using a selective PTP-MEG2 inhibitor in combination with cellular and electrochemical amperometric recording. The current study extended the regulatory role of PTP-MEG2 in various steps of exocytosis in hormone secretion beyond the previously known simple vesicle fusion step of the immune system (Huynh et al., 2004). We then determined the crystal structure of PTP-MEG2 in complex with the pY^83^ phospho-segment of the NSF, the key energy provider for disassembling fusion-incompetent cis SNARE complexes in the process of vesicle fusion in immunocytes (Huynh et al., 2004). The complex structure not only revealed the structural rearrangement in PTP-MEG2 in response to binding of the substrate NSF, identifying Q^559^:D^335^ as the key pair for substrate specificity of the pY+1 site, but also provided clues that PTP-MEG2 regulated the initial opening of the fusion pore through another unknown substrate. Fortunately, we were able to deduce the signature of the pY+1 and pY+2 positions of this unknown substrate by carefully inspecting the PTP-MEG2/phospho-NSF-E^79^-pY^83^-K^87^ complex structure and analysing the functional data of the PTP-MEG2 interface mutants. Further bioinformatics studies and cellular and physiological experiments enabled us to discover that PTP-MEG2 regulates the initial opening of the fusion pore by modulating the tyrosine phosphorylation states of the MUNC18-1 at the pY^145^ site and the DYNAMIN2 at the pY^125^ site. Therefore, we have revealed that PTP-MEG2 regulates different steps of the exocytosis processes via dephosphorylating distinct substrates. PTP-MEG2 regulates the vesicle size and vesicle-vesicle fusion step by dephosphorylating NSF at its NSF-pY^83^ site, whereas it regulates the process of LDCV fusion pore initiation and expansion by controlling specific phosphorylation sites of MUNC18-1 and DYNAMIN2. Moreover, our studies highlight that the combination of structural determination and functional delineation of the interface mutants of the protein complex is a powerful approach to characterizing the signalling events and identifying unknown downstream signalling molecules.

Fusion pore opening and expansion is thought to be a complex process requiring the docking of apposed lipid bilayers and involvement of multiple proteins to form a hemi-fusion diaphragm including SNAREs, MUNC18-1, and DYNAMIN (Baker and Hughson, 2016; Hernandez et al., 2012; Jones et al., 2017; Mattila et al., 2015; Sudhof and Rothman, 2009; Zhao et al., 2016). Dissection of the molecular mechanism underlying pore fusion dynamics in exocytosis is challenging because direct observation of this process is difficult to achieve due to the short expansion time and the tiny size of the pore (Baker and Hughson, 2016; Gaisano, 2017; Hong and Lev, 2014). Importantly, the DYNAMINs are key terminators of the fusion pore expansion by scissoring vesicles from the cell membrane (Eitzen, 2003) (Jones et al., 2017; Shin et al., 2018; Zhao et al., 2016). The large scale phosphoproteomic screens have identified multiple tyrosine phosphorylation sites of DYNAMIN, such as that occurring at pY^80^, pY^125^, pY^354^ and pY^597^ (Ballif et al., 2008) (Mallozzi et al., 2013). Among them, the Y^597^ was a well-studied phosphorylation site of DYNAMIN, which was previously reported to increase self-assembly and GTP hydrolysis in both DYNAMIN1 and 2 (Kar et al., 2017). However, the Y^597^ of DYNAMIN was probably not the target of PTP-MEG2 because the interaction of Y^597^ or A^597^ with PTP-MEG2 exhibited no significant differences. However, the regulation of other tyrosine phosphorylation site, such as pY^125^, as well as how this particular tyrosine phosphorylation participated in a selective secretion process, has not been elucidated. Here, by using pharmacological approach with high selectivity, functional alanine mutagenesis according to structural characterization, combined with bioinformatics, electrochemistry and enzymology, we have demonstrated that PTP-MEG2 negatively regulated the pY^125^ phosphorylation state during the catecholamine secretion, which promoted the cessation of the fusion pore expansion through increasing its GTP activity.

In addition to DYNAMIN, MUNC18-1 and its closely related subfamily members have been demonstrated to participate in several processes during vesicle secretion by interacting with SNAREs, such as docking, priming and vesicle fusion (Chai et al., 2016; Cijsouw et al., 2014; Fisher et al., 2001; Gulyas-Kovacs et al., 2007; He et al., 2017; Korteweg et al., 2005; Ma et al., 2013; Ma et al., 2015; Meijer et al., 2018; Sitarska et al., 2017). Importantly, the tyrosine phosphorylation of MUNC18-1 at Y^473^ was recently reported as a key step in modulating vesicle priming by inhibiting synaptic transmission and preventing SNARE assembly (Meijer et al., 2018). In addition, the tyrosine phosphorylation of MUNC18-3 on Y^521^ was essential for the dissociation of MUNC18-3 and SYNTAXIN4 (Umahara et al., 2008). Moreover, the dephosphorylation of MUNC18-1-Y^145^ was suggested to be essential in maintaining the association between MUNC18-1 and SYNTAXIN1 (Lim et al., 2013). In the present study, we demonstrated that the MUNC18-1 Y^145^E phospho-mimic mutation, but not the non-phosphorylated mutation Y^145^F, significantly decreased the PSF and the SAF probability in cultured primary chromaffin cells. Notably, either the Y^145^E phospho-mimic mutation or the epileptic encephalopathy associated Y^145^H mutant disrupted their interactions with SYNTAXIN1. Structural inspection also suggested that both phosphorylation of Y^145^ and Y^145^H mutant could destabilize the arc shape of native MUNC18-1. Therefore, it’s very likely that either the association of the MUNC18-1 with the SYNTAXIN1 or the maintenance of the arc shape of the MUNC18-1 actively participated in initial pore opening and expansion. Future studies by solving the fusion machinery structures encompassing with MUNC18-1 at different stages with high resolutions, as well as more detailed fusion pore dynamics analysis using in vitro reconstitution system could provide deeper insights for these key events in proe fusion processes. In the present study, the structural analysis of PTP-MEG2 in complex with MUNC18-1-pY^145^ and the enzymatic analysis confirmed that PTP-MEG2 regulated two different substrates, the MUNC18-1-pY^145^ and the DYNAMIN2-pY^125^ with similar structural features and generated similar effects on fusion pore dynamics. Therefore, our studies exemplified how a PTPase regulated one important physiological process through two different substrates, adding new information of the fusion pore regulation from another aspect.

Notably, the MUNC18-1 Y^145^H mutation is a known SNP that is associated with epileptic encephalopathy (Stamberger et al., 2017). Y^145^H behaves similarly to the MUNC18-1 phosphorylation mimic mutant Y^145^E by disrupting its interaction with SYNTAXIN1 and reducing the probability of PSF elicited by AngII in primary chromaffin cells. This observation provided a clue for the pathological effects of the MUNC18-1 Y^145^H mutation. In addition to MUNC18-1, we found that VAMP7 interacted with PTP-MEG2 via their Y^45^ tyrosine phosphorylation site, respectively. DYNAMIN1, PASCIN1 and SNAP25 are also potential PTP-MEG2 substrates depending on specific cellular contexts. The functions of VAMP7 phosphorylation at the Y^45^ site and its dephosphorylation by PTP-MEG2, the interaction of PTP-MEG2 with DYNAMIN1, etc. in the exocytosis process await further investigation.

Finally, by solving the two crystal structures of PTP-MEG2 in complex with two substrates, the phospho-NSF-E^79^-pY^83^-K^87^ segment and the phospho-MUNC18-1-E^141^-pY^145^ -S^149^ segment, we revealed that PTP-MEG2 recognized these functionally different substrates through distinct structural bases. Whereas K^411^, Y^471^ and I^519^ contributed most to the selective interaction of PTP-MEG2 with NSF, another set of residues, including R^409^ and R^410^, mediated the specific binding of PTP-MEG2 to MUNC18-1 (Fig. 8B). Most importantly, mutating Y^471^ and I^519^ to A significantly decreased the activity of PTP-MEG2 towards the phospho-NSF-E^79^-pY^83^-K^87^ segment but not the phospho-MUNC18-1-E^141^-pY^145^-S^149^ segment. The biochemical data agreed well with the functional data that PTP-MEG2 Y^471^A and I^519^A of PTP-MEG2 affected only the vesicle fusion procedure but not the fusion pore opening and expansion processes. These data not only indicate that PTP-MEG2 regulates different steps of exocytosis through different substrates in an explicit temporal and spatial context but also afforded important guidance for the design of selective PTP-MEG2 inhibitors according to the different interfaces between PTP-MEG2 and its substrates to explicitly regulate specific physiological processes, supporting the hypothesis of “substrate-specific PTP inhibitors” (Doody and Bottini, 2014). The design of such inhibitors will certainly help to delineate specific roles of PTP-MEG2 in different physiological and pathological processes.

In conclusion, we have found that PTP-MEG2 regulates two different processes of exocytosis during catecholamine secretion, namely, vesicle fusion and the opening and extension of the fusion pore, through two different substrates with distinct structural bases. We achieved this knowledge by determining the complex structure and performing functional delineation of the protein complex interface mutants. The present study supports the hypothesis that the tyrosine phosphorylation of secretion machinery proteins is an important category of regulatory events for hormone secretion and is explicitly regulated by protein tyrosine phosphatases, such as PTP-MEG2. Dissecting the molecular and structural mechanisms of such modulation processes will provide an in-depth understanding of the exocytosis process and guide further therapeutic development for exocytosis-related diseases, such as epileptic encephalopathy (Stamberger et al., 2017).

## METHODS

## KEY RESOURCES TABLE

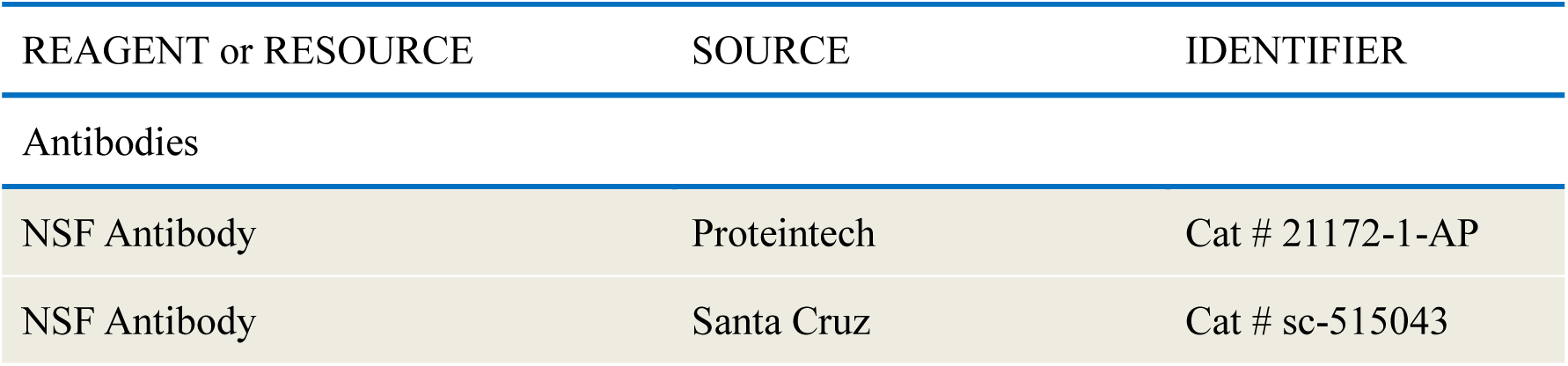
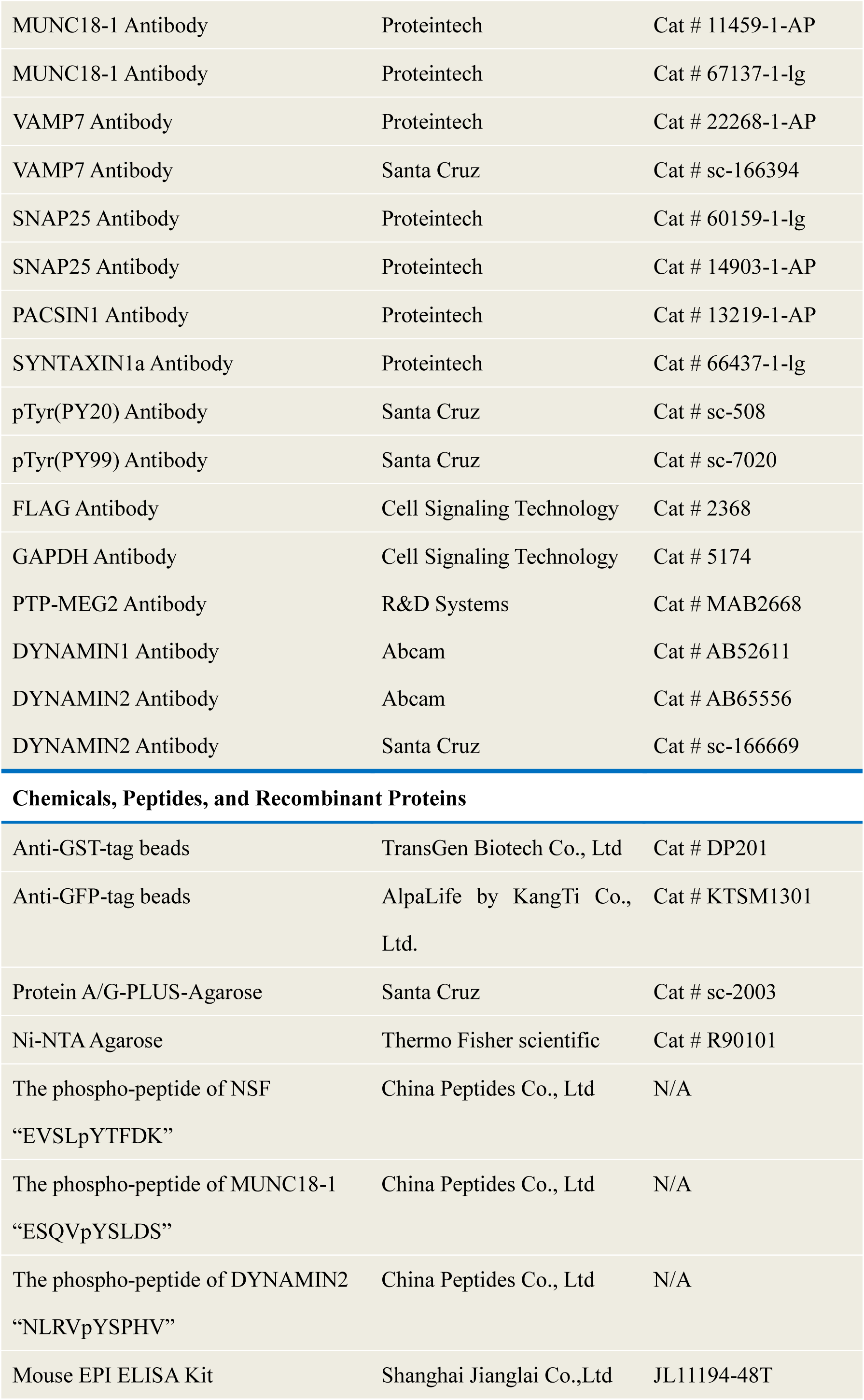
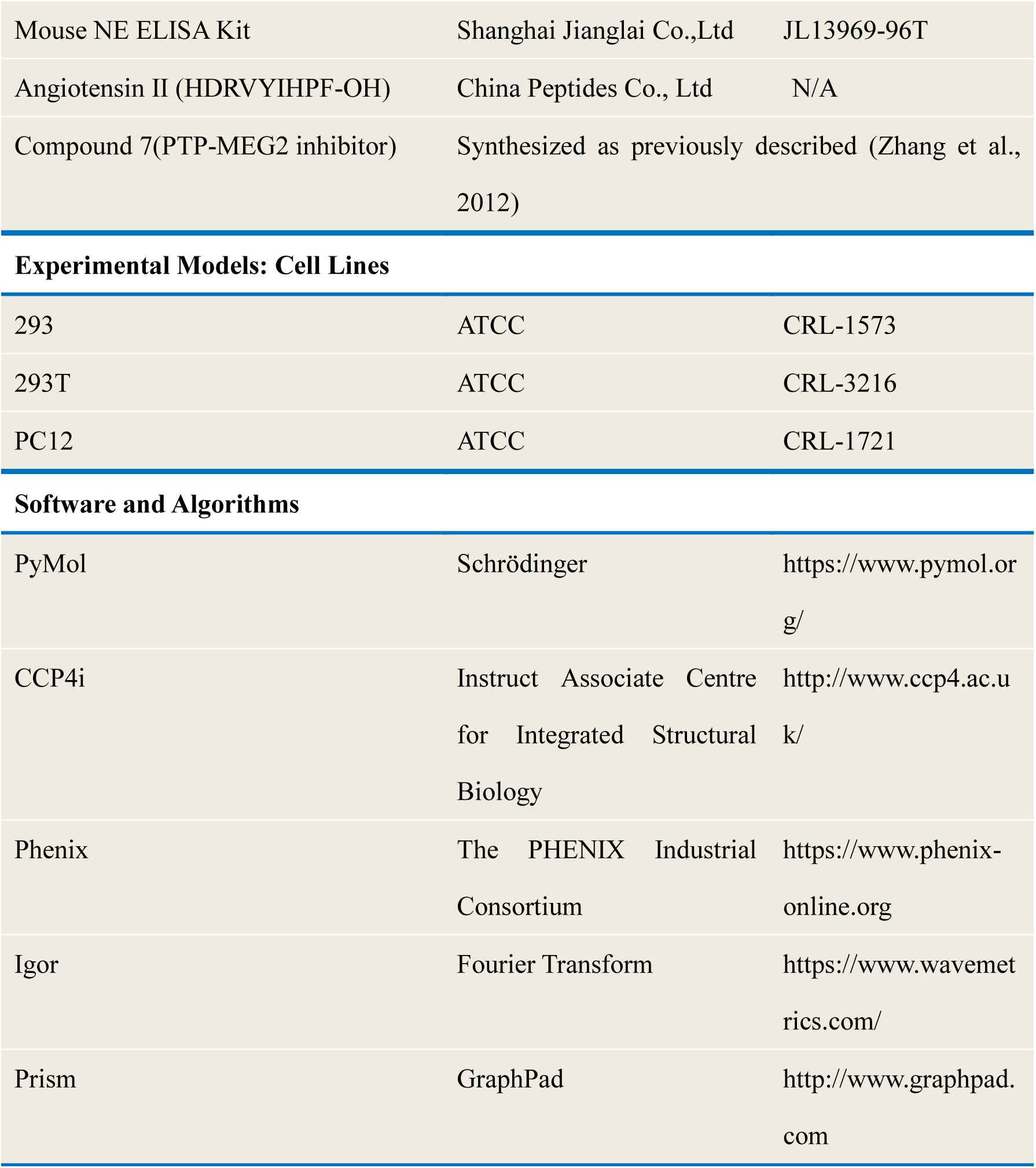

| REAGENT or RESOURCE | SOURCE | IDENTIFIER |
| --- | --- | --- |
| Antibodies |  |  |
| NSF Antibody | Proteintech | Cat # 21172-1-AP |
| NSF Antibody | Santa Cruz | Cat # sc-515043 |
| MUNC18-1 Antibody | Proteintech | Cat # 11459-1-AP |
| MUNC18-1 Antibody | Proteintech | Cat # 67137-1-lg |
| VAMP7 Antibody | Proteintech | Cat # 22268-1-AP |
| VAMP7 Antibody | Santa Cruz | Cat # sc-166394 |
| SNAP25 Antibody | Proteintech | Cat # 60159-1-lg |
| SNAP25 Antibody | Proteintech | Cat # 14903-1-AP |
| PACSIN1 Antibody | Proteintech | Cat # 13219-1-AP |
| SYNTAXIN1a Antibody | Proteintech | Cat # 66437-1-lg |
| pTyr(PY20) Antibody | Santa Cruz | Cat # sc-508 |
| pTyr(PY99) Antibody | Santa Cruz | Cat # sc-7020 |
| FLAG Antibody | Cell Signaling Technology | Cat # 2368 |
| GAPDH Antibody | Cell Signaling Technology | Cat # 5174 |
| PTP-MEG2 Antibody | R&D Systems | Cat # MAB2668 |
| DYNAMIN1 Antibody | Abcam | Cat # AB52611 |
| DYNAMIN2 Antibody | Abcam | Cat # AB65556 |
| DYNAMIN2 Antibody | Santa Cruz | Cat # sc-166669 |
| **Chemicals, Peptides, and Recombinant Proteins** |  |  |
| Anti-GST-tag beads | TransGen Biotech Co., Ltd | Cat # DP201 |
| Anti-GFP-tag beads | AlpaLife by KangTi Co., Ltd. | Cat # KTSM1301 |
| Protein A/G-PLUS-Agarose | Santa Cruz | Cat # sc-2003 |
| Ni-NTA Agarose | Thermo Fisher scientific | Cat # R90101 |
| The phospho-peptide of NSF<br>“EVSLpYTFDK” | China Peptides Co., Ltd | N/A |
| The phospho-peptide of MUNC18-1<br>“ESQVpYSLDS” | China Peptides Co., Ltd | N/A |
| The phospho-peptide of DYNAMIN2<br>“NLRVpYSPHV” | China Peptides Co., Ltd | N/A |
| Mouse EPI ELISA Kit | Shanghai Jianglai Co.,Ltd | JL11194-48T |
| Mouse NE ELISA Kit | Shanghai Jianglai Co.,Ltd | JL13969-96T |
| Angiotensin II (HDRVYIHPF-OH) | China Peptides Co., Ltd | N/A |
| Compound 7(PTP-MEG2 inhibitor) | Synthesized as previously described (Zhang et al., 2012) |  |
| Experimental Models: Cell Lines |  |  |
| 293 | ATCC | CRL-1573 |
| 293T | ATCC | CRL-3216 |
| PC12 | ATCC | CRL-1721 |
| Software and Algorithms |  |  |
| PyMol | Schrödinger | https://www.pymol.org/ |
| CCP4i | Instruct Associate Centre for Integrated Structural Biology | http://www.ccp4.ac.uk/ |
| Phenix | The PHENIX Industrial Consortium | https://www.phenix-online.org |
| Igor | Fourier Transform | https://www.wavemetrics.com/ |
| Prism | GraphPad | http://www.graphpad.com |

### CONTACT FOR REAGENT AND RESOURCE SHARING

Further information and requests for resources and reagents should be directed to and will be fulfilled by the Lead Contact, Professor Xiao Yu. or Jinpeng Sun

### EXPERIMENTAL MODEL AND SUBJECT DETAILS

#### Cell Culture

The HEK293 cell lines, the 293T cell lines and PC12 cell lines were originally obtained from the American Type Culture Collection (ATCC). The HEK293 cell lines and 293T cell lines were grown in DMEM with 10% FBS (Gibco, Grand Island, NY, US) and 1% penicillin/streptomycin at 37 °C. PC12 cells were maintained at 37 °C in DMEM medium containing 10% FBS (Gibco, US), 5% donor equine serum (Gibco, US) and 1% penicillin/streptomycin.

#### Constructs

Sequences of PTP-MEG2 catalytic domain were subcloned into PET-15b expression vector with an N-terminal His tag or PGEX-6P-2 expression vector containing an N-terminal GST tag. The mutations of PTP-MEG2 were constructed by the Quikchange kit from Stratagene.

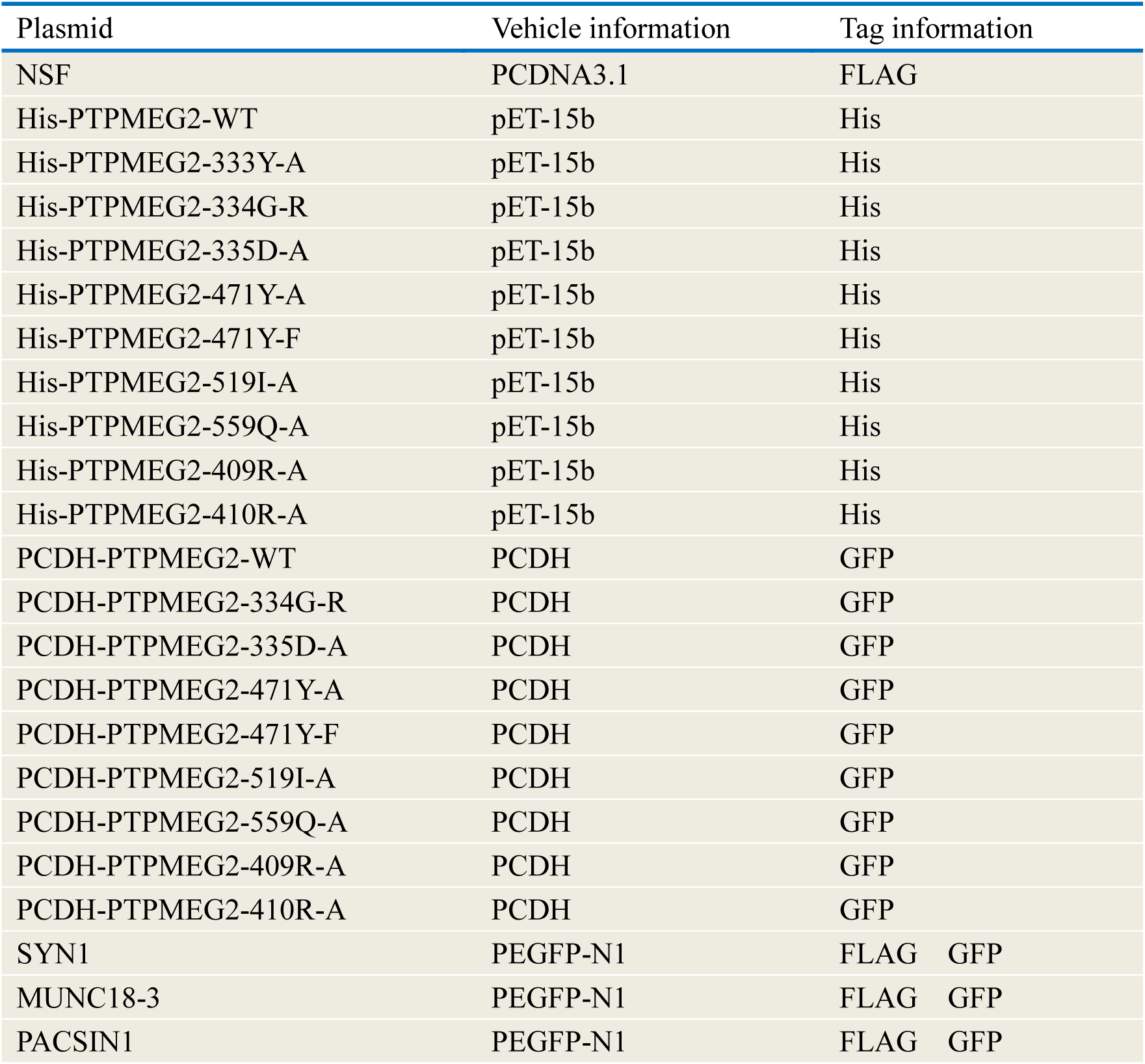
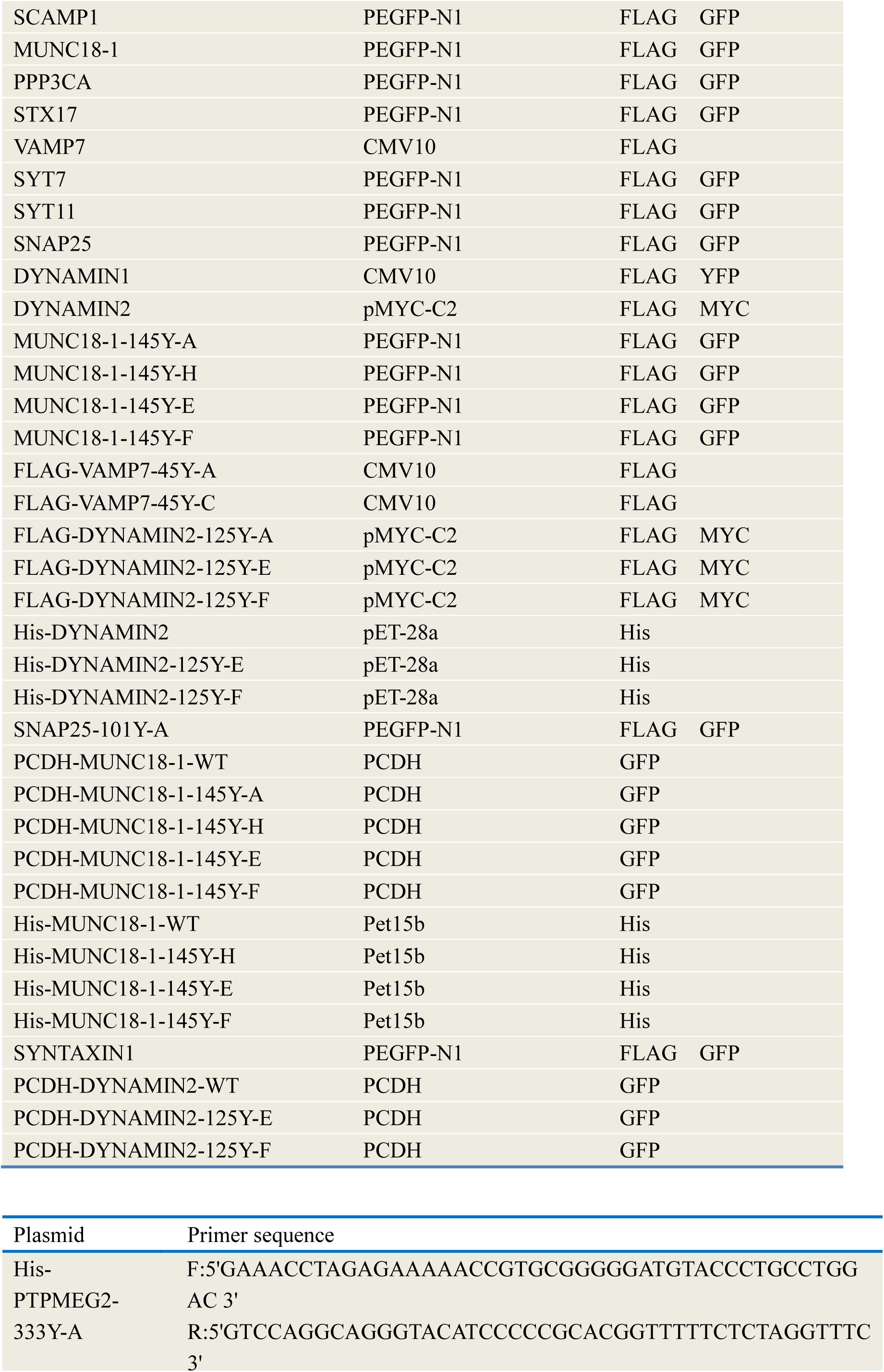
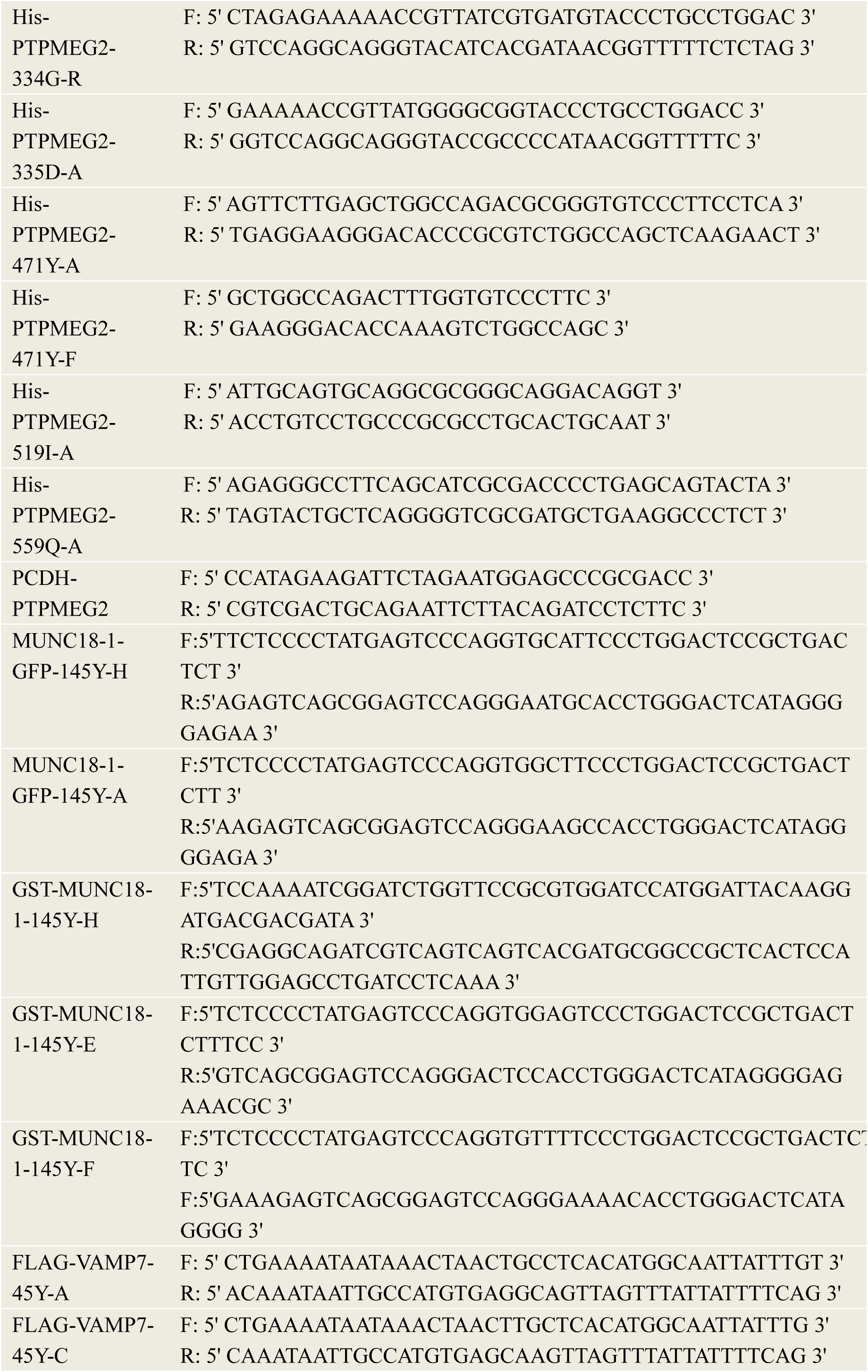
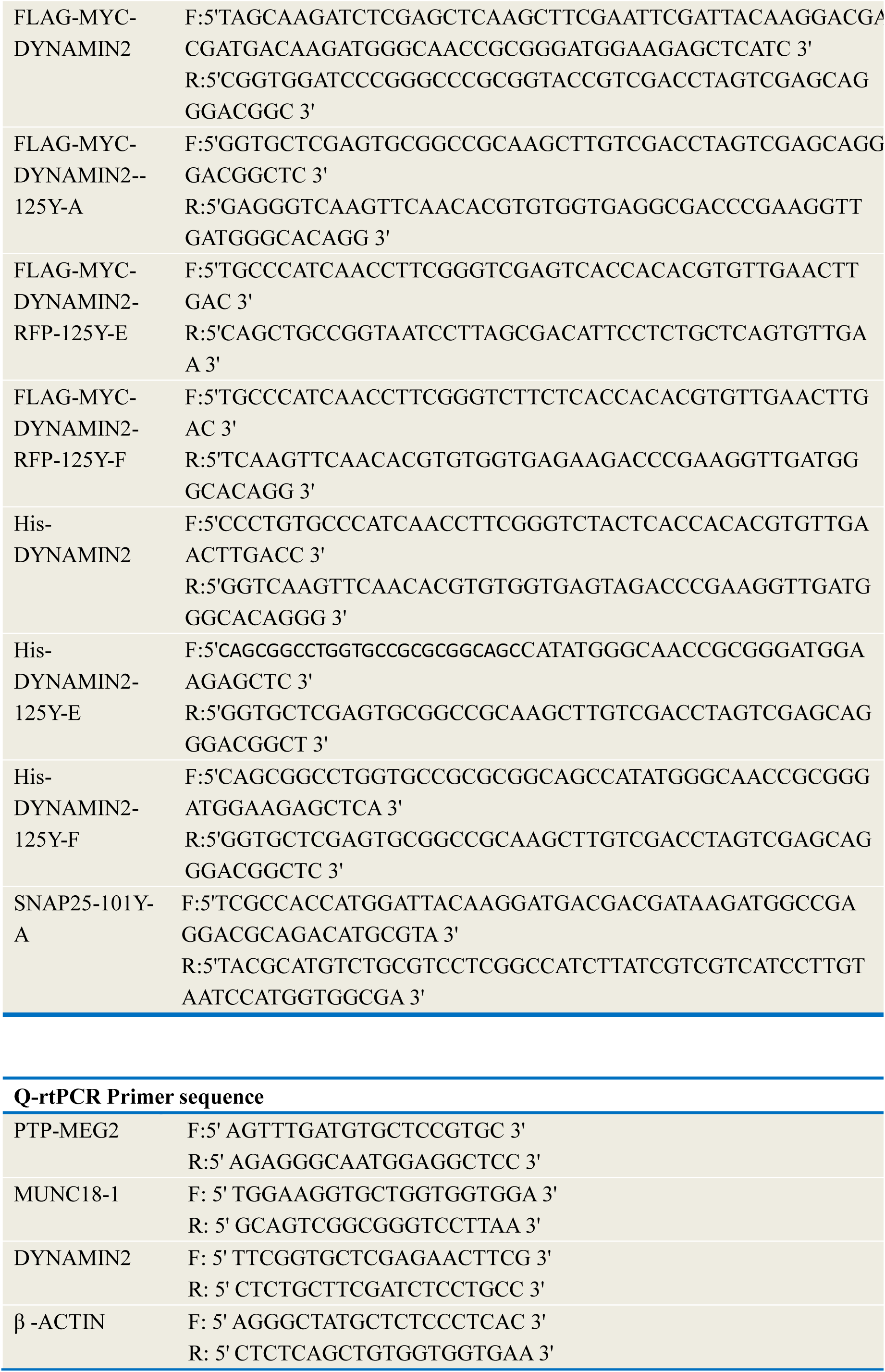

#### Recombinant lentivirus construction and lentivirus infection

For recombinant lentivirus packaging, the construction and infection was carried out as previously reported (Zhang et al., 2018). Plasmids carrying different genes including pCDH-PTP-MEG2-G^334^R/D^335^A/Y^471^A/Y^471^F/I^519^A/Q^559^A/ R^409^A/ R^410^A /WT-GFP, pCDH-MUNC18-1-Y^145^H/Y^145^E/Y^145^F/WT-GFP and pCDH-DYNAMIN2-Y^125^E/Y^125^F/WT-GFP were transfected into 293T cells using Lipofectamine TM 2000 (Thermo Fisher, Waltham, MA, USA) according to the manufacturer’s instructions. Three days after transfection, the supernatant of virus encoding PTP-MEG2, MUNC18-1 or DYNAMIN2 were collected and filtered. The PTP-MEG2, MUNC18-1 or DYNAMIN2 lentivirus (1×10 ^6^ TU/ml) was used to infect the primary chromaffin cells in later experiments.

#### ELISA

Freshly isolated adrenal medullas from adult female mice (6–8 weeks) were cultured in DMEM medium containing 1% penicillin/streptomycin and 10% FBS. After 2 hours starvation, adrenal medullas were stimulated with high KCl (70mM), AngII (100nM) for 1 min, with or without pre-incubation of a specific PTP-MEG2 inhibitor (400nM) for 1 hours. The supernatants were collected, and the epinephrine or norepinephrine secretion were determined by using the Epinephrine or norepinephrine ELISA kit (Shanghai Jianglai Co.,Ltd, JL11194-48T/JL13969-96T) according to the manufacture’s protocol.

#### Quantitative real-time PCR

Adrenal medullas were freshly isolated from adult female mice (6–8 weeks). The lentivirus encoding wild type or mutants of PTP-MEG2, MUNC18-1 or DYNAMIN2 (1×10 ^6^ TU/ml) were used to infect the primary chromaffin cells. On the third day, about 10 to 20 pheochromoffin cells with the different brightness were selected in each plate under the fluorescence microscope and the fluorescence photos were taken. After classification by different brightness, the cells were glued with the tips of ultra-fine glass tubes, and the tips of glass tubes with the cells were directly broken in the PCR tube. The PCR tube was preloaded with 5ul RNA enzyme inhibitors (Invitrogen, 10777019). 1ul DNA scavenger was added to the PCR tube, which was left at room temperature for half an hour and then terminated with a terminator. The purpose of this step is to remove the interference of genomic DNA. All the samples were reverse-transcribed using the Revertra Ace qPCR RT Kit (TOYOBO FSQ-101) to obtain cDNA. Then the quantitative real-time PCR was performed with the LightCycler apparatus (Bio-Rad) using the FastStart Universal SYBR Green Master (Roche). The expression of actin was used as a control.

#### Electrochemical amperometry

5-mm glass carbon fiber electrode (CFE) was used to measure quantal CA released from the mouse adrenal medulla chromaffin cell as previously described (Liu et al., 2017). We used a Multiclamp 700B amplifier (2012, Axon, Molecular Devices, USA) to perform electrochemical amperometry, which interfaced to Digidata 1440A with the pClamp 10.2 software (Liu et al., 2017). The holding potential of 780 mV was used to record the amperometric current (Iamp). All experiments were performed at room temperature (20–25°C). The CFE surface was positioned in contact with the membrane of clean primary chromaffin cells to monitor the quantal release of the hormone containing catecholamine substances. In our kinetic analysis of single amperometric spike, we only used the amperometric spikes with S/N > 3 (signal/noise). The standard external solution for our amperometry measurement is as follows: 5 mM KCl, 10 mM glucose, 10 mM HEPES pH 7.4, 2 mM CaCl_2_, 150 mM NaCl and 2 mM MgCl_2_. We analyzed all data using Igor (WaveMetrix, Lake Oswego, Oregon) and a custom-made macro program. Statistical data were given as the mean ± SEM and analyzed with one-way ANOVA.

#### Electron microscopy

The female mice (6–8 weeks) were decapitated, and the adrenal medullas were freshly isolated and cut to 150-μm-thick sections. The sections were immersed in Ringer’s saline (125mM NaCl, 2.5mM KCl, 1.25mM NaH_2_PO_4_, 26mM NaHCO_3_, 10mM D-Glucose, 2mM CaCl_2_, 1mM MgCl_2_) for 40 minutes at room temperature. During this period, continuous gases of 5% CO_2_ and 95% O_2_ were offered to the saline to ensure the survival of the tissue slice. After 40 minutes of starvation, the sections were stimulated with different conditions (control; only 100nM Ang II agonists for 1 min; only 400nM PTP-MEG2 inhibitor for 45 min; 100nM AngII agonists and 400nM PTP-MEG2 inhibitor for 1 min or 45 min) at 37°C respectively. These sections were firstly immersed in precooled 3% glutaraldehyde and fixed at 4°C for 2 hours, and then rinsed in PBS isotonic buffer, with repeated liquid exchanges and cleaning overnight, so that the samples were thoroughly rinsed and soaked in the buffer. After rinsing, the sample was fixed at 4°C with 1% osmium acid for 2 hours. It was rinsed with isosmotic buffer solution at 0.1M PBS for 15 minutes. The sections were dehydrated with ethanol at concentrations of 50%, 70%, 90%, then ethanol at concentration of 90% and acetone at concentration of 90%, at last only acetone at concentrations of 90%, 100%. We then replaced the acetone with the Epon gradually. The sections were added to Epon and polymerized at 60°C for 36 hours. Ultra -thin sections were performed at the thickness of 60nm by the LKB-1 ultra microtome, and then the ultra-thin sections were collected with the single-hole copper ring attached with formvar film. The sample was stained with 2% uranium acetate for 30 minutes, and then stained with 0.5% lead citrate for 15 minutes. These prepared samples were examined by JEM-1200EX electron microscope (Japan).

#### Western

Cells or medulla sections were lysed in lysis buffer (50 mM Tris pH 8.0, 150 mM NaCl, 1 mM NaF, 1% NP-40, 2mM EDTA, Tris-HCl pH 8.0, 10% glycerol, 0.25% sodium deoxycholate, 1mM Na_3_VO_4_, 0.3μM aprotinin, 130μM bestatin, 1μM leupeptin, 1μM repstatin and 0.5% IAA) after rinsing with the pre-chilled PBS on ice. Cell and tissue lysates were kept on ice for 35 min and then sediment via centrifugation for 15 min at 4 °C. The whole -cell and tissue protein lysates samples (30μg) were prepared for SDS-PAGE. The proteins in the gel were transferred to a nitrocellulose filter membrane by electro blotting, then probed with the appropriate primary and secondary antibodies. Antibody binding was detected by an HRP system.

#### Immunofluorescence

For the acquisition of tissue cells used in immunofluorescence, the mice were decapitated, and the adrenal medullas were freshly isolated (female mice, 6–8 weeks).The isolated adrenal medullas was immersed in 4% paraformaldehyde for fixation overnight at 4°C. Then the fixed tissues were washed for 4 hours in PBS containing 10% sucrose at 4°C for 8 hours in 20% sucrose, and in 30% sucrose overnight. Then these adrenal medullas were imbedded in Tissue-Tek OCT compound and then mounted and frozen them at −25°C. Subsequently, the adrenal medulla was cut to 4-μm-thick coronal serial sections. The adrenal medullas sections were blocked with 1% (vol/vol) donkey serum, 2.5% (wt/vol) BSA and 0.1% (vol/vol) Triton X-100 in PBS for 1.5 h. Then, the slides were incubated with primary antibodies against PTP-MEG2 (1:100), NSF (1:50), MUNC18-1 (1:50), VAMP7 (1:50), DYNAMIN2 (1:100), SNAP25 (1:100), PACSIN1 (1:50) at 4°C overnight. After washing with PBS for 3 times, the slides were incubated with the secondary antibody (1:500) for 1 h at room temperature. The slides were stained with DAPI (1:2000). Images were captured using a confocal microscope (ZEISS, LSM780). The Pearson’s co-localization coefficients were analyzed with Image-Pro Plus.

#### *K*_m_ and *k*_cat_ measurements

Enzymatic activity measurement was carried out as previously reported (Li et al., 2016; Wang et al., 2014). The standard solution (DMG buffer) for our enzymatic reactions is following: 50 mM 3, 3-dimethyl glutarate pH 7.0, 1 mM EDTA, 1 mM DTT. The ionic strength was maintained at 0.15 M (adjusted by NaCl). For the pNPP activity measurement, 100 µl reaction mixtures were set up in a total volume in a 96-well polystyrene plate (Thermo Fisher Scientific, Waltham, MA, US). The substrate concentration ranging from 0.2 to 5 *K*_m_ was used to determine the *k*_cat_ and *K*_m_ values. Reactions were started by the addition of an appropriate amount of His-PTP-MEG2-CD-WT or corresponding mutants, such as Y^333^A, G^334^R, D^335^A, Y^471^A, Y^471^F, I^519^A, Q559A, R^409^A, and R^410^A. The dephosphorylation of pNPP was terminated by adding 120µl 1M NaOH, and the enzymatic activity was monitored by measuring the absorbance at 405 nm. The activities toward phospho-peptide segment derived from NSF or MUNC18-1 were measured as following: In the first column, 90μl diluted NSF/MUNC18-1/DYNAMIN2 phospho-peptide substrate (100μM) was added. The successive columns were diluted by 1.5 times. The phospho-peptide substrates were preincubated at 37°C for 5 min. Reactions were started by the addition of an appropriate amount of enzymes. The dephosphorylation of NSF/MUNC18-1/DYNAMIN2 was terminated by adding 120µl Biomol green, and the enzymatic activities were monitored by measuring the absorbance at 620 nm. The steady-state kinetic parameters were determined from a direct fit of the data to the Michaelis-Menten equation using GraphPad Prism 5.0.

#### GST-pull down

To screen the candidate proteins interacting with PTP-MEG2, the GST beads were washed five times by cold binding buffer (20mM HEPES pH7.5, 1mM DTT, 1mM EDTA, and 100mM NaCl) and incubated with 5µg purified GST-PTP-MEG2-CD-D^470^A protein for 2 hours at 4°C. PC12 cells were transfected with FLAG-SYN1-GFP, FLAG-MUNC18-1-GFP, FLAG-MUNC18-1-Y^145^A-GFP, FLAG-MUNC18-1-Y^145^H-GFP, FLAG-MUNC18-1-Y^145^E-GFP, FLAG-MUNC18-1-Y^145^F-GFP, FLAG-MUNC18-3-GFP, FLAG-PACSIN1-GFP, FLAG-SCAMP1-GFP, FLAG-PPP3CA-GFP, FLAG-STX17-GFP, FLAG-VAMP7, FLAG-VAMP7-Y^45^A, FLAG-VAMP7-Y^45^C, FLAG-SYT7-GFP, FLAG-SYT11-GFP, FLAG-SNAP25-GFP, FLAG-SNAP25-Y^101^A-GFP, FLAG-DYNAMIN1-GFP, FLAG-DYNAMIN2, FLAG-DYNAMIN2-Y^125^A, FLAG-DYNAMIN2-Y^125^E, FLAG-DYNAMIN2-Y^125^F. After stimulation with 100nM AngII for 5 minutes at 37 °C, the cells were washed and then lysed in lysis buffer [20 mM HEPES pH 7.5, 100 mM NaCl, 0.5% NP-40, 5 mM iodoacetic acid, and a protease inhibitor mixture (final concentrations, 10μg of leupeptin, 1μg of aprotinin, 1μg of pepstatin, 1μg of antipain, and 20μg of phenylmethylsulfonyl fluoride per ml), 10mM DTT, 2mM EDTA] on ice for 30 minutes, then centrifuged at 12000rpm for 15 minutes at 4 °C. 20μl GST beads-PTP-MEG2-D^470^A protein was added into 500μl supernatants and the mixtures were subjected to end-to-end rotation at 4 °C for 2 hours. The GST beads and their binding proteins were washed five times with cold binding buffer to exclude the unspecific binding proteins. The pull-down experiment of MUNC18-1 and SYNTAXIN1 were described similar to above description. The GST beads were washed five times by cold binding buffer (described as above). After that, 5µg purified GST-MUNC18-1-WT or its Y^145^H, Y^145^E or Y^145^F mutant proteins were added into 20μl GST-agarose and incubated at for 4 °C 2 hours with end to end rotation. PC12 cells transfected with FLAG-SYNTAXIN1 were lysed in lysis buffer (described as above) and centrifuged to remove the pallets. The supernatants were added with 20μl GST beads/GST-fusion protein and the mixtures were subjected to end-to-end rotation at 4 °C for 2 hours. The FLAG-SYNTAXIN1 was detected with FLAG antibody.

#### GFP-pull down

PC12 cells were infected with lentiviruses encoding GFP-tagged PTP-MEG2-D^470^A of wild type and different mutant types. After stimulation with 100nM AngII for 5 minutes at 37 °C, the cells were washed and then lysed in lysis buffer [20 mM HEPES pH 7.5, 100 mM NaCl, 0.5% NP-40, 5 mM iodoacetic acid, and a protease inhibitor mixture (final concentrations, 10μg of leupeptin, 1μg of aprotinin, 1μg of pepstatin, 1μg of antipain, and 20μg of phenylmethylsulfonyl fluoride per ml), 10mM DTT, 2mM EDTA] on ice for 30 minutes, then centrifuged at 12000rpm for 15 minutes at 4 °C. The GFP beads were washed five times by cold binding buffer (20mM HEPES pH7.5, 1mM DTT, 1mM EDTA, and 100mM NaCl). 25μl anti-GFP-antibody conjugated beads were added into 500μl supernatants and the mixtures were subjected to end-to-end rotation at 4 °C for 1 hours. The PTP-MEG2-D^470^A-GFP proteins were pulled down and washed five times with cold binding buffer to exclude the unspecific binding proteins. The associated PTP-MEG2 substrates in the GFP-pull-down beads were then examined by western blot.

#### Immunoprecipitation and in vitro dephosphorylation

Rat adrenal medullas were isolated and cut into pieces in D-hanks buffer and stimulated with 70mM KCl or 100nM AngII for 5 minutes. The tissues were then lysed and grinded on ice in lysis buffer supplemented with proteinase inhibitor as described before. The lysates were centrifuged at 12000 rpm for 20 minutes at 4 °C and the pallets were removed. Before incubation with the lysates in the later step, the 1µg primary antibodies of NSF, MUNC18-1, VAMP7, SNAP25 or DYNAMIN2 were incubated with 20µl Protein A/G beads by end-to-end rotation overnight at 4 °C. Then the supernatants were incubated with primary antibody pre-coated with Protein A/G beads and rotated for 2 hours. 5µg purified PTP-MEG2-WT protein or control solution were added into the lysates for the in vitro dephosphorylation. A pan-phospho-tyrosine antibody pY^20^/ pY^99^ was used in Western blotting to detect the phosphorylated tyrosine of NSF/MUNC18-1/VAMP7/SNAP25/DYNAMIN2 in adrenal medullar with or without incubation with the PTP-MEG2.

#### Protein Expression and Purification

The wide type and mutant proteins of His-tagged PTP-MEG2-catalytic domain were expressed in BL21-DE3 Escherichia coli as previously described (Pan et al., 2013). In brief, 0.4 mM isopropyl1-thio-D-galactopyranoside (IPTG) was used to induce the expression of His-PTP-MEG2, and the bacteria lysates were centrifuged at 12000 rpm for 1 hour. Ni-NTA Agarose was applied to bind the His-tagged PTP-MEG2 and an imidazole gradation was used to elute the binding proteins. His-PTP-MEG2 was then purified with gel filtration chromatography to achieve at least 95% purity. The wide type and mutant proteins of GST-PTP-MEG2 and GST-MUNC18-1 were also expression in E.coli in presence of 0.4 mM IPTG for 16 hours at 25°C. After centrifugation and lysis, the proteins were purified by binding with GST-Sepharose for 2 hours and eluted by GSH.

#### Crystallization and Data Collection

For crystallization, His-PTP-MEG2-C^515^A/D^470^A protein (concentration at 15mg/ml) was mixed with NSF-pY^83^ peptide (EVSLpYTFDK) or MUNC18-1-pY^145^ peptide (ESQVpYSLDS) with molar ratio as 1:3 in buffer A (pH 7.2, 20 mM HEPES, 350 mM NaCl, and 1 mM DTT). 1μl mixed protein was blended with 1μl buffer B (pH 6.4, 20% PEG 4000, 0.2M KSCN, 10% ethylene glycol, 0.1M bis-tris propane) at 4°C for 3 days before crystals appears. The cubic crystals were preserved in liquid nitrogen very quickly dipped in storage buffer (buffer B supplemented with 10% glycerol). The data were collected at Shanghai Synchrotron Radiation Facility beamline BL17U1 using 0.98Å X-ray wavelength and analyzed by HKL2000.

#### Structural determination and refinement

The crystals of PTP-MEG2-NSF-pY^83^ and PTP-MEG2-MUNC18-1-pY^145^ peptides belong to the P2_1_2_1_2 space group. In each asymmetric unit, both PTPMEG2-NSF-pY^83^ and PTPMEG2-MUNC18-1-pY^145^ contain one monomer in one unit. Molecular replacement with Phaser in the CCP4 software package, with PTP-MEG2 catalytic domain (PDB code: 2PA5, water deleted) as the initial search model. Further refinements were carried out using the PHENIX program with iterative manual building in COOT. The data of the final refined structures are shown in Supplementary information Table 1.

#### BRET assay

The BRET experiment was performed to monitor the mutation effects of DYNAMIN2 on AT1aR endocytosis in response to AngII (1μM) stimulation as previously described (Liu et al., 2017). Plasmids encoding DYNAMIN2 (DYNAMIN2-WT, DYNAMIN2-Y^125^E or DYNAMIN2-Y^125^F) and LYN-YFP, AT1aR-C-RLUC were co-transferred into HEK293 cells at 1:1:1 ratio for 48 h. After 12 h of starvation, cells were digested with trypsin and harvested. Then the transfected cells were washed with PBS at least three times and evenly distributed into a 96-well plate (Corning Costar Cat. # 3912). Cells were stimulated with AngII (1μM) for 25 min at 37 °C. Afterwards, Coelenterazine h was incubated with the cells at 25 °C (Promega S2011, final concentration, 5μM). We used two different light emissions for BRET measurements (530/20 nm for yellow fluorescent protein and 480/20 nm for luciferase).

#### GTPase activity of DYNAMIN2

To prepare His-tagged DYNAMIN2-WT, DYNAMIN2-Y^125^E, DYNAMIN2-Y^125^F (His-DYNAMIN2-WT, His-DYNAMIN2-Y^125^E, His-DYNAMIN2-Y^125^F), DNA fragments were generated by PCR using MYC-DYNAMIN2 as the template and subcloned into pET-28a (+) plasmid. The resulting plasmids were transformed into bacterial BL21-DE3-RIPL cells for protein expression. The DYNAMIN proteins were then purified by affinity chromatography. We measured the DYNAMIN2 GTPase activity by using GTPase-Glo assay test kit (V7681) from Promega Corporation according to its instructions.

#### Bioinformatic search of PTP-protein interactions

We identified the PTP-protein interactions by two independent bioinformatic analyses. On one site, we extracted the potential PTP-interacting proteins from the STRING database (Szklarczyk et al., 2015) by setting the parameters of Homo sapiens. On the other site, we used Pubtator (Wei et al., 2013) to text-mine potential PTP-protein associations from PubMed literature (by 2018.9) via the search of combinational keywords such as “tyrosine phosphorylation mutation, vesicular, fusion, and human”. We compared these two protein lists to produce a consensus PTP-interacting protein list. Subsequently, we narrowed down the protein list by satisfying following constraints: (a) The protein should express in the adrenal gland as documented in the TissueAtlas (Thul et al., 2017); (b) The protein should expose tyrosine residue(s) on surface for phosphorylation as simulated by the Molecular Operating Environment (MOE) (Vilar et al., 2008); (c) On the protein, at least one predicted tyrosine phosphorylation site predicted by the PhosphoSitePlus (Hornbeck et al., 2015) is also experiment-validated. The refined PTP-protein pairs were then ready for later experimental analyses.

#### LC-MS/MS analysis

Rat adrenal medullas were isolated and cut into pieces in D-hanks buffer and stimulated with 100nM AngII for 5 minutes. The tissues were then lysed and grinded on ice in lysis buffer supplemented with protease inhibitors. The lysates were centrifuged at 12000 rpm for 20 minutes at 4 °C and the pallets were removed. Before incubation with the lysates in the later step, the GST beads were washed five times by cold binding buffer (20mM HEPES pH7.5, 1mM DTT, 1mM EDTA, and 100mM NaCl) and incubated with 5µg purified GST-PTP-MEG2-CD-D^470^A protein for 2 hours at 4°C. 20μl GST beads-PTP-MEG2-D^470^A protein was added into 500μl supernatants and the mixtures were subjected to end-to-end rotation at 4 °C for 2 hours. The GST beads and their binding proteins were washed five times with cold binding buffer to exclude the unspecific binding proteins. Denatured proteins were separated by 10% SDS–PAGE and subjected to trypsin digestion. Phosphopeptides were analyzed by the LTQ Orbitrap Elite (Thermo Scientific, Beijing Qinglian Biotech Co., Ltd). Spectra were analyzed by Proteome Discoverer software, and phosphorylation sites were confirmed manually.

#### Protein degradation assay

HEK293 cell were seeded in 6-well plates. Plasmids encoding MUNC18-1 (MUNC18-1-WT, MUNC18-1-Y^145^E or MUNC18-1-Y^145^F) or DYNAMIN2 (DYNAMIN2-WT, DYNAMIN2-Y^125^E or DYNAMIN2-Y^125^F) was transferred into HEK293 cells for 24 h. Cells were treated with cycloheximide (CHX 10μg/ml) and harvested at different time points (0, 4, 8, 12h) for western blot analysis.

#### Statistical analysis

All data are presented as mean ± SEM. All data were analyzed using two-tailed Student’s t-test or one-way ANOVA. All of the Western films were scanned, and band intensity was quantified with ImageJ software (National Institutes of Health, Bethesda MD). P < 0.05 was considered as statistically significant.

## DATA AVAILABILITY

The coordinates and density map for the PTP-MEG2-NSF-pY^83^ (PDB ID: 6KZQ) and PTP-MEG2-MUNC18-1-pY^145^ (PDB ID: 6L03) peptides has been deposited in protein data bank. All other data are available upon request to the corresponding authors.

## ACKNOWLEDGEMENTS

We thank Dr Michael Xi Zhu for stimulating discussions and critical reading of the manuscript. We thank Yanmei Lu from Shandong jiaotong hospital, for her help with Transmission electron microscopy analysis. We thank Daolai Zhang and Mingliang Ma for their technical assistance in lentivirus packaging. We thank Yujing Sun and Zhixin Liu for their technical assistance in electrochemical recording. We acknowledge support from the National Key Basic Research Program of China Grant 2018YFC1003600 (to X.Y. and J.-P.S.), the National Natural Science Foundation of China Grant 81773704 (to J.-P.S.) and Grant 81601668 (to Y.-F.X.), the National Science Fund for Distinguished Young Scholars Grant 81825022 (to J.-P.S.), the National Science Fund for Excellent Young Scholars Grant 81822008 (to X. Y.) and the Rolling program of ChangJiang Scholars and Innovative Research Team in University Grant IRT_17R68 (to Y. S.). S.Z. and Z.-Y. Z. are supported by NIH RO1 CA69202.

## AUTHOR CONTRIBUTIONS

J.-P.S. and X.Y. conceived the whole research and initiated the project. J.-P.S., X.Y., Y.-F.X. and X.C. designed all the experiments. J.-P.S. and X.Y. supervised the overall project design and execution. X.C., Y.-F.X., Z.Y., X.Y. and J.-P.S. participated in data analysis and interpretation. C.-H.L. and Y.-J.W. performed electrochemical experiments. Z.Y., P.X., Z.-L.Z. helped us collect crystal data and analyze crystal structure. M.C., K.-S.L., Y.-C.S. performed cell biology, molecular biology and biochemistry experiments. Z.-G.X, provided DYNAMIN2 plasmid. Z.-Y.Z. and S.Z. synthesized and purified Compound 7 (PTP-MEG2 inhibitor). X.-Z.Y. and Z.-L.J. performed substrate bioinformatics screening. Z.-Y.Z., W.-D.Z., C.-H.W. and C.W. provided insightful idea and experimental designs. J.-P.S, Y.-F.X., X.C. and X.Y. wrote the manuscript. All of the authors have seen and commented on the manuscript.

## ABBREVIATIONS

PTP: protein tyrosine phosphatase
NSF: N-ethylmaleimide-sensitive fusion protein
AngII: Angiotensin II
EPI: epinephrine
NE: norepinephrine
CFE: carbon fiber electrode
LDCVs: large-dense-core vesicles
PSF: pre-spike foot
SAF: stand-alone foot
pNPP: p-nitrophenyl phosphate
MUNC18: Mammalian homolog of Unc-18
Cav1.2: Voltage-gated calcium channel subunit alpha Cav1.2
VAMP7: Vesicle-associated membrane protein 7
PPP3CA: Serine/threonine-protein phosphatase 2B catalytic subunit alpha isoform
PM: plasma membrane
LYP: Lymphoid phosphatase
SKAP-HOM: Src kinase-associated phosphoprotein
HER2: Receptor tyrosine-protein kinase erbB-2
SHP1: Protein-tyrosine phosphatase SHP-1
PTPH1: Protein-tyrosine phosphatase H1.

