## Supplementary figure for "PTP-MEG2 regulates quantal size and fusion pore opening through two distinct structural bases and substrates"

Yun-Fei Xu et al.

### Supplemental figure 1

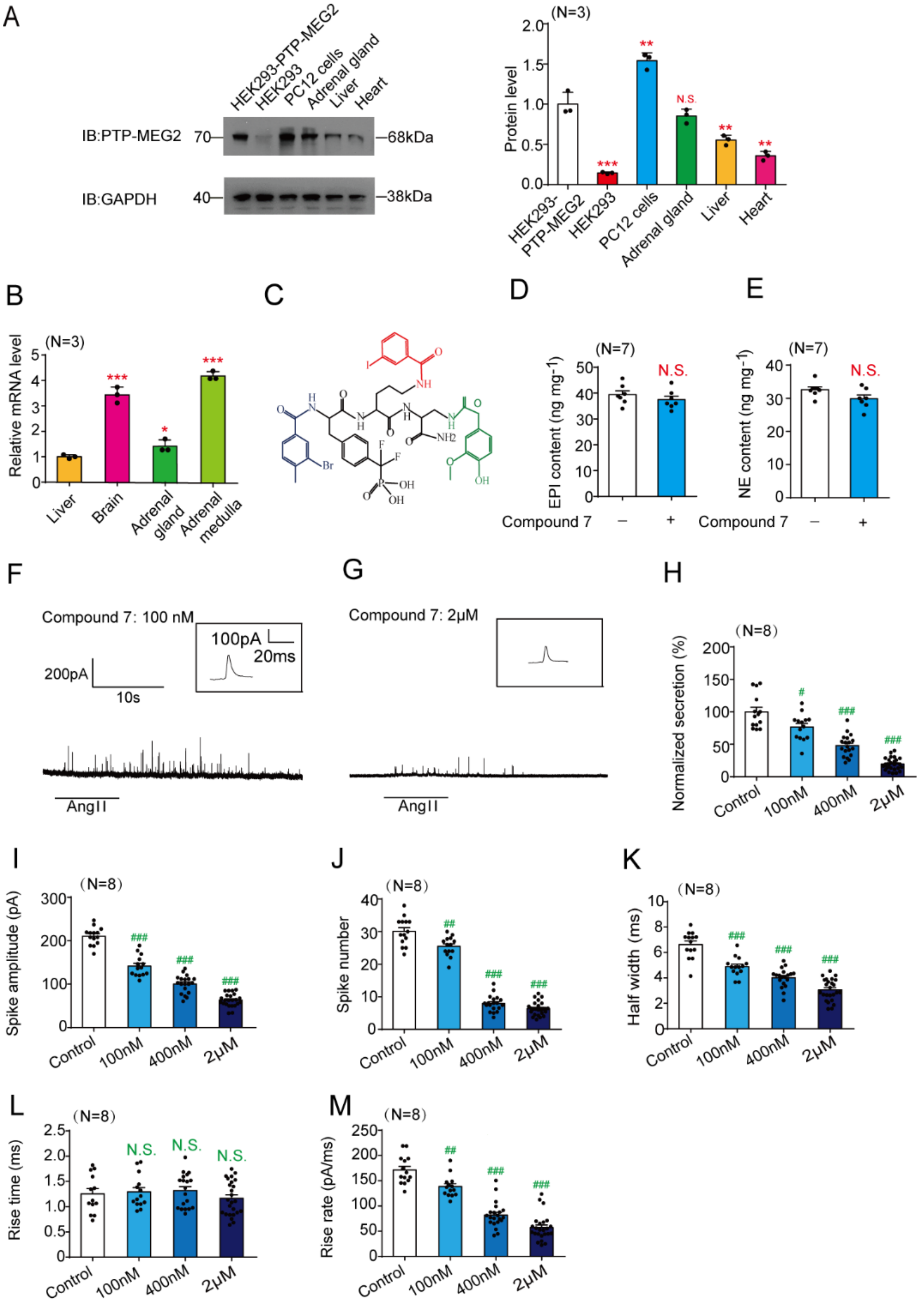

**Supplemental Figure 1. Inhibition of PTP-MEG2 impairs epinephrine and norepinephrine secretion.**

(A). Expression of PTP-MEG2 in different cell lines and tissues by western blot. Left panel: Representative western blot from at least three independent experiments was shown for the PTP-MEG2 expression in PTP-MEG2 transfected HEK-293 cells, control HEK-293 cells, PC12 cells, mouse adrenal gland, liver and heart, with GAPDH as an internal control. Right panel: Bar graph representation of the expression of PTP-MEG2 in different cell lines and tissues which were quantified from three independent experiments.

(B). PTP-MEG2 mRNA levels in mouse liver, brain, adrenal gland, and adrenal medulla were determined by quantitative reverse transcription PCR (Q-RTPCR). The differences of cycle threshold values ($C_T$) between the samples ($\Delta C_T$) were calculated after ratio to control GAPDH and normalized with liver expression. Data were obtained from 3 independent experiments and displayed with mean ± SEM.

(C). Schematic representation of structure of the specific PTP-MEG2 inhibitor used in the manuscript.

(D-E). Total amount of EPI (D) and NE (E) in isolated adrenal medulla were detected with ELISA with or without Compound 7 (400 nM) incubation for 2 hours. PTP-MEG2 inhibitor Compound 7 had no significant effects on total amounts of EPI or NE contents in adrenal medulla. N.S means not significant difference.

(F-G). AngiotensinII (AngII) at concentration of 100nM induced typical amperometric spikes of primary mouse chromaffin cells after incubation with PTP-MEG2 inhibitor at different concentration of 100nM (F) or 2μM (G).

(H).Statistical diagram of secretory amount of catecholamine from cells incubated with different doses of PTP-MEG2 inhibitor. The average secretory amount of cells without inhibitor was set as 100% and secretory amounts of other subgroups were normalized by ratio to control group.

(I-M). Quantitative analysis of the amperometric spikes stimulated by AngII (100nM) with or without PTP-MEG2 inhibitor at different concentrations, including the spike amplitude (I), spike number (J), half width (K), rise time (L) and rise rate (M). A total of 507 amperometric spikes were analysed from 72 chromaffin cells.

(A), (B), (D), (E), and from (H) to (M): * in (A) indicates protein expression level of PTP-MEG2 in different tissues or cells compared with overexpression of PTP-MEG2 in HEK293 cells. * in (B) indicates mRNA expression level of PTP-MEG2 in different tissues or cells compared with overexpression of PTP-MEG2 in HEK293 cells. # indicates the PTP-MEG2 inhibitor group compared with the control group. *, $P<0.05$; **, $P<0.01$; *** $P<0.001$ and #, $P<0.05$; ##, $P<0.01$; ### $P<0.001$. (D), (E), the data were analysed using Student's t-test. (A), (B) and from (H) to (M), the data were analysed using one-way ANOVA.

### Supplemental figure 2

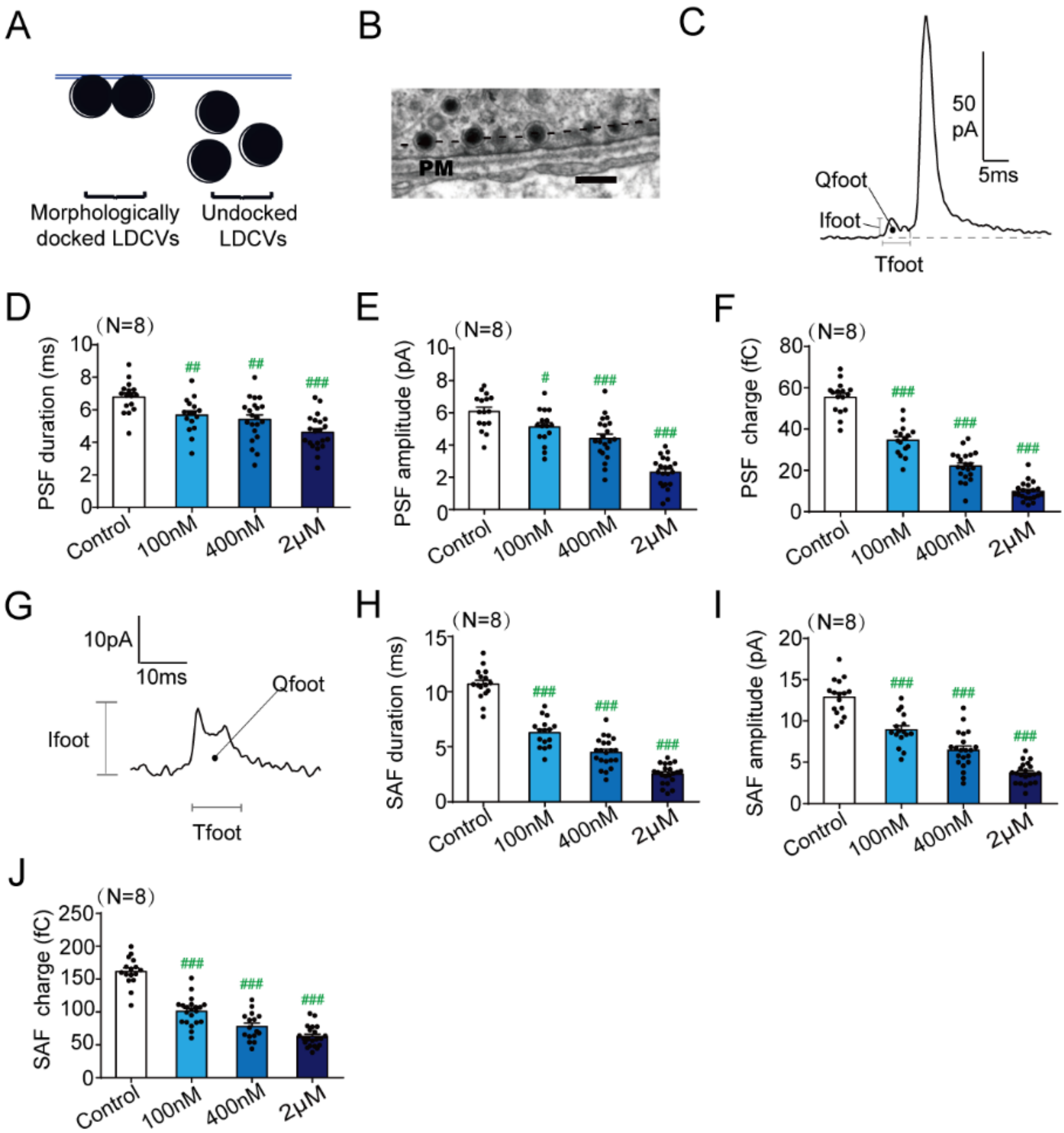

**Supplemental Figure 2. Effects of PTP-MEG2 inhibitor on the docking vesicles and PSF/SAF.**

(A). Schematic image of the manifestations of morphological docking and undocked large dense core vesicles (LDCV).

(B). Representative image of morphological docking vesicles. Vesicles with distance less than 50nm (dashed line) from plasma membrane were defined as morphological docking vesicles. Scale bars: 150 nm.

(C). Representation of the pre-spike foot (PSF) by a typical spike in our experiments. T-foot indicated the duration of a PSF, the Ifoot indicated the amplititude of a PSF and the Qfoot indicated the total charge of the PSF.

(D-F). Different parameters of pre-spike foot including the PSF duration (Tfoot) (D), PSF amplititude (Ifoot) (E), PSF charge (Qfoot) (F) were calculated from 72 chromaffin cells.

(G). Representation of the stand alone foot (SAF) in our experiments.

(H-J). Different parameters of stand alone foot (SAF) of cells incubated with different dose of PTP-MEG2 inhibitor were calculated, including duration（Tfoot）(H), amplitude (Ifoot) (I), and the average total charge (Qfoot) (J). D-F, H-J, # indicates the PTP-MEG2 inhibitor group compared with the control vehicle group. #, $P<0.05$; ##, $P<0.01$; ### $P<0.001$; N.S. means no significant difference. All the data were analysed using one-way ANOVA.

Supplemental figure 3

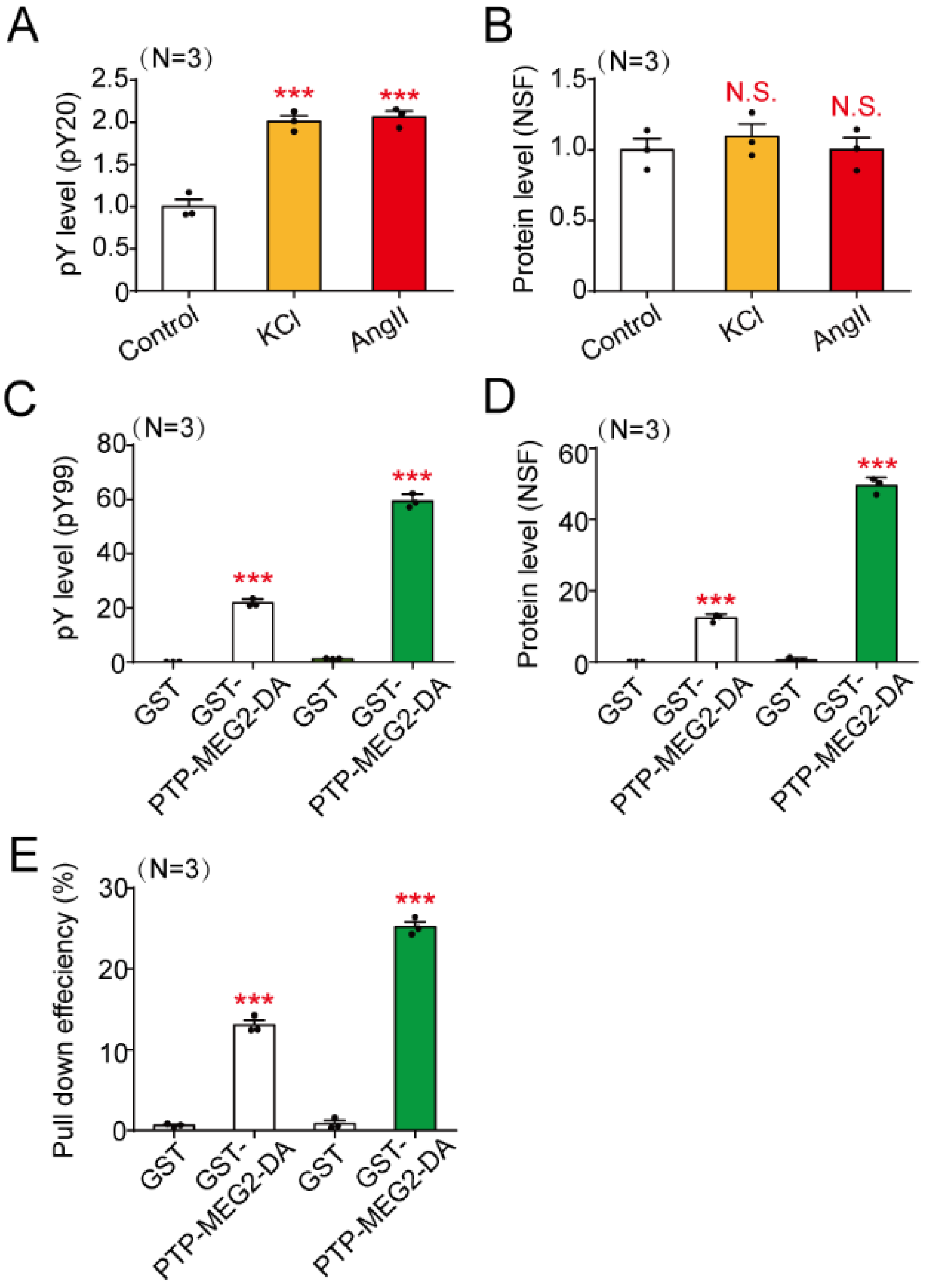

**Supplemental Figure 3. PTP-MEG2 interacts with and dephosphorylates tyrosine phosphorylated NSF in chromaffin cells.**

(A-B). Western blots of the phosphorylated NSF (A) and total immunoprecipitated NSF (B) in Figure 2A were quantified. *** represented $P<0.001$ of KCl or AngII stimulated cells compared with control groups without stimulation. N.S. means no significant difference.

(C-D). $pY^{99}$ (C) and NSF (D) levels in Figure 2D were quantified.

(E) The pull-down efficiency of GST-MEG2-DA on NSF in Figure 2D was calculated by NSF (amount of output)/ NSF(amount of input)*100%.

*** represented $P<0.001$ compared with GST group. All the data were analysed using one-way ANOVA.

### Supplemental figure 4

**A** pNPP

| Enzymes | $k_{cat}$ [ $s^{-1}$ ] | $K_m$ [ mM ] | $k_{cat}/K_m$ [ $10^3$, $M^{-1}s^{-1}$ ] | Fold of Activity Decrease |
|---|---|---|---|---|
| WT | 6.65±0.13 | 10.9±0.4 | 0.61±0.01 | 1 |
| Y333A | 1.39±0.05 | 23.3±1 | 0.06±0.01 | 10.14 |
| G334R | 4.76±0.24 | 6.9±1 | 0.69±0.07 | 0.88 |
| D335A | 3.47±0.28 | 5.1±0.6 | 0.68±0.03 | 0.89 |
| Y471A | 1.43±0.04 | 2.6±0.7 | 0.55±0.13 | 1.11 |
| Y471F | 2.70±0.05 | 5.4±0.6 | 0.50±0.05 | 1.21 |
| I519A | 2.55±0.03 | 15±0.8 | 0.17±0.01 | 3.53 |
| Q559A | 0.49±0.01 | 1.1±0.1 | 0.45±0.03 | 1.35 |

**B** NSF

| Enzymes | $k_{cat}/K_m$ [ $10^5$, $M^{-1}s^{-1}$ ] | Fold of Activity Decrease | Selectivity for Peptide |
|---|---|---|---|
| WT | 85.2±2.5 | 1 | 1 |
| Y333A | 0.20±0.03 | 436.79 | 43 |
| G334R | 14.1±1.5 | 6.04 | 6.86 |
| D335A | 8.15±0.14 | 10.45 | 11.74 |
| Y471A | 5.59±0.17 | 15.24 | 13.73 |
| Y471F | 3.69±0.12 | 23.09 | 19.08 |
| I519A | 0.15±0.01 | 550.32 | 155.89 |
| Q559A | 0.04±0.01 | 2027.4 | 1501.8 |

**Supplemental Figure 4. Catalytic activity of PTP-MEG2-WT and different mutants toward pNPP and pY$^{83}$-NSF.**

(A). Parameters of the catalytic activity of PTP-MEG2-WT and different mutants toward the pNPP, including $k_{cat}$, $K_m$, and $k_{cat}/K_m$.

(B). $k_{cat}/K_m$ of PTP-MEG2-WT and different mutants towards the pY$^{83}$-NSF phospho-peptide.

The assays were performed with DMG buffer: 50 mM 3,3-dimethyl glutarate pH 7.0, 1 mM EDTA, 1 mM DTT. The ionic strength was maintained at 0.15 M (adjusted by NaCl). The enzymatic reactions were carried out at room temperature.

### Supplemental figure 5

| | pY loop 331–338 | β3-β4 loop 393–411 | WPD loop 466–472 | P loop 513–523 | Q loop 554–564 |
|---|---|---|---|---|---|
| hPTP-MEG2 | NRYGDVPC | QKVLVIVMTTRFEEGGRRK | LSWPDYG | VHCSAGIGRTG | RAFSIQTPEQY |
| bPTP-MEG2 | NRYGDVPC | QKVLVIVMTTRFEEGGRRK | LSWPDYG | VHCSAGIGRTG | RAFSIQTPEQY |
| rPTP-MEG2 | NRYGDVPC | QKVLVIVMTTRFEEGGRRK | LSWPDYG | VHCSAGIGRTG | RAFSIQTPEQY |
| mPTP-MEG2 | NRYGDVPC | QKVLVIVMTTRFEEGGRRK | LSWPDYG | VHCSAGIGRTG | RAFSIQTPEQY |
| hPTP-MEG1 | NRYRDISP | QGSSMVVMLTTQVERGRVK | IAWPDHG | VHCSAGIGRTG | RAMMIQTPSQY |
| hPTP1B | NRYRDVSP | QKSRGVVMLNRVMEKGSLK | TTWPDFG | VHCSAGIGRSG | RMGLIQTADQL |
| hTCPTP | NRYRDVSP | QKTKAVVMLNRIVEKESVK | TTWPDFG | IHCSAGIGRSG | RMGLIQTPDQL |
| hPTPH1 | NRYKDVLP | QKLSLIVMLTTLTERGRTK | VAWPDHG | VHCSAGIGRTG | RAMMVQTSSQY |
| hSTEP | NRYKTILP | EHTPIIVMITNIEE-MNEK | TSWPDQK | VHCSAGIGRTG | RGGMIQTCEQY |
| hSHP-1 | NRYKNILP | ENSRVIVMTTREVEKGRNK | LSWPDHG | VHCSAGIGRTG | RSGMVQTEAQY |
| hSHP-2 | NRYKNILP | ENSRVIVMTTKEVERGKSK | RTWPDHG | VHCSAGIGRTG | RSGMVQTEAQY |
| hHePTP | DRYKTILP | EEVSLIVMLTQLRE-GKEK | SAWPDHQ | VHCSAGIGRTG | RGGMIQTAEQY |
| hPEST | NRYKDILP | YNVVIIVMACREFEMGRKK | VNWPDHD | IHCSAGCGRTG | RHSAVQTKEQY |
| hLYP | NRYKDILP | YSVLIIVMACMEYEMGKKK | KNWPDHD | IHCSAGCGRTG | RPSLVQTQEQY |
| hBDP1 | NRYKDVLP | FGVKVILMACREIENGRKR | MSWPDRG | VHCSAGCGRTG | RPAAVQTEEQY |
| hPTPD1 | NRFQDVLP | QGIAIIAMVTAEEEGGREK | TDWPEHG | VHCSAGVGRTG | RMMLVQTLCQY |
| hPTPD2 | SRIREVVP | QGVNVIAMVTAEEEGGRTK | TDWPDHG | VHCSAGVGRTG | RMFMIQTIAQY |
| hPTPBAS | NRYKNILP | QKSTVIAMMTQEVEGEKIK | TAWPDHD | THCSAGIGRSG | RHGMVQTEDQY |
| hPTPTyp | NRYRDILP | NNCNVIAMITREIECGVIK | TKWPDHG | VHCSAGVGRTG | RCGMIQTKEQY |
| hHDPTP | NRHQDVMP | QKVSVIVMLVSEAEMEKQK | PTWPELG | VHCSSGVGRTG | RKHMLQEKLHL |
| hRPTPalpha | NRYVNILP | QNTATIVMVTNLKERKECK | TSWPDFG | VHCSAGVGRTG | RCQMVQTDMQY |
| hRPTPepsilon | NRYPNILP | QKSATIVMLTNLKERKEEK | TSWPDFG | VHCSAGVGRTG | RPQMVQTDMQY |
| hRPTPkappa | NRYGNIIA | EQSACIVMVTNLVEVGRVK | TGWPDHG | VHCSAGAGRTG | RINMVQTEEQY |
| hRPTPmu | NRYGNIIA | ENTASIIMVTNLVEVGRVK | TGWPDHG | VHCSAGAGRTG | RVNMVQTEEQY |
| hRPTPrho | NRYGNIIS | ENSASIVMVTNLVEVGRVK | TSWPDHG | VHCSAGAGRTG | RVNLVQTEEQY |
| hRPTPdelta | NRYANVIA | QRSATVVMMTKLEERSRVK | TAWPDHG | VHCSAGVGRTG | RNYMVQTEDQY |
| hRPTPsigma | NRYANVIA | QRSATIVMMTRLEEKSRIK | TAWPDHG | VHCSAGVGRTG | RNYMVQTEDQY |
| hRPTPgamma | NRYINILA | NLVEKGRRKCDQYWPTENS | TQWPDMG | VHCSAGVGRTG | RNYLVQTEEQY |
| hRPTPzeta | NRYINIVA | HNVEVIVMITNLVEKGRRK | TQWPDMG | VHCSAGVGRTG | RNYLVQTEEQY |
| hLAR | NRYANVIA | QRTATVVMMTRLEEKSRVK | MAWPDHG | VHCSAGVGRTG | RNYMVQTEDQY |
| hCD45 | NRYVDILP | VQKATVIVMVTRCEEGNRN | TSWPDHG | VHCSAGVGRTG | RCLMVQVEAQY |
| hGlepp1 | NRYTNILP | QKSQIIVMLTQCNEKRRVK | TAWPDHG | IHCSAGVGRTG | RMSMVQTEEQY |
| hDEP-1 | NRYNNVLP | KNVYAIIMLTKCVEQGRTK | TSWPDHG | VHCSAGVGRTG | RPLMVQTEDQY |
| hRPTPbeta | NRYNNILP | QNVHNIVMVTQCVEKGRVK | TVWPDHG | VHCSAGVGRTG | RVHMVQTECQY |
| hSAP-1 | NRYRNVLP | QQSHTLVMLTNCMEAGRVK | QAWPDHG | VHCSAGVGRTG | RPLMVQTEAQY |
| hPCPTP1 | NRYKTILP | KLKE-KNEKCVLYWPEK-- | TSWPDHK | VHCSAGIGRTG | RGGMVQTSEQY |
| hIA2 | NRHPDFLP | SGCTVIVMLTPLVEDGVKQ | LSWPAEG | VHCSDGAGRTG | RPGLVRSKDQF |
| hIA-2beta | NRSLAVLT | SGCVVIVMLTPLAENGVRQ | LSWYDRG | VHCSDGAGRSG | RPGMVQTKEQF |

**Supplemental Figure 5. Sequence alignment of the PTP-MEG2.**

The sequence alignment of the pY loop, β3-β4 loop, WPD loop, P loop, Q loop from different PTPs. Key residues involved in the interactions between PTP-MEG2 and NSF, as well as the PTP-MEG2 and MUNC18-1 are highlighted and compared with other PTP members.

### Supplemental figure 6

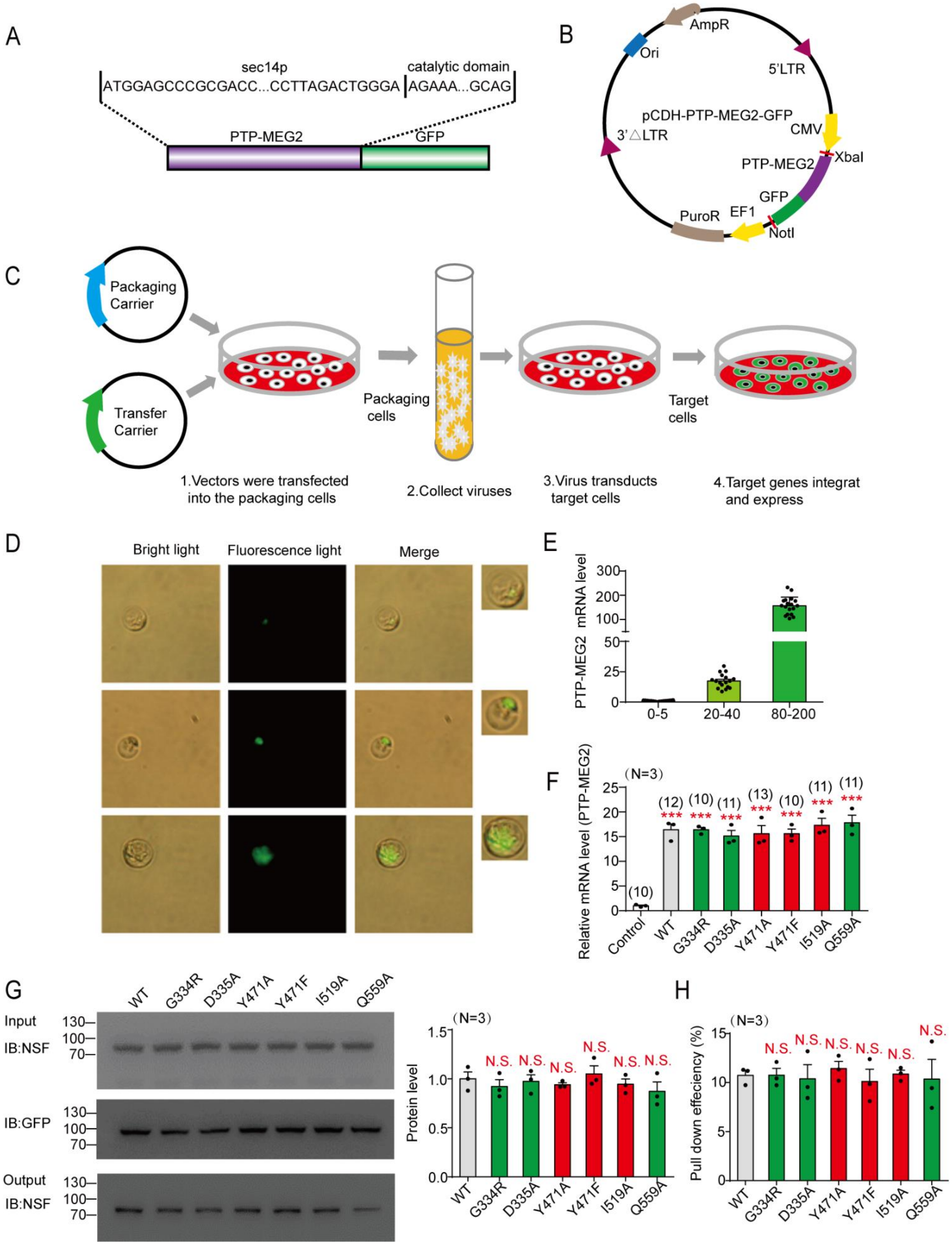

**Supplemental Figure 6. Effect of PTP-MEG2 mutations on catecholamine secretion from primary chromaffin cells.**

(A).The nucleotide sequence of the PTP-MEG2-GFP construct used for overexpressing PTP-MEG2 in primary mouse chromaffin cells.

(B). Schematic diagram of PTP-MEG2-GFP lentivirus vector.

(C).Schematic representation of lentivirus encompassing PTP-MEG2 wild type or mutants production, transduction and expression in mouse primary chromaffin cells.

(D).Primary mouse chromaffin cells were transduced with lentivirus containing empty vector, the vector encoding wild-type PTP-MEG2, or different mutants with a GFP tag at the C-terminus. Expression of PTP-MEG2 in cells was recorded with their corresponding fluorescence intensity. The intensity of light: 100ms. Resolution : 1920*1080.

(E). The histogram shows the relationship between quantified fluorescence intensity levels and mRNA levels of primary chromaffin cells transduced with lentivirus encoding PTP-MEG2 wild type.

(F). After virus infection, the selected cells with fluorescence values between 20 and 40 in Supplemental Figure 6 (D)-(E) were selected for mRNA quantification analysis. The histogram shows the quantified mRNA levels for each different PTP-MEG2 mutants. Each measurement was performed at least 3 times with indicated cell numbers (at the top of each column) selected for each group.

(G). PC12 cells were infected with lentiviruses encoding PTP-MEG2-$D^{470}$A-GFP of wild type and different mutants. The GFP-affinity-beads were used in GFP-pull down assay to examine the binding of PTP-MEG2 to endogenous NSF from PC12 cells.

(H). The pull-down efficiency of GST-MEG2-WT and different mutants on interacting NSF in Supplemental Figure 6G was calculated with NSF (amount of output)/ NSF (amount of input)*100%.

F, G, * indicates the transduced PTP-MEG2 group compared with the empty vector group. ***, P< 0.001 compared with the control groups. N.S. represented no difference compared with the control groups. All the data were analysed using one-way ANOVA.

### Supplemental figure 7

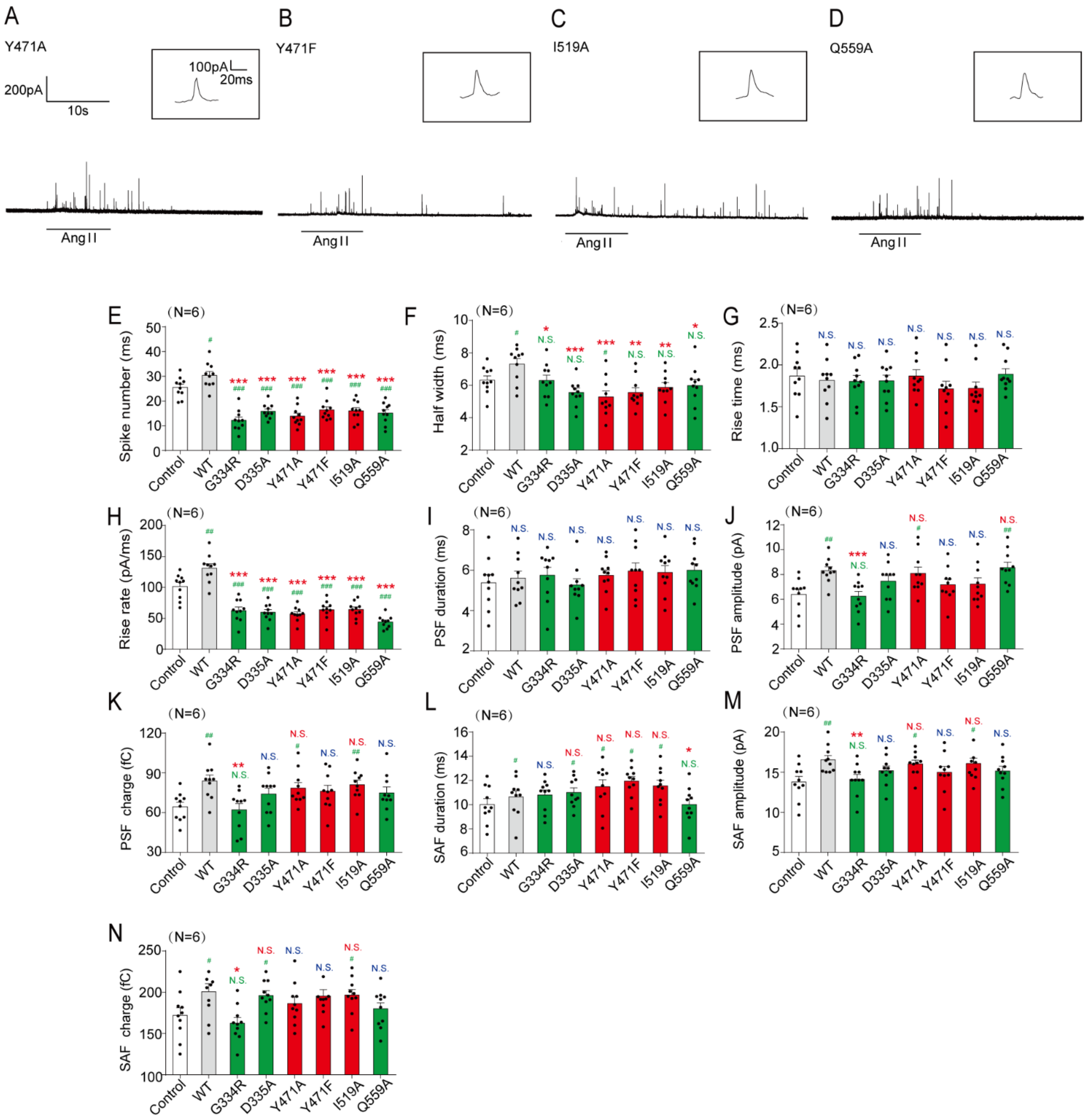

**Supplemental Figure 7. Effects of PTP-MEG2 mutations on catecholamine secretion from primary chromaffin cells.**

(A-D). Representative image of amperometric spikes of primary mouse chromaffin cells overexpressing PTP-MEG2-$Y^{471}$A (A), PTP-MEG2-$Y^{471}$F (B), PTP-MEG2-$I^{519}$A (C) and PTP-MEG2-$Q^{559}$A (D) in response to 100nM AngII stimulation.

(E-N). Quantitative analysis of the amperometric spikes of primary mouse chromaffin cells transduced with different PTP-MEG2 mutations in PTP-MEG2-phospho-NSF-$pY^{83}$ segment interface. Parameters of amperometric spikes included spike number (E), half width (F), rise time (G), rise rate (H), PSF duration (I), PSF amplitude (J), PSF charge (K), SAF duration (L), SAF amplitude (M) and SAF charge (N).

E-N, * indicates PTP-MEG2 mutants overexpression groups compared with the WT overexpression group. # indicates the overexpression group compared with the control group. *, P<0.05; **, P<0.01; *** P<0.001 and #, P<0.05; ##, P<0.01; ### P<0.001. N.S. represented no difference compared with the control groups. All the data were analysed using one-way ANOVA.

### Supplemental figure 8

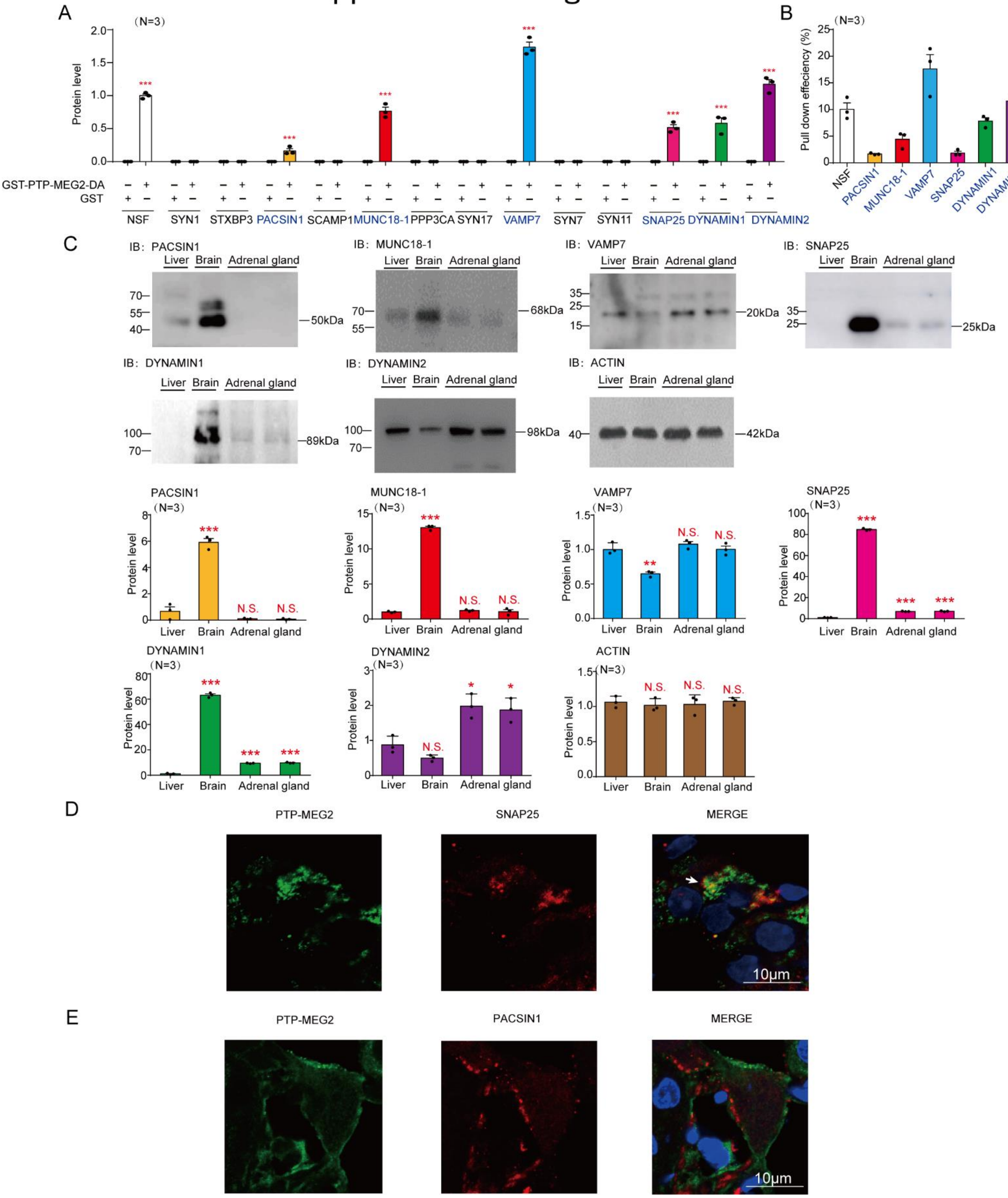

**Supplemental Figure 8. Screening and identification of candidate PTP-MEG2 substrates involved in PSF/SAF regulation.**

(A). Protein levels in Figure 6C were quantified from at least three independent experiments. * represented GST-PTP-MEG2-DA group compared with corresponding control GST group.

(B). The pull-down efficiency of GST-MEG2-DA in Figure 6C on NSF, PACSIN1, MUNC18-1, VAMP7, SNAP25, DYNAMIN1, and DYNAMIN2 was calculated with (protein amount of output)/ (protein amount of input)*100%.

(C). Upper panel: the expression of PACSIN1, MUNC18-1, VAMP7, SNAP25, DYNAMIN1, and DYNAMIN2 in mouse liver, brain and adrenal gland was detected with specific antibodies. Bottom panel: qualification of PACSIN1, MUNC18-1, VAMP7, SNAP25, DYNAMIN1 and DYNAMIN2 levels in different tissues. * indicates liver compared with the other tissues.

(D-E). Co-localization of PTP-MEG2 , SNAP25 (D), and PACSIN1 (E) in adrenal gland medulla was detected with immunofluorescence. White arrow stands for merged part. Rat adrenal medulla was incubated in 100 nM AngII for 1 minutes and fixed with formalin and sliced into 4-μm slides. Primary antibodies of PTP-MEG2 or candidate proteins (SNAP25, PACSIN1) were used to incubated with the slides overnight at 4℃. Secondary antibodies labeled with FITC (green) or TRITC (red) were applied for 1 hour at 25℃. Images were captured with confocal microscope. (A) (C) *, $P<0.05$; **, $P<0.01$; *** $P<0.001$. N.S. represented no difference compared with the control groups. All the data were analysed using one-way ANOVA.

### Supplemental figure 9

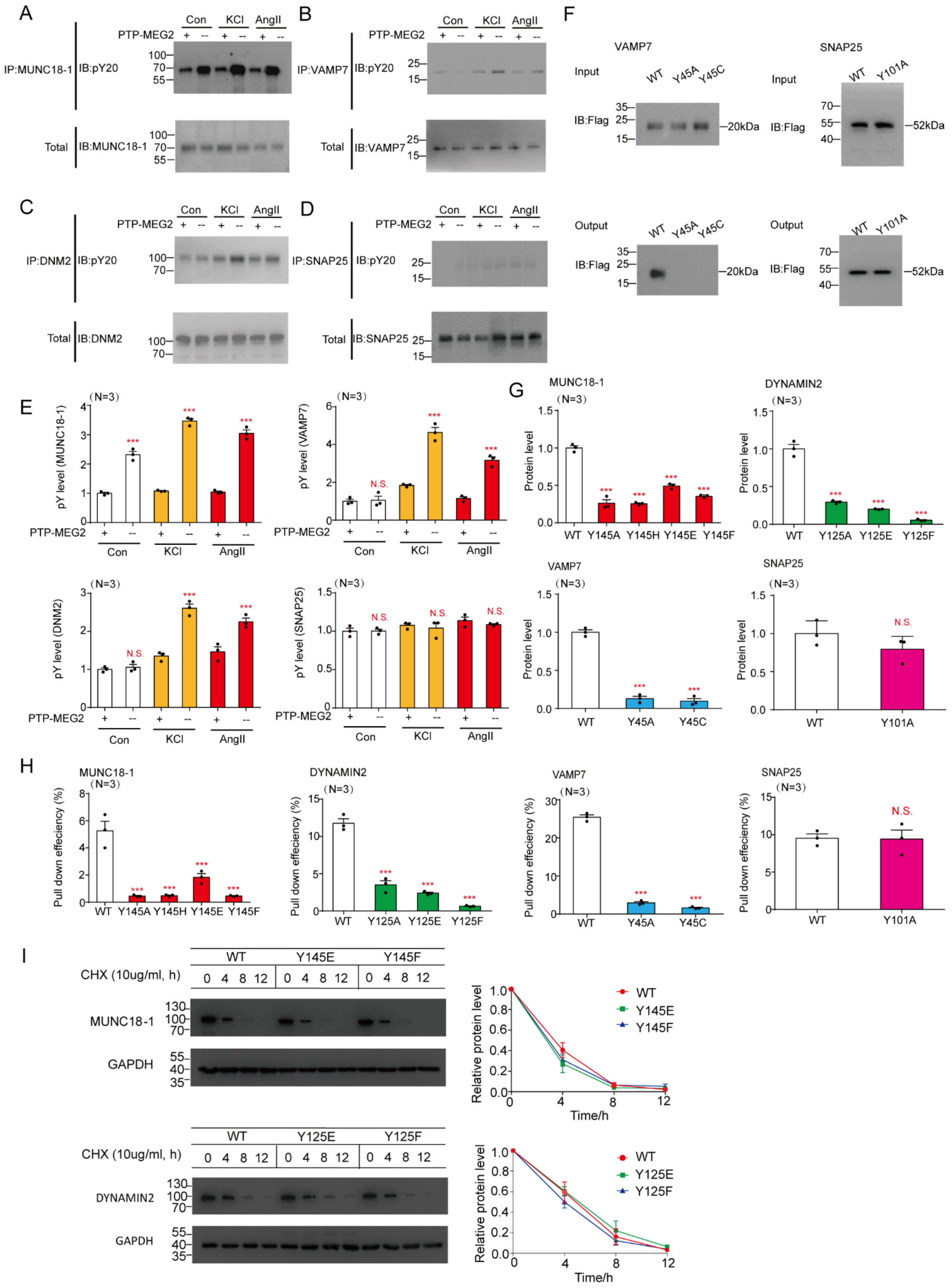

**Supplemental Figure 9. Influence of PTP-MEG2 on the phosphorylation of MUNC18-1, VAMP7, DYNAMIN2 and SNAP25.**

(A-D). Tyrosine phosphorylation levels of MUNC18-1 (A), VAMP7 (B), DYNAMIN2 (C) , and SNAP25 (D) in primary rat chromaffin cells were monitored. Rat adrenal medulla was lysed and incubated with Protein A/G beads after stimulation with 100nM AngII for 1 minutes, in presence or absence of 5μg PTP-MEG2 catalytic domain protein. Protein A/G beads were pre-incubated with primary antibody of MUNC18-1, VAMP7, DYNAMIN2, or SNAP25. Pan-phosphorylation antibody $pY^{20}$ was used to detect the phosphorylation of proteins binding with the complex of antibody-Protein A/G beads.

(E). The quantification of the tyrosine phosphorylation levels of MUNC18-1, VAMP7, DYNAMIN2, and SNAP25 recognized by $pY^{20}$ antibody. Quantification of total amount of MUNC18-1, VAMP7, DYNAMIN2, and SNAP25 binding with Protein A/G beads. *** represented P<0.001 and N.S. represented no difference compared with corresponding WT group.

(F). Interactions of the PTP-MEG2-trapping mutants with the VAMP7-$Y^{45}$A or $Y^{45}$C mutants (left), SNAP25-$Y^{101}$A (right). PC12 cells were transfected with the FLAG-tagged VAMP7 wild type or $Y^{45}$A or $Y^{45}$C mutants; FLAG-GFP-tagged SNAP25 wild type or $Y^{101}$A mutant 24 hours before stimulation with 100 nM AngII respectively. The cell lysates were then incubated with the GST-PTP-MEG2-$D^{470}$A for 4 hours under constant rotation. The potential PTP-MEG2 substrates were pulled down by GST-beads and their levels were examined by the FLAG antibody with western blotting.

(G). Protein levels in Figure 7 (D) and Supplemental Figure 9 (F) were quantified. A representative image of at least three independent experiments was shown.

(H). The pull-down efficiency of GST-MEG2-DA in Figure 7(D) and supplemental Figure 9 (F) on MUNC18-1 wide type and mutations, DYNAMIN2 wide type and mutations, VAMP7 wide type and mutations, SNAP25 wide type and mutations, was calculated with (protein amount of output)/(protein amount of input)*100%.

(I). HEK293 cells were seeded in 6-well plates. WT or different mutants' plasmids of MUNC18-1/DYNAMIN2 were transfected for 24 h. Cells were treated with cycloheximide (CHX) and harvested at different time points (0, 4, 8, 12h) for western blot analysis. The band intensity of MUNC18-1 or DYNAMIN2 was quantified (right panel). *** represented P<0.001 and N.S. represented no difference compared with corresponding control group. All the data were analysed using one-way ANOVA.

### Supplemental figure 10

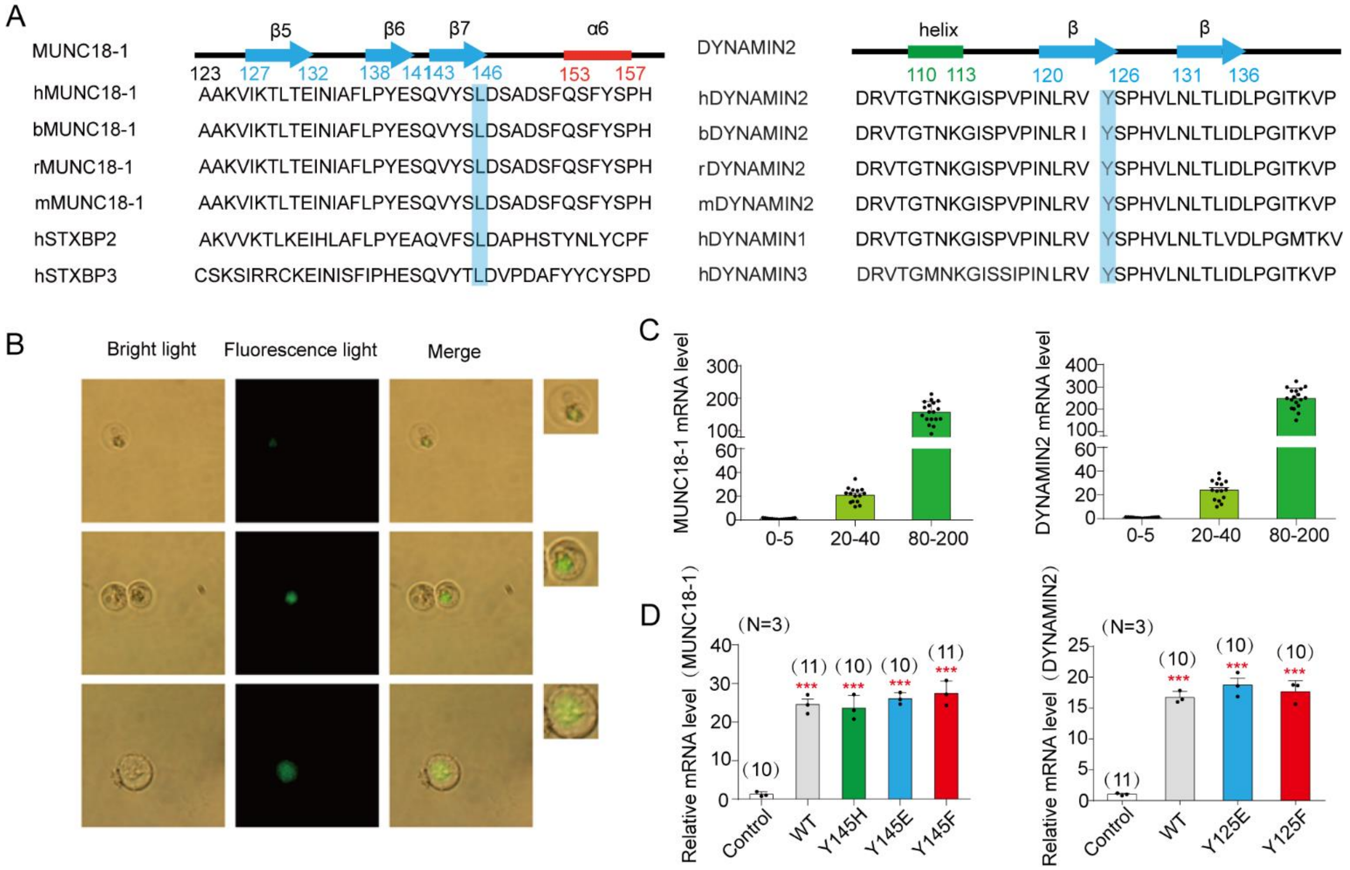

**Supplemental Figure 10. MUNC18-1-$Y^{145}$ and DYNAMIN2-$Y^{125}$ are essential in PTP-MEG2 regulation of PSF/SAF.**

(A). Sequence alignment of amino acids surrounding MUNC18-1-$Y^{145}$ and DYNAMIN2-$Y^{125}$. The MUNC18-1-$Y^{145}$ and DYNAMIN2-$Y^{125}$ were sites highlighted with blue.

(B).Primary mouse chromaffin cells were transduced with lentivirus containing empty vector, the vector encoding wild-type MUNC18-1, DYNAMIN2, or different mutants of these two proteins with a GFP-tag at their C-terminus. Expression of MUNC18-1 or DYNAMIN2 in cells was recorded with their fluorescence intensities. The intensity used for light: 100ms. Resolution : 1920*1080.

(C). The histogram shows the relationship between quantified fluorescence intensity levels and mRNA levels of MUNC18-1 and DYNAMIN2.

(D). After virus infection, cells with fluorescence values between 20 and 40 in Supplemental Figure 10 (B)-(C) were selected for mRNA quantification analysis. The histogram shows the quantified mRNA levels for each different MUNC18-1 (left panel) or DYNAMIN2 (right panel) mutants. Each measurement was performed at least 3 times with indicated cell numbers (at the top of each column) selected for each group.* indicates the transduced MUNC18-1 or DYNAMIN2 group compared with the empty vector group. *** represents $P<0.001$ compared with the control groups. All the data were analysed using one-way ANOVA.

### Supplemental figure 11

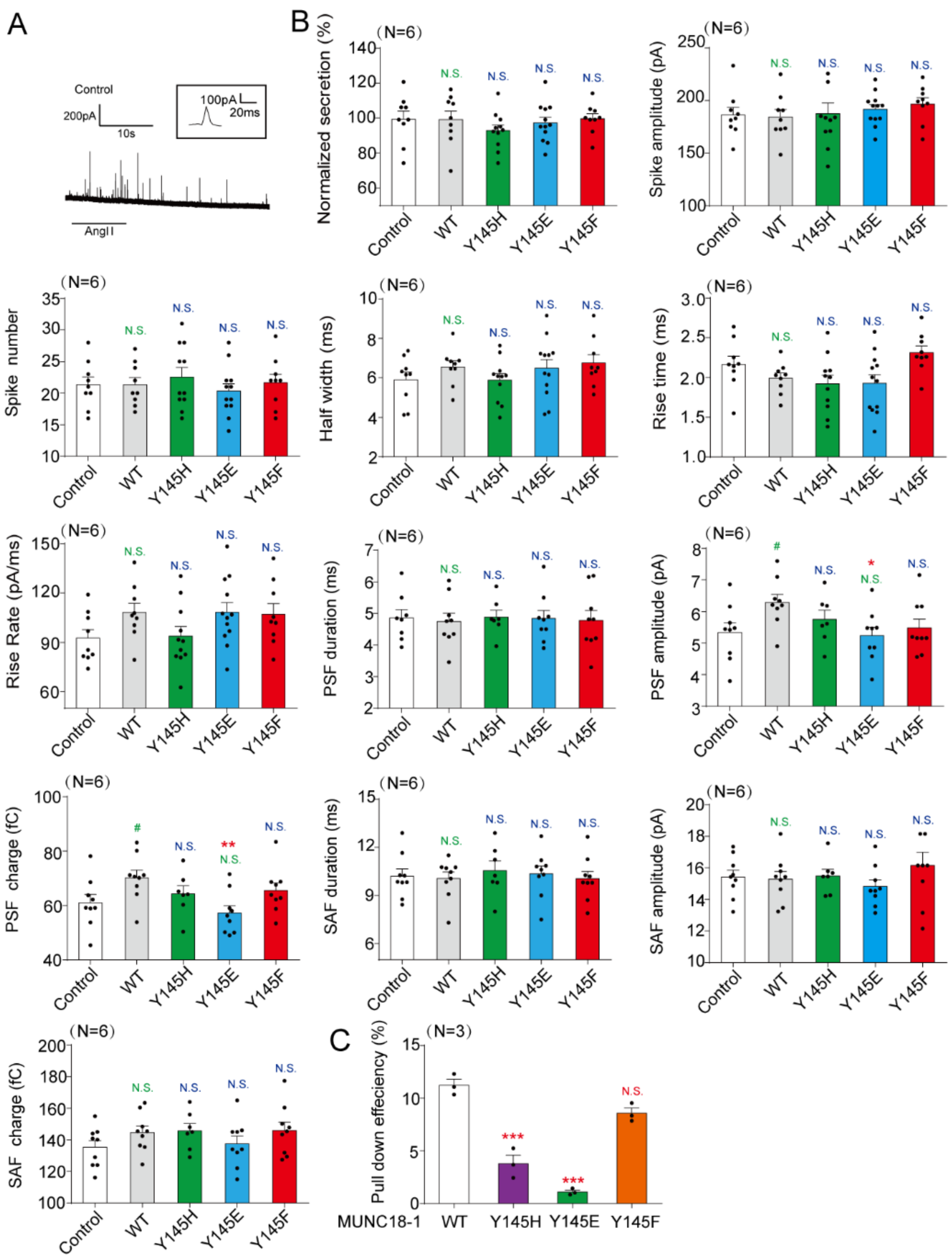

**Supplemental Figure 11. MUNC18-1-Y$^{145}$ is essential in PTP-MEG2 regulation of PSF/SAF.**

(A). Primary chromaffin cells were transduced with lentivirus containing empty vector. These cells were stimulated with 100nM AngII. The amperometric spikes were detected with CFE. Typical amperometric traces are shown.

(B). Parameters of amperometric spikes of primary chromaffin cells overexpressing MUNC18-1-Y$^{145}$H, Y$^{145}$E or Y$^{145}$F mutants, MUNC18-1 wild type, or empty vector. Primary chromaffin cells were transfected with empty vector, MUNC18-1-WT, Y$^{145}$H, Y$^{145}$E, or Y$^{145}$F by lentivirus and stimulated with AngII (100nM). The amperometric spikes including the secretion amount, spike amplitude, spike number, half width, rise time, rise rate, parameters of PSF and SAF were compared between empty vector, wild type, and mutation.

(C). Pull down efficiency in Figure 7H was evaluated by calculating SYNTAXIN1 output/input.

* indicates MUNC18-1 mutants overexpression groups compared with the WT overexpression group. # indicates MUNC18-1 overexpression group compared with the control group. *, $P<0.05$; **, $P<0.01$; ***, $P<0.001$ and #, $P<0.05$. N.S. represented no difference compared with corresponding control group. All the data were analysed using one-way ANOVA.

### Supplemental figure 12

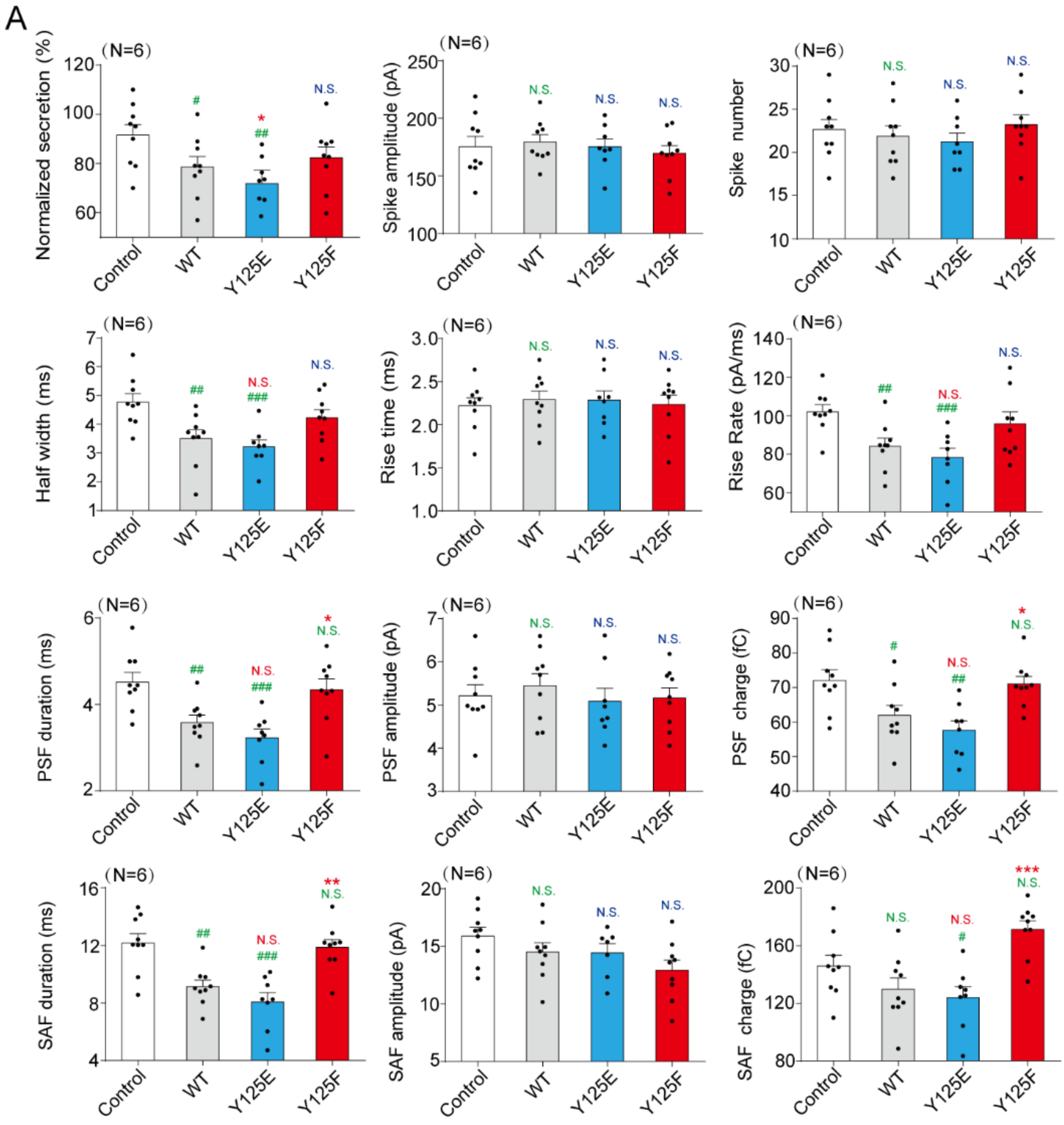

**Supplemental Figure 12. DYNAMIN2-$Y^{125}$ is essential in PTP-MEG2 regulation of PSF/SAF.**

(A). Parameters of amperometric spikes of primary chromaffin cells overexpressing DYNAMIN2-$Y^{125}$E or $Y^{125}$F mutants, DYNAMIN2 wild type, or empty vector. Primary chromaffin cells were transducted with empty vector, DYNAMIN2-WT, $Y^{125}$E, or $Y^{125}$F by lentivirus and stimulated with AngII (100nM). The amperometric spikes including the secretion amount, spike amplitude, spike number, half width, rise time, rise rate, parameters of PSF and SAF were compared between empty vector, wild type, and mutants. * indicates DYNAMIN2 mutants overexpression compared with the WT overexpression group. # indicates DYNAMIN2 overexpression group compared with the control group. *, $P<0.05$; **, $P<0.01$; ***, $P<0.001$ and #, $P<0.05$; ##, $P<0.01$; ###, $P<0.001$. N.S. represented no difference compared with corresponding control group. All statistical significance was calculated with one-way ANOVA.

### Supplemental figure 13

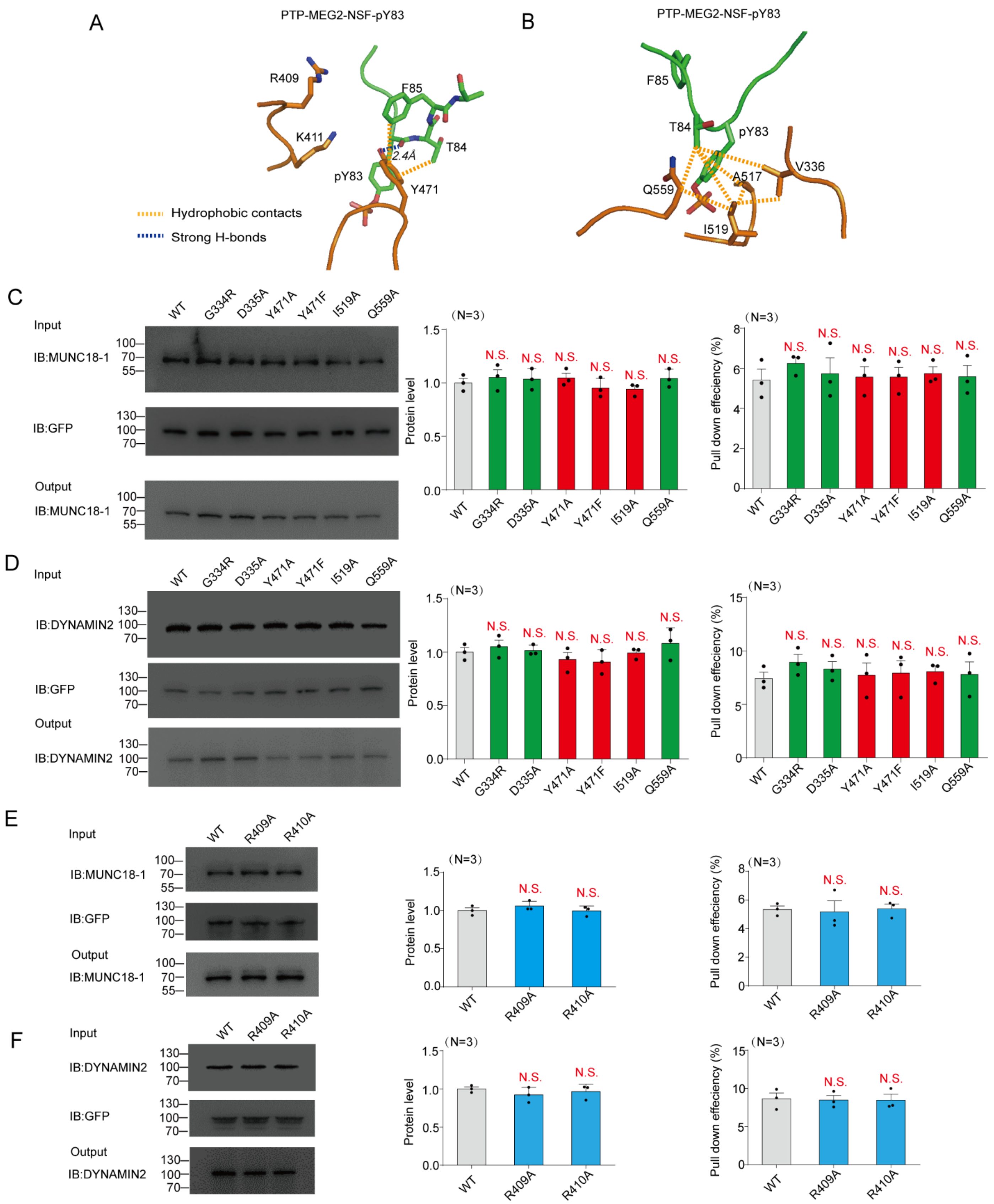

**Supplemental Figure 13. The structural comparison of PTP-MEG2 in complex with phospho-segment derived from different substrates and interaction of PTP-MEG2-WT and different mutants toward MUNC18-1 and DYNAMIN2.**

(A-B). The detailed interactions of the PTP-MEG2 $Y^{471}$ (A) or $I^{519}$ (B) and the NSF-pY83 phospho-segment.

(C-F). PC12 cells were infected with lentiviruses encoding PTP-MEG2-$D^{470}$A-GFP of wild type and different mutant types. The GFP-beads were used in GFP-pull down assays to examine the binding of different PTP-MEG2 mutants to endogenous MUNC18-1 and DYNAMIN2. Protein levels and pull-down efficiency were quantified. A representative image of at least three independent experiments was shown. N.S. represented no difference compared with corresponding WT group. All statistical significance was calculated with one-way ANOVA.

### Supplemental figure 14

A MUNC18-1

| Enzymes | $k_{cat}/K_m$ [ $10^5$, $M^{-1}s^{-1}$ ] | Fold of Activity Decrease | Selectivity for Peptide |
|---|---|---|---|
| WT | 146±12 | 1 | 1 |
| Y333A | 6.81±0.25 | 21.6 | 2.12 |
| G334R | 18.9±1.6 | 7.75 | 8.81 |
| D335A | 3.19±0.2 | 46 | 51.28 |
| R409A | 27.55±0.62 | 5.33 | 6.56 |
| R410A | 10.97±0.47 | 13.39 | 5.70 |
| Y471A | 108±4.6 | 1.35 | 1.21 |
| Y471F | 115±4.7 | 1.28 | 1.05 |
| I519A | 187±14 | 0.78 | 0.22 |
| Q559A | 17.5±0.67 | 8.41 | 6.21 |

DYNAMIN2

| Enzymes | $k_{cat}/K_m$ [ $10^5$, $M^{-1}s^{-1}$ ] | Fold of Activity Decrease | Selectivity for Peptide |
|---|---|---|---|
| WT | 106±8.3 | 1 | 1 |
| Y333A | 23.4±1.25 | 4.54 | 0.45 |
| G334R | 24.8±1.2 | 4.29 | 4.88 |
| D335A | 7.02±0.3 | 15.2 | 16.9 |
| R409A | 24.12±0.57 | 4.41 | 5.43 |
| R410A | 9.98±0.62 | 10.67 | 4.55 |
| Y471A | 97.9±3.3 | 1.08 | 0.98 |
| Y471F | 61.9±2.7 | 1.72 | 1.41 |
| I519A | 31.9±4.2 | 3.33 | 0.93 |
| Q559A | 8.93±0.32 | 11.9 | 8.78 |

**Supplemental Figure 14. Catalytic activity of PTP-MEG2-WT and different mutants toward $pY^{145}$-MUNC18-1 and $pY^{125}$-DYNAMIN2.**

(A).$k_{cat}/K_m$ of PTP-MEG2-WT towards MUNC18-1-$pY^{145}$ phospho-segment and DYNAMIN2-$pY^{125}$ phospho-segment were determined and the ratios of this activity compared to $k_{cat}/K_m$ of PTP-MEG2-WT towards the small artificial substrate pNPP were calculated. The assays were performed with DMG buffer: 50 mM 3,3-dimethyl glutarate pH 7.0, 1 mM EDTA, 1 mM DTT. The ionic strength was maintained at 0.15 M (adjusted by NaCl). The enzymatic reactions were carried out at room temperature.

### Supplemental figure 15

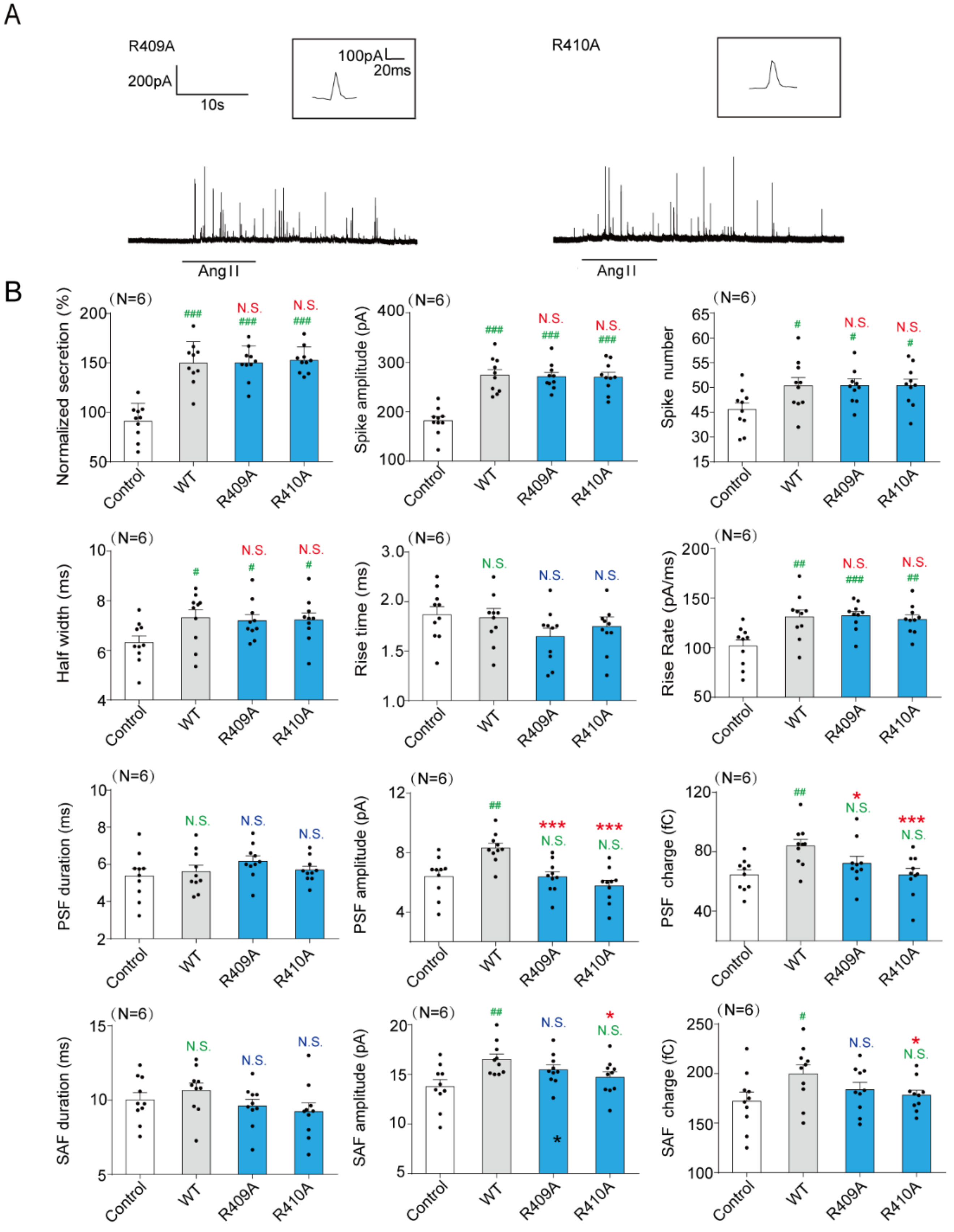

**Supplemental Figure 15. Effect of PTP-MEG2-$R^{409}A$ or $R^{410}A$ mutations on catecholamine secretion from primary chromaffin cells.**

(A). Primary chromaffin cells were transduced with lentivirus containing PTP-MEG2-$R^{409}A/R^{410}A$. These cells were stimulated with 100nM AngII. The amperometric spikes were detected with CFE. Typical amperometric traces are shown.

(B). Parameters of amperometric spikes of primary chromaffin cells overexpressing PTP-MEG2-$R^{409}A$ or $R^{410}A$ mutants. Primary chromaffin cells were transfected with PTP-MEG2-$R^{409}A$ or $R^{410}A$ mutants by lentivirus and stimulated with AngII (100nM). The amperometric spikes including the secretion amount, spike amplitude, spike number, half width, rise time, rise rate, parameters of PSF and SAF were compared between empty vector, wild type, and mutation. * indicates MUNC18-1 mutants overexpression groups compared with the WT overexpression group. # indicates MUNC18-1 overexpression group compared with the control group. *, P<0.05; ***, P<0.001 and #, P<0.05; ##, P<0.01; ### P<0.001. N.S. represented no difference compared with corresponding control group. All the data were analysed using one-way ANOVA.
